# Conspecific sociability is regulated by associative learning circuits

**DOI:** 10.1101/2024.11.25.624845

**Authors:** Victor Lobato-Rios, Thomas Ka Chung Lam, Pavan Ramdya

## Abstract

Sociability, an animal’s ease and propensity to interact with members of its own species, is a prerequisite for many important social interactions including courtship, mating, brood rearing, and collective behavior. Despite its fundamental nature, the manner through which sociability arises—whether it is innate or acquired and its neural mechanisms— remains unknown. Here, we found that the fly, *Drosophila melanogaster*, produces a constellation of fearful reactions when it encounters another fly for the first time. However, these animals become sociable following several hours of exposure to conspecific (but not heterospecific) odors. Two large-scale neural silencing screens of 188 brain cell types revealed that both initial fearful reactions and learned sociability depend upon overlapping networks including circuits in the mushroom body, the principal center for associative learning and memory in the insect brain. Functional recordings of key mushroom body output neurons (MBONs) from this network over two hours of social interactions support a mechanistic model whereby fly odors modulate social valence by rebalancing MBON population activity, biasing action selection, and thereby driving sociable rather than fearful responses towards other flies. Thus, a center for learning and memory plays a fundamental role in establishing the basis for most social interactions.

## Introduction

In the natural world, animals live in a diverse milieu among numerous other species. Therefore, they must constantly decide who to avoid, ignore, or interact with. The propensity for animals to seek and interact with conspecifics (members of their own species) is a trait known as sociability ^1,2^. Being sociable forms the foundation of numerous ethologically important social behaviors including courtship ^3^, caring for the young, hunting in packs ^4^, making collective decisions ^5^, and working as a colony ^6^. These group-level interactions have been shown to reduce the risk of predation, improve foraging success, increase mating rates, and facilitate the transmission of information ^4,7,8^. By contrast, isolation and the absence of social integration can increase anxiety and stress ^9^ and reduce an animal’s life span ^10^.

These observations raise the question of how animals become sociable in the first place. This tendency to welcome interactions with conspecifics may be innate, like the urge to follow the scent of a potential mate ^11^ or to escape a looming threat ^12,13^. Or it may be modulated by experience ^14^ and internal state ^15^. Numerous findings suggest that sociability is a dynamic rather than a fixed trait. First, environmental factors and molecules, including neuropeptides, have been shown to regulate sociability across diverse species including insects and humans ^16,17,18,19,20^. Second, early life experiences like social isolation can make individuals more ^21,6^ or less ^22,23^ willing to interact with conspecifics. Nevertheless, the neural circuit mechanisms regulating this dynamism remain unknown. On one hand, sociability might arise through the relatively rapid habituation of peripheral sensory pathways tuned to conspecific odors and visual cues. On the other hand, sociable behaviors may be modulated by central circuits for long-term memory formation and maintenance.

To distinguish between these and other potential models, we studied sociability in the adult fly, *Drosophila melanogaster*. Flies are not considered a eusocial species as they do not form structured societies and colonies like other insects. However, flies aggregate in both natural ^24,25^ and laboratory environments even in the absence of an attractive food source ^26,27^. Additionally, they depend on conspecific aggregation for social behaviors like courtship and mating ^11^, aggressive competition ^28,29^, and collective action ^8,5^. In *D. mel* the study of social behavior is facilitated by a numerically compact and stereotyped nervous system, genetic tools for perturbing and recording identified neurons, and a complete map of the adult brain ^30,31,32,33,34^. This has allowed the neural mechanisms for courtship ^35,11^ and aggression ^36,28^ to be deeply investigated, demonstrating that social interactions are multimodal: they involve the integration of olfaction and taste ^37,38^, vision ^39^, and audition ^40^. Still, the mechanisms giving rise to conspecific sociability—the basis for these and other social interactions—remain unclear. For instance to what extent does it depend on exposure to other flies? It is known that individual ^41^ and group behaviors can be influenced by social experience: although flies tend to cluster, this trait can be disrupted by social isolation ^23,27^ and can recover following conspecific exposure ^42^. Furthermore, prior social experience can modulate courtship behaviors in a manner that resembles associative learning ^43,44^.

Here we dissect the mechanisms of *Drosophila* sociability. We found that, flies exhibit a constellation of fearful reactions when they encounter another fly for the first time. These fearful reactions are lost over several hours of exposure to conspecifics. Furthermore, multiple days of social isolation causes group-raised animals to revert to being less sociable. Thus, sociability is learned and can be forgotten. We found that this sociability learning arises from exposure to conspecific odors, but not to odors from other species. Visual, gustatory, and tactile cues are not required. Two neural silencing screens of 188 brain cell types show that both fearful reactions and learned sociability depend upon overlapping and interconnected networks in the mushroom body, the principal brain region implicated in insect learning and memory. By recording the activities of key mushroom body output neurons (MBONs) in this network, we found that their dynamics initially correlate with behavioral vigor. However, the baseline and locomotor-related activity of MBON-β’2mp, neurons previously implicated in aversive learning, greatly diminishes over two hours of conspecific exposure. These data support a model in which fly odors modulate social valence through long-term learning rather than habituation at the sensory periphery by rebalancing MBON population activity, biasing action selection, and thereby driving sociable rather than fearful responses towards other flies. Thus, sociability is a dynamic trait that is regulated by circuits in the brain’s center for learning and memory.

## Results

### Sociability is learned and can be forgotten

An adult fly can emerge from its pupal case in very different social environments. On one hand, it might be surrounded by a thriving community of other flies, as on a mass of rotting fruit. On the other hand, it might eclose alone, surrounded only by other species. In the first case a fly might benefit from immediately engaging in social exchanges like courtship, whereas in the second case it might be better served to remaining cautious until it is certain that these exchanges might be reciprocated appropriately. These two scenarios raise the fundamental question of how flies react when they see another fly for the first time. Are they hard-wired to immediately initiate social interactions or do they need time to gradually learn to become sociable?

To address this question, we compared interactions between flies that were raised in isolation as pupae to 5–8 days post eclosion (‘single-housed’) to a control group that were housed together for an equivalent period of time (‘group-housed’) **(Fig. 1a)**. We placed pairs of flies raised under the same conditions in an arena **(Extended Data Fig. 1a–b)**, recorded their behavior for 10 minutes, and tracked their movements using deep network-based 2D pose estimation ^45^ **(Fig. 1b–c)**. Strikingly, we found that animals raised in isolation avoid one another and generate a variety of fearful behaviors including careful monitoring of the other fly’s movements, rapidly retreating and running away, or freezing in place **(Fig. 1b; Supplementary Video 1)**. By contrast, group-housed animals were sociable: they did not exhibit fearful reactions but instead came into close proximity with the other fly to walk in tandem or even to interact with their legs **(Fig. 1c; Supplementary Video 2)**.

**Fig. 1:**
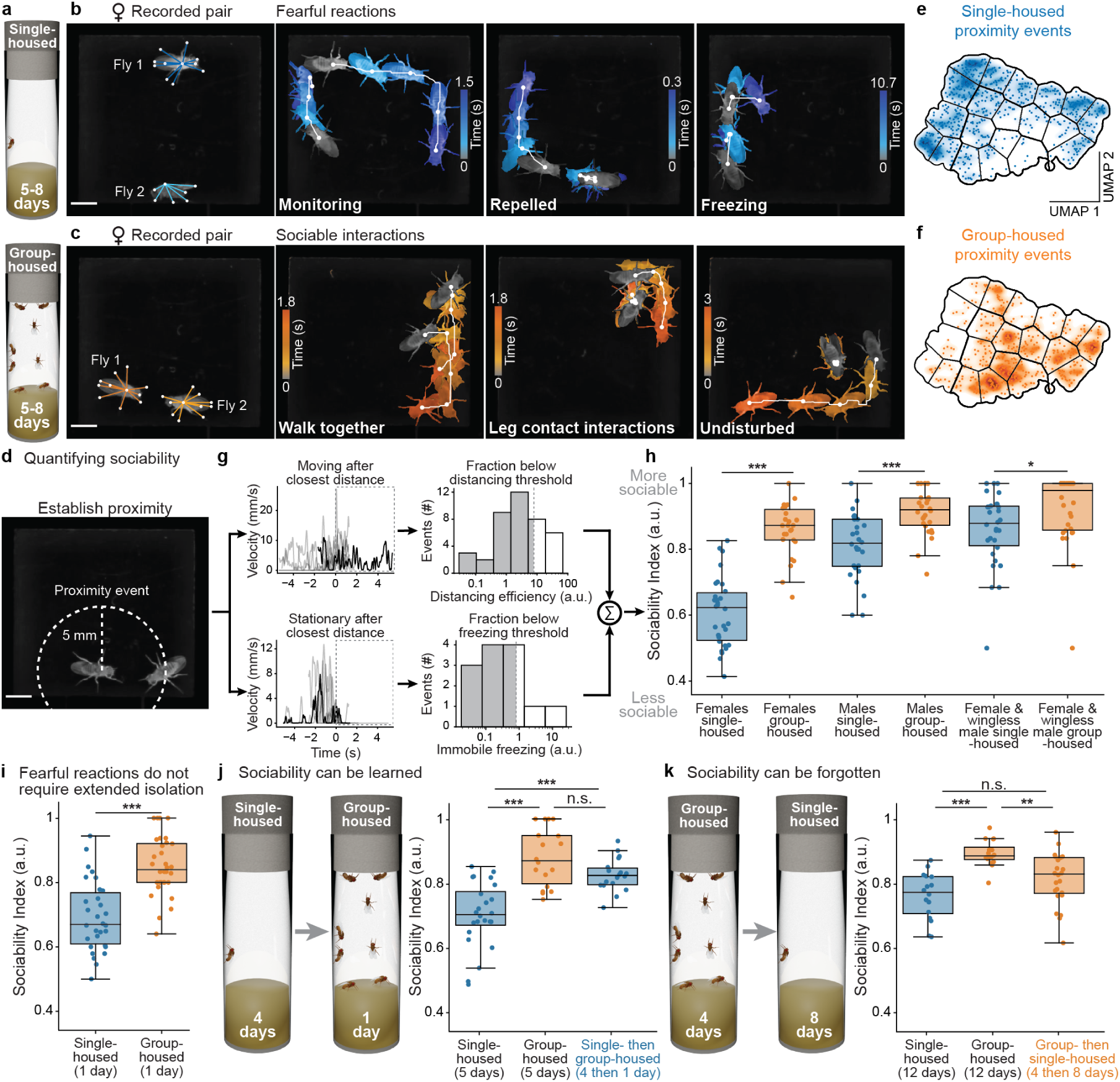
Sociability is learned and can be forgotten. **(a)** From prior to eclosion as adults, flies are raised either single-housed (top) or group-housed (bottom). **(b)** Single-housed flies (females in this example) are paired within an arena in which their behavior is recorded and tracked while maintaining their identities. They show fearful reactions including careful monitoring of, repulsion from, or freezing in reaction to the other fly **(Supplementary Video 1)**. Time across video frames is represented by a gray-blue color gradient that is evenly spaced between the start and end of reactions. Overlaid are the real trajectories of flies over time (white lines). Scale bar is 2 mm in panels **b-d**. **(c)** Group-housed flies are paired within an arena in which their behavior is recorded and tracked while maintaining their identities. They show sociable behaviors including walking amongst one another, contacting each other with their legs, and being undisturbed **(Supplementary Video 2)**. Time across video frames is represented by a gray-orange color gradient that is evenly spaced between the start and end of behaviors. Overlaid are the real trajectories of flies over time (white lines). **(d)** Proximity events are defined as when flies are within 5 mm of one another. **(e–f)** Proximity events in **(e)** single (blue) or **(f)** group-housed (orange) animals, visualized in a low-dimensional embedding of behavioral space generated by UMAP. Each dot represents one proximity event. Overlaid are kernel density estimates to highlight distributions. Regions separated by lines are 20 clusters identified using *k*-means clustering. **(g)** To calculate the sociability index, proximity events are obtained from each pairing experiment and separated into two categories as a function of the fly’s reaction after coming into closest proximity. For flies that move, distancing efficiency is quantified. For flies that do not move, immobile freezing is quantified. Distancing and freezing thresholds are determined by control group data for each experiment. **(h)** The sociability index represents an individual’s proportion of undisturbed interactions. More sociable animals have higher values. Shown are the sociability indices for female pairings (*n* = 30 single-housed and *n* = 26 group-housed animals), male pairings (*n* = 28 single-housed and *n* = 30 group-housed animals), and mixed female-male pairings (*n* = 30 animals). Males are wingless in this last pairing to prevent courtship songs and wing displays that modulate female locomotion. Group-housed female data are used to determine index thresholds. Mann–Whitney statistical tests were computed for female-female, male-male, and female-male experiments separately. **(i)** To test if fearful reactions already occur shortly after eclosion, we compared the sociability of single (*n* = 30) or group-housed (*n* = 32) flies 1 day post eclosion. **(j)** To test if sociability could be learned, flies were single-housed then group-housed. Shown are sociability indices for single-housed flies (*n* = 24), group-housed flies (*n* = 18), and flies single-housed and then group-housed for 1 day (*n* = 18). **(k)** To test if sociability could be forgotten, flies were group-housed and then isolated. Shown are sociability indices for single-housed flies (*n* = 16), group-housed flies (*n* = 16), and flies group-housed and then single-housed for eight days (*n* = 24). For box plots, the box limits indicate the central 50% of the data and the line inside the box depicts the median value. The whiskers extend out of the box to the full range of the remaining data. Data points placed beyond the whiskers are outliers. For panels **i–k**, we used a Kruskal–Wallis test followed by a post hoc Conover’s test with a Holm correction for multiple comparisons. *** *P <* 0.0001, ** *P <* 0.001, * *P <* 0.01, n.s. *P* ≥ 0.01.

To compare these distinct reactions of single- and group-housed animals in an unbiased manner, we analyzed behavioral tracking data using unsupervised clustering. We identified ‘proximity events’ as periods when each flies were close (*<*5 mm) to one another for more than 0.5 s **(Fig. 1d)**. Then we measured the relative positions, velocities, and orientations of flies during each proximity event. We performed UMAP ^46,47^, an unsupervised non-linear dimensionality reduction technique, to produce a 2D embedding of these data and then applied *k*-means clustering to the resulting 2D points. These plots confirmed that single- and group-housed animals generate a distinct suite of behaviors whose data were enriched in separate regions of the embedding space representing predominantly avoidance and undisturbed behaviors, respectively **(Fig. 1e–f; Extended Data Fig. 2a–d; Supplementary Video 3)**.

To have a second, more interpretable means of comparing these differences in social interactions of single- and group-housed animals, we devised a ‘sociability index’ ranging from 0 (fearful and not sociable) to 1 (highly sociable). We split proximity events into two classes based on whether, in response to their neighbors, flies (i) move away, as during monitoring or retreat, or (ii) stay in place, as during freezing **(Fig. 1g)**. In the first case we quantified a ‘distancing efficiency’ for which a low value (i.e., not running away) indicates higher sociability **(Supplementary Video 4)**. For stationary flies, we quantified ‘immobile freezing’ where a low value (e.g., the continuation of leg movements for grooming or postural adjustments) also indicates higher sociability **(Supplementary Video 5)**. Finally, to obtain a ‘sociability index’, we summed the fraction of distancing efficiency and immobile freezing events below a threshold defined by the distribution of these metrics in control, group-housed animals (see Methods). Importantly, proximity events above and below these thresholds were also located in regions of our 2D behavioral embedding associated with single and group-housed animals, respectively **(Extended Data Fig. 2e)**. Thus, a high sociability index value indicates that, upon close proximity, flies do not move apart quickly but rather either groom or interact with one another. On the other hand, a low sociability index value indicates that flies run away or freeze in response to their neighbors.

We found that this index effectively summarizes and distinguishes between fearful reactions of single-housed and undisturbed reactions of group raised animals **(Fig. 1h)**. Importantly, this distinction is not strongly sensitive to the thresholds chosen **(Extended Data Fig. 3)** and is not strongly correlated with the general activity level of flies: more active flies do not trivially have a higher distancing efficiency causing them to be considered less sociable **(Extended Data Fig. 4)**.

To test how well these behaviors generalize across sexes, we performed the same experiments with pairs of males. There we observed the same, although slightly less pronounced differences in behavior between single- and group-housed males **(Fig. 1h, middle)** compared to females **(Fig. 1h, left)**. We also examined sociability among mixed pairs of female and male flies. To prohibit the production of courtship songs (which cause females to halt their locomotion ^48,49^) we clipped the wings of each male. Despite the fact that males raised in isolation have been shown to more aggressively court and mate ^3^, we still observed significantly higher repulsion between single-housed females and males **(Fig. 1h, right)**. Thus, all pairings of sexes are more sociable when raised group-rather than single-housed. Based on these findings, and to avoid known male-male aggression behaviors ^28^, hereon we focused on sociability between pairs of female flies.

We next wondered to what extent the innate, default state of a newly eclosed fly is to be cautious and fearful towards conspecifics, or if this fear arises from the absence of social exposure. To address this question, we examined animals that were single-housed for only one day following eclosion. We found that these animals were already significantly less sociable than age-matched group-housed controls **(Fig. 1i)**. These data suggest that fear towards conspecifics is the initial state of a newly eclosed adult fly. Fully establishing this would require experiments shortly after eclosion, but such experiments would be confounded by a variety of hormone-driven maturation events during this period including wing inflation and cuticle tanning ^50^.

The fact that sociability only appears following exposure to conspecifics (i.e., group housing) suggests that it is learned rather than hard-wired. To test this, we first asked if the fearful reactions of single-housed animals can be transformed into sociable behaviors by exposure to conspecifics. We placed animals that had previously been isolated for four days into group housing for one additional day. Remarkably, after only one day of group housing their behaviors more closely resembled those of animals raised for five days in a group **(Fig. 1j)**. A second hallmark of learning is the capacity to forget. Therefore, in a second experiment we asked if isolating group raised animals reduces their sociability. Indeed, we found that eight days of isolation causes group-housed animals to behave less sociably towards conspecifics **(Fig. 1k; Extended Data Fig. 5)**. Notably, this effect was not as profound with fewer days of isolation, suggesting that forgetting to be sociable occurs more slowly than learning. Taken together these results show that conspecific sociability is a trait that must be learned through social exposure and can be forgotten in its absence.

### Conspecific odors drive sociability learning

We hypothesized that this learning process entails biasing action selection circuits from driving fearful reactions towards conspecifics to instead generating unperturbed behaviors. This transition might arise from exposure to a variety of sensory cues including the other animal’s visual appearance, odor profile (including pheromones), or gustatory and mechanical cues detected through tactile interactions. To identify which of those are important we first defined the time scale over which animals become sociable. By continuously monitoring and quantifying the sociability of pairs of single- or group-housed flies across two hours of recording, we found that single-housed animals show a gradual increase in sociability over time. By contrast, group-housed flies had no change in sociability over the same period. After two hours, the sociability indices of the two groups were similar, although still statistically different **(Fig. 2a)**.

**Fig. 2:**
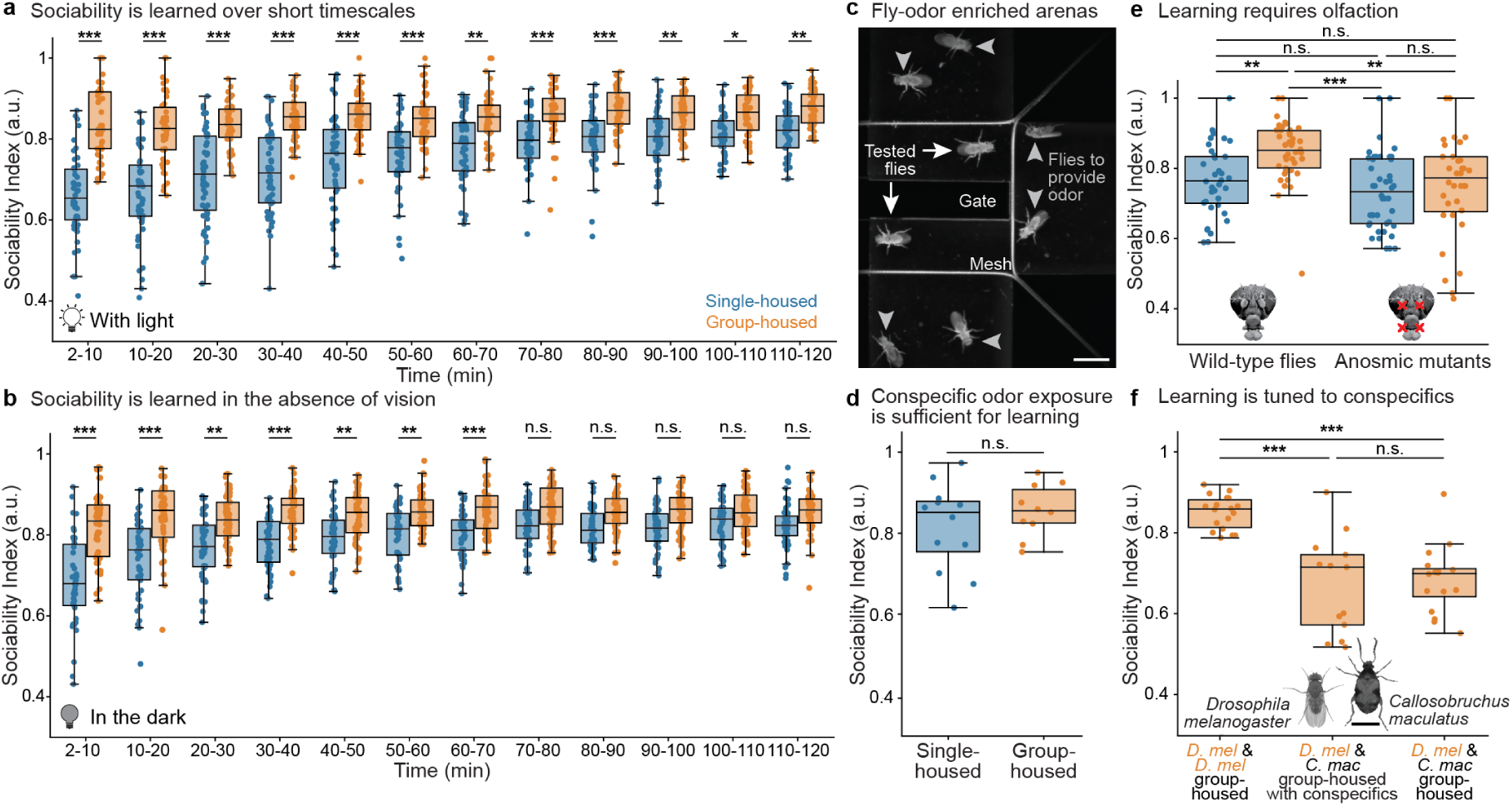
Sociability learning is regulated by conspecific odors. **(a)** To examine the temporal dynamics of sociability learning, flies were paired and recorded for two hours in an illuminated arena. Shown are sociability indices for single (*n* = 48) and group-housed (*n* = 47) animals for consecutive time bins of 10 min each. For the first time bin the first 2 min were not analyzed because during this period animals tend to explore the novel arena environment. **(b)** To test the role of vision in sociability learning, flies were paired and recorded for two hours in the dark. Shown are sociability indices for single (*n* = 46) and group-housed (*n* = 54) animals for consecutive time bins of 10 min each. **(c–d)** The relative roles of smell, touch, and taste in sociability learning were assessed by placing flies in **(c)** a fly odor-enriched arena in which a mesh separates experimental animals from odor-conferring animals to prevent physical contact (i.e., touch and taste). A movable gate that separates the pair of experimental flies is opened after 2 h to assess the impact of odor exposure on sociability learning. Scale bar is 3 mm. **(d)** Shown are sociability indices for single (*n* = 12) or group-housed (*n* = 10) flies after spending 2 h in odor-enriched arenas without interacting with another fly. A Mann–Whitney statistical test was performed. n.s.: *P* ≥ 0.01. **(e)** To assess the necessity of olfaction in sociability learning, anosmic mutants were tested. Shown are sociability indices for single and group-housed wild-type (*n* = 36 and 42, respectively) and anosmic mutant (*n* = 36 and 44, respectively) animals. **(f)** To explore the specificity of sociability learning to conspecific odors, the cowpea beetle, *C. maculatus*, was paired with flies, *D. mel.*, in behavioral arenas. Shown are sociability indices for *D. mel* when they are (left) group-housed with flies and paired with another fly (*n* = 18), (middle) group-housed with flies and paired with a beetle that was raised with other beetles (*n* = 13), and (right) group-housed with flies and beetles and paired with a beetle from the same mixed group (*n* = 16). Photos of female *D. mel.* and *C. maculatus* are shown to enable a comparison of their sizes. Scale bar is 2 mm. For box plots, the box limits indicate the central 50% of the data and the line inside the box depicts the median value. The whiskers extend out of the box to the full range of the remaining data. Data points placed beyond the whiskers are outliers. For panels **a–b; e–f**, a Kruskal–Wallis test followed by a post hoc Conover’s test with a Holm correction for multiple comparisons was used. *** *P <* 0.0001, ** *P <* 0.001, * *P <* 0.01, n.s. *P* ≥ 0.01.

Then we performed experiments in which we systematically suppressed individual sensory cues and modalities during sociability learning. First, to test the role of vision, we measured sociability learning over two hours as before but in the dark. Flies in well-lit environments react to one another from a distance. By contrast, we found that flies in the dark do not react fearfully towards one another until they come close enough to touch one another **(Extended Data Fig. 6; Supplementary Video 6)**. Nevertheless, single-housed flies in the dark still became increasingly sociable over time: they were nearly indistinguishable from group-housed animals after ∼70 minutes **(Fig. 2b)**. Thus, vision is important for driving fearful reactions but not required for sociability learning.

Next, we tested the potential contributions of olfaction, touch and taste. We designed a ‘fly-odor enriched’ arena in which six flies (conspecific odor sources) were placed behind a mesh perimeter, making them inaccessible for tactile and taste interactions using leg mechanosensors ^51^ and gustatory sensors ^52^ **(Fig. 2c; Extended Data Fig. 1c–d)**. At the center of this arena, a pair of single-housed or group-housed animals were separated by a gate. After two hours, the gate was opened and interactions between this pair of animals were then recorded for 15 min. We found that, after two hours of exposure to fly odors, single-housed flies behaved as sociably as group-raised animals **(Fig. 2d; Supplementary Video 7)**. Thus, olfaction, even in the absence of touch and taste, is sufficient to drive sociability learning.

To further test the importance of olfaction in sociability learning, we compared the sociability of wild-type animals to anosmic animals with mutated olfactory co-receptors (*IR8a*^1^*, IR25a*^2^*, GR63a*^1^*,and ORCO*^1^) ^8^. As expected, wild-type animals that were group-housed for one day were more sociable than flies that were single-housed **(Fig. 2e, left)**. However, both group and single-housed anosmic mutant animals exhibited low sociability. Both were statistically indistinguishable from single-housed wild-type animals **(Fig. 2e, right)**, demonstrating that olfaction is required for sociability learning.

These data raise two further questions about the specificity of the odors required. First, does this odor-mediated learning lead to less fearful, more undisturbed behaviors towards conspecifics only, or to any other species, due to a general reduction in behavioral excitability? Second, to what extent is sociability learned from exposure to conspecific odors specifically, versus any animal odor? To distinguish among these possibilities, we tested how group-housed *Drosophila melanogaster* behave towards the cowpea seed beetle, *Callosobruchus maculatus*, a comparably sized species that is easy to obtain and maintain.

We recorded the behaviors of flies paired with beetles and found that, even though they were group-housed with other flies, they nevertheless reacted fearfully towards beetles **(Fig. 2f, middle; Supplementary Video 8)**. This was true even for flies that were raised housed with both flies and beetles for one day **(Fig. 2f, right)**. In line with this, group raised *Drosophila melanogaster* also reacted fearfully towards group raised *Drosophila simulans* **(Extended Data Fig. 7)**, a distant cousin with a distinct set of cuticular hydrocarbons ^38^. However, co-housed *D. mel.* and *D. sim.* have an intermediate level of sociability, perhaps due to the overlap in pheromonal signatures ^38^. Taken together, these results are consistent with a scenario in which sociability learning is tuned to conspecific odors and only modulates reactions towards conspecifics; it is not due to habituation to any odor in the fly’s surroundings, nor to a general decrease in behavioral excitability. Flies will react fearfully towards other species, but can learn to become sociable towards one another through conspecific odor exposure.

### The mushroom body regulates fearful reactions and sociability learning

Our results suggested the existence of two neural pathways: one mediating visually-driven fearful reactions exhibited by single-housed flies towards other animals **(Extended Data Fig. 6)**, and another mediating conspecific odor-dependent sociability learning **(Fig. 2c-f)**. Because fearful reactions appear to be innate (they occur early post-eclosion), we posited that they may be mediated by visual circuits in the optic lobe projecting to the lateral horn (LH) ^53^, an area implicated in innate behaviors ^54,55^. Alternatively, visual circuits might directly recruit descending neurons (DNs) to drive reflexive motor behaviors, as has been shown for escape responses ^56,57^. We hypothesized that the odor-driven sociability learning pathway might include neurons responsible for detecting pheromones, information which eventually reaches the mushroom body, a brain region responsible for olfactory learning in insects ^58,59,60^. The mushroom body has also been implicated in social attraction ^26^ as well as in courtship conditioning, another form of social learning in which a male-deposited mated female pheromone, cVA, becomes associated with the failure to successfully mate ^14^. Alternatively, sociability learning might arise through the influence of neuromodulators or neuropeptides on innate fear reaction pathways ^61,62^. In this case, the systems for fearful reactions and sociability learning might heavily overlap.

To distinguish between these different mechanistic models, we performed two large-scale neural silencing screens using 188 transgenic driver lines that specifically target candidate neurons for vision, olfaction, higher order sensory processing in the lateral horn and mushroom body, neuromodulation, neuropeptides, and descending pathways **(Extended Data Fig. 8a; Fig. 3a)**. In the first neural silencing screen we aimed to identify neurons responsible for generating fearful reactions. Therefore, we quantified the sociability indices of single-housed animals with neurons silenced **(Extended Data Fig. 8b)**. Here we would expect that silencing neurons responsible for fearful behaviors would cause single-housed animals to be less reactive and thus appear more sociable. In the second silencing screen we intended to uncover neurons responsible for sociability learning. Therefore, we quantified the sociability indices of animals that were initially single-housed but then group-housed for one day prior to recordings **(Extended Data Fig. 8c)**. Here we would expect that silencing neurons regulating sociability learning should cause these group-housed animals to act more fearfully towards one another. To silence neurons, we blocked synaptic transmission by constitutively expressing tetanus toxin (TNT). We note that this does not block volume transmission ^63^ nor influence gap junctions, which may increase the number of potential false negatives.

**Fig. 3:**
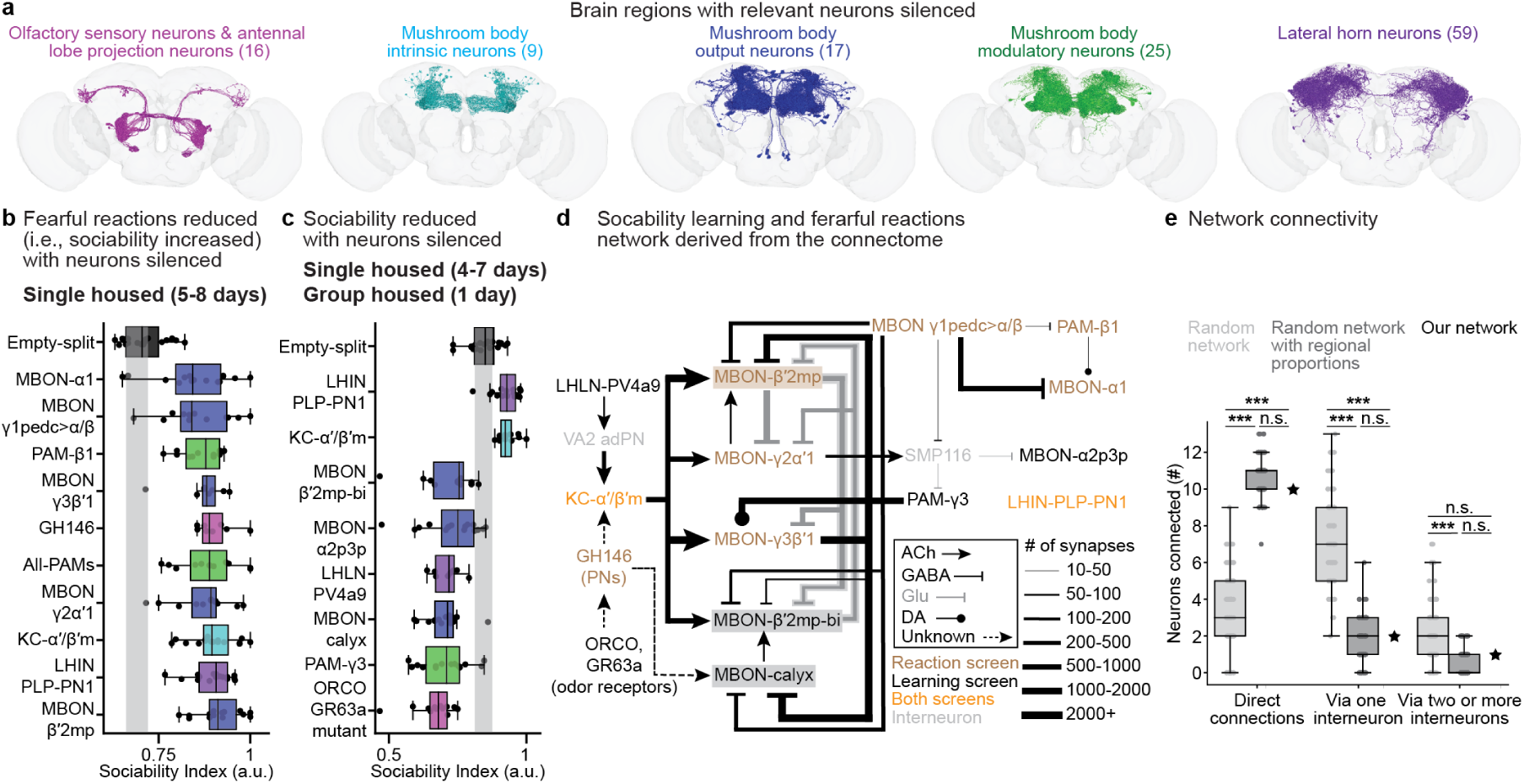
Mushroom body neurons are enriched in a brain network regulating fearful reactions and sociability learning. **(a)** Fly brain connectome renderings of up to 100 neurons in the brain regions with hits from neural silencing screens. Highlighted are olfactory sensory neurons, antennal lobe projection neurons, mushroom body intrinsic neurons, mushroom body output neurons, mushroom body modulatory neurons, and lateral horn neurons. **(b–c)** Driver lines statistically different from the empty-split-Gal4 driver line used as a control in neural silencing screens for **(b)** fearful reaction and **(c)** sociability learning. For hits from the fearful reaction screen, neural silencing causes an increase in sociability despite the absence of conspecific exposure. For the sociability learning screen most hits block the learning of sociability despite 1 day of exposure to conspecifics. Exceptionally, silencing LHIN-PLP-PN1, or KC-α’/β’m appears to increase sociability. Neurons are color-coded by brain region as in panel **a**. Gray-shaded area indicates the 95% bootstrapped confidence interval for control (empty-split-Gal4) data. **(d)** A fly brain connectome-based network derived using the hits from neural silencing screens in panels **b** and **c**. Included are direct connections between nodes as well as indirect connections requiring at most one interneuron. Only LHIN-PLP-PN1 could not be connected to this network given these constraints. Neuron names are color-coded by the screen(s) from which they were derived. Connectome-derived neurotransmitter types and synapse numbers are represented by arrowhead type, grayscale level, and line thickness/type. **(e)** Analysis of the likelihood of obtaining an equivalently interconnected network with the neurons in our silencing screens. Numbers of neurons connected either (left) directly, (middle) with one intermediate interneuron, or (right) with two or more indirect interneurons for: (light gray) a network derived from randomly sampled screened neurons, (dark gray) randomly sampled screened neurons with an equivalent regional enrichment as in our real network, and (black star) our real network. For box plots, the box limits indicate the central 50% of the data and the line inside the box depicts the median value. The whiskers extend out of the box to the full range of the remaining data. Data points placed beyond the whiskers are outliers. The Empirical Cumulative Distribution Function (ECDF) was used to estimate the relative position of our model within the distribution of randomly generated networks. We compared the generated networks using a Mann–Whitney statistical test. *** *P <* 0.0001, n.s. *P* ≥ 0.01.

Remarkably, for both neural silencing screens we observed that the majority of hits (i.e., those significantly different from an empty-split-Gal4 control) are in the mushroom body (*P <* 0.05, Mann– Whitney tests followed by a Holm correction for multiple comparisons). Specifically, in the screen for neurons controlling fearful reactions, we identified ten hits which increased the sociability of single-housed animals **(Fig. 3b)**—MBON-α1, MBON-γ1pedc*>*α/β, PAM-β1, MBON-γ3β’1, most antennal lobe projection neurons (‘GH146’), all dopaminergic PAM neurons (‘All-PAMs’), MBON-γ2α’1, KC-α’/β’m, LHIN-PLP-PN1, and MBON-β’2mp. In the screen for neurons regulating sociability learning, we identified eight significant hits, six of which decreased sociability **(Fig. 3c)**—MBON-β’2mp-bi, MBON-α2p3p, LHLN-PV4a9, MBON-calyx, PAM-γ3, ORCO-GR63a mutant—and two that appeared to increase sociability—LHIN-PLP-PN1, KC-α’/β’m. Notably, these latter two cell classes were also identified in the fearful reaction screen. Thus, rather than influencing learning specifically, their silencing may instead reduce a fly’s ability to react, increasing its apparent sociability.

Our observation that mushroom body neurons mediate both fearful reactions towards other animals as well as sociability learning suggests that the brain, rather than relying on two loosely connected pathways for innate and learned behavior, might instead employ a single interconnected network. To explore this network further, we used the fly brain connectome to construct a graph of connections between 15 hits from both of our neural silencing screens. This excludes a broad line targeting the PAM cluster of dopamine neurons (‘All PAMs’) and the duplication of two cell types across both screens. To generate this graph, we searched for direct connections between our neurons of interest and, in their absence, interneurons that enable indirect connections via one hop. This yielded a tightly interlinked network requiring only two additional interneurons **(Fig. 3d)**.

In this network odor signals converge upon mushroom body Kenyon cells which route these signals to a highly interconnected set of mushroom body output neurons (MBONs) and their associated PAM dopaminergic neurons. Importantly, the directly interlinked nature of this network could not arise trivially from building similar networks out of 13 random cell types taken from our screen. However, our graph does resemble randomly generated networks that conserve the proportion of neuronal ‘hits’ for each brain region **(Fig. 3e)**. Thus, the prominence of the mushroom body among hits in our screens is significant and their functional properties may inform the mechanism whereby fearful reactions are transformed into sociable behavior.

### MBONs encode behavioral reactions and long timescale social exposure

The prominence of mushroom body neurons in both fearful reactions as well as sociability learning suggests several potential models for how such a network might regulate social valence. These models assume that mushroom body output neurons lie just upstream of action selection circuits driving either fearful reactions or social behaviors. On one hand, the mushroom body might contain two functionally independent pathways that are topologically connected but that have a relatively minor impact upon each others neural dynamics. The first pathway might drive innate fearful behavioral reactions while the other might accumulate olfactory experience to regulate downstream action selection circuits. In another model, the mushroom body might directly convey behavioral decisions using two functionally dependent and interacting pathways: one driving fearful reactions and another that is strengthened by conspecific odor exposure and directly inhibits the first pathway. This latter possibility is suggested by inhibitory connections among MBON hits from our fearful reaction and sociability learning screens **(Fig. 3d)**. These models might be tested by measuring the activity of these key MBONs during sociability learning. In the first ‘functionally independent’ model, the fearful reaction pathway may remain cryptically active even after sociability is learned. In the second ‘functionally dependent’ model, the activity of this innate reaction pathway may become suppressed during learning.

To distinguish between these models, we recorded the activities of MBONs in our network over the course of two hours of inter-fly interactions. We focused on three cell classes representing core neurons in our network **(Fig. 3d; highlighted)**: MBON-β’2mp, a hit from our reaction screen **(Supplementary Video 9)** that was previously shown to modulate olfactory decisions by driving odor avoidance when activated and odor attraction when silenced ^64^; MBON-β’2mp-bi, a hit from our learning screen **(Supplementary Video 10)** that inhibits MBON-β’2mp and is known to play a role in innate CO_2_ avoidance ^65^; and MBON-calyx, a hit from our learning screen **(Supplementary Video 11)** for which little is known but which directly excites MBON-β’2mp-bi and receives inputs not only from Kenyon cells (like most MBONs) but also directly from antennal lobe projection neurons closer to the olfactory sensory periphery.

We gained optical access to MBONs in the brain by dissecting out the dorsally-located head vertex as well as underlying fat bodies and tracheae **(Extended Data Fig. 9)**. Using a custom designed experimental system **(Extended Data Fig. 10a)** we could perform two-photon microscopy recordings of our MBONs of interest **(Extended Data Fig. 10b–c)** in tethered animals behaving on a spherical treadmill while they were exposed to another freely moving fly for two hours **(Supplementary Video 12)**. We took this approach rather than simply presenting the tethered fly with odors (i.e., fly pheromones) to make this paradigm as similar as possible to our behavioral experiments. To measure the proximity of this pair of animals, we tracked the position of the freely walking fly using two orthogonal camera views ^45^. Here, as a proxy for locomotor reactions of the tethered fly, we measured accelerations in rotations of the spherical treadmill ^66^**(Supplementary Video 13)**.

Due to the large volume of data acquired by our system and to minimize perturbations of the tethered fly, we measured MBON activity for five epochs of five minutes each over the course of two hours **(Fig. 4a)**. First, we recorded MBON spontaneous activity while the fly was alone in the social arena (‘Alone’). Two minutes later we placed another fly into a nearby chamber for five minutes to test if MBONs already respond to the presence (and scent ^67^) of this other fly (‘Before’). After another two minutes, we allowed the waiting fly to freely enter the arena while recording neural activity during these first interactions (‘During’). Finally, to obtain data after social exposure, we recorded neural activity for five minutes after one (‘After 1 h’) and then two hours (‘After 2 h’) of conspecific co-habitation. We confirmed that, although they were tethered with their brains exposed, single-housed animals express fear-like locomotor accelerations during proximity events and that these reactions diminish to group-housed levels after two hours of social exposure **(Extended Data Fig. 11)**.

**Fig. 4:**
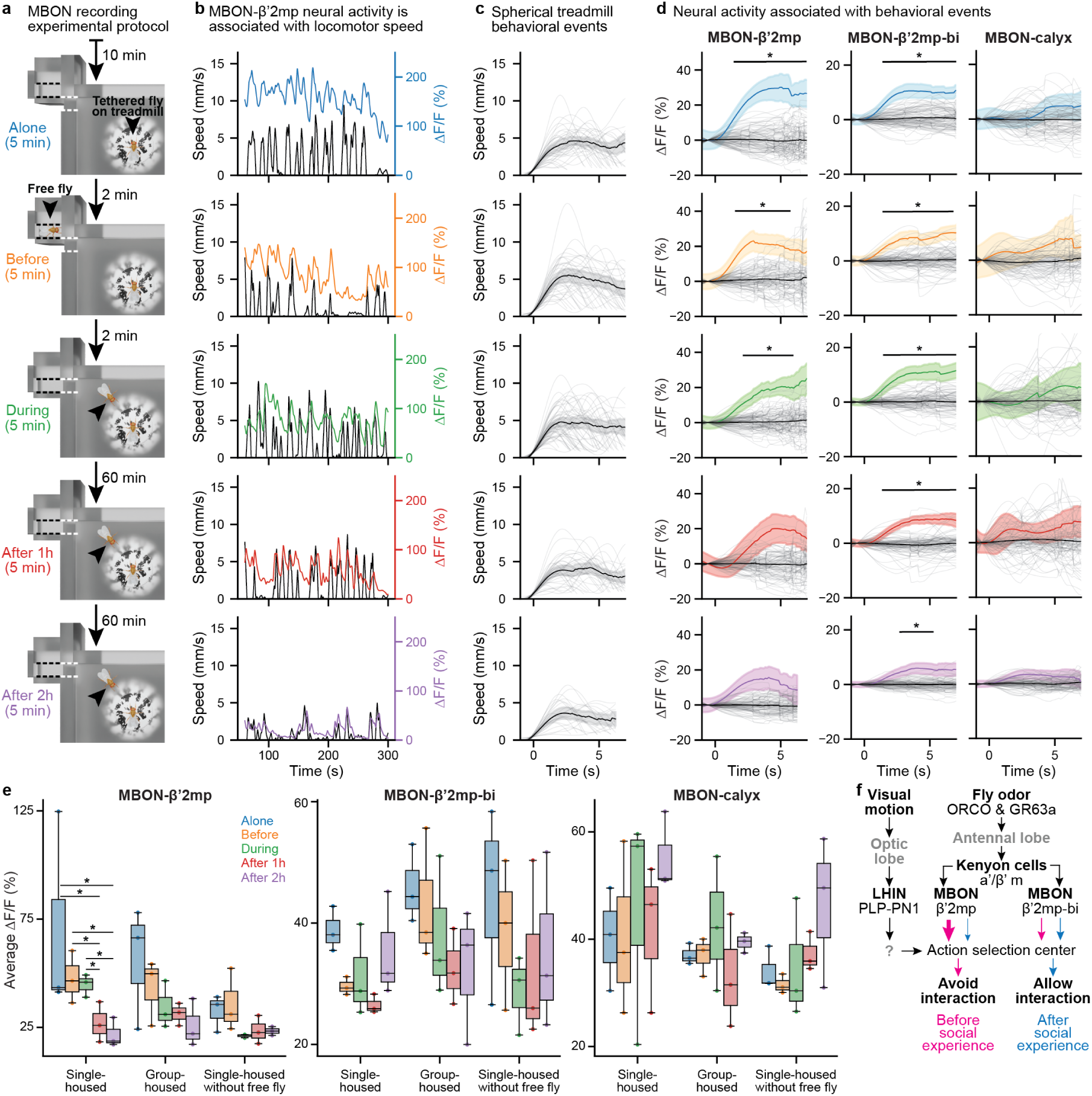
Mushroom body output neuron dynamics are correlated with behavioral events and long-term sociability learning. **(a)** Experimental protocol used to assess the degree to which MBON activities reflect the presence or position of another freely behaving fly, locomotor reactions of the tethered fly, or long-term sociability learning. After flies habituated on the spherical treadmill for 10 minutes, MBONs were recorded for five minutes to capture their spontaneous activity in the absence of another fly (‘Alone’). After 2 min, a second fly (black arrowhead) was placed into a waiting room (black dashed lines), out of sight but potentially detectable through olfaction (‘Before’). Then, 2 min later, this second fly is allowed to enter the arena through an entrance (white dashed lines) and to interact with the tethered animal (‘During’). The fourth and fifth epochs of 5 min recordings are one hour (‘After 1 h’) and two hours (‘After 2 h’) after the second fly was introduced into the arena. This allowed us to identify potential correlates of sociability learning. **(b)** Example treadmill-derived locomotor speed (black) and MBON-β’2mp ΔF/F neural activity traces for a single-housed fly across the five experimental epochs (color-coded as in panel **a**). Neural activity appears to correlate with early epoch locomotor bouts and baseline activity diminishes dramatically across epochs. **(c)** Behavioral events were selected for each epoch by thresholding treadmill speed and time-locking to event onset (0 s). Overlaid are individual events (translucent traces) and the mean of all events for three animals during a given epoch (solid black traces) along with the 95% confidence interval (gray transparency). **(d)** Mean neural activity and 95% confidence interval associated with behavioral events (traces color-coded by experimental epoch) for MBON-β’2mp (left), MBON-β’2mp-bi (middle), and MBON-calyx (right). Overlaid are traces (translucent gray) representing the mean activity of each set of randomly selected events. The means and confidence intervals of all sets of randomly selected events are also shown for comparison (black traces). Mean traces were compared at each time point using a permutation test followed by a correction for multiple comparisons using the false discovery rate for positively correlated tests. * *P <* 0.05. **(e)** Average MBON baseline activity (i.e., segments between behaviors) across flies. Each dot represents the mean ΔF/F for a given fly on a given epoch (color-coded) (n=3 for each condition). Different epochs within the same fly condition (single-housed or group-housed) were compared using a Kruskal–Wallis test followed by a post hoc Conover’s test and correcting for multiple comparisons using the false discovery rate for positively correlated tests. * *P <* 0.05. All other comparisons were statistically similar to one another (i.e., *P* ≥ 0.05). For box plots, the box limits indicate the central 50% of the data and the line inside the box depicts the median value. The whiskers extend out of the box to the full range of the remaining data. Data points placed beyond the whiskers are outliers. **(f)** Schematic illustrating how information transmission may be altered by social experience to enable the transition from fearful reactions (‘Avoid interaction’) to learned sociability (‘Allow interaction’). Shown are neural drives that are either prior to (magenta) or following (blue) social experience.

First we asked whether MBONs might encode the generation of these fear-like behavioral reactions to nearby conspecifics. When we examined MBON neural dynamics over these experimental epochs, we noted a strong relationship between the activity of MBON-β’2mp and locomotor speed **(Fig. 4b)**. To more precisely quantify this relationship, we defined and time-aligned behavioral events—periods when tethered flies generated bouts of fictive locomotion **(Fig. 4c)** (see Methods). We then compared neural activity during these time windows as a function of housing condition, MBON identity, and experimental epochs. Indeed, we found significantly higher MBON-β’2mp activity levels when single-housed animals were behaviorally active. This activity diminished after two hours of social exposure **(Fig. 4d, left column; Extended Data Fig. 12)**. Notably, we did not observe this diminishment in MBON-β’2mp activity if single-housed animals were not exposed to a freely-behaving conspecific **(Supplementary Information File, Figure 7)**. By contrast, MBON-β’2mp-bi activity levels were significantly higher when single-housed animals became behaviorally active throughout the entire experiment, irrespective of their social exposure **(Fig. 4d, middle column)**, as long as another fly was in the arena **(Supplementary Information File, Figures 9** and **13)**. It is known that many neurons in the fly brain become active during locomotion ^68^. However, we did not observe that MBONs uniformly encode behavior: MBON-calyx activity was not clearly related to fly locomotion at any stage of the experiment **(Fig. 4d; right column)**. The activities of these MBONs also did not relate to the position of the freely behaving fly **(Supplementary Information File)**. Thus, they do not appear to encode the intensity of fly odors.

In addition to the short timescale correlation of specific MBON activity to behavioral reactions, we wondered if MBONs might also encode the experience of long timescale social exposure. We analyzed neural activity across each epoch of our experiment and indeed observed a marked and continuous reduction in MBON-β’2mp baseline **(Fig. 4b)** neural activity over two hours. By calculating the median change in fluorescence for each epoch we found that the reduction in MBON-β’2mp baseline activity was significant for single-, but not group-housed animals **(Fig. 4e, left)**. As well, this reduction was not observed when single-housed animals were not exposed to a freely moving conspecific animal. Importantly, this change in baseline activity was not due to photobleaching as it was not observed for the other two MBONs **(Fig. 4e, middle and right)**. On the contrary, MBON-calyx showed a slight, but not significant, increase in baseline activity in single-housed animals after two hours of conspecific exposure.

Taken together, these findings are consistent with a model in which fly pheromones activate olfactory pathways (ORCO- and GR63a-expressing OSNs and their downstream projection neurons targeting Kenyon cells in the α’/β’m subregion) and ultimately engage MBONs including MBON-β’2mp and MBON-β’2mp-bi. Simultaneously, visual motion of this neighboring fly stimulates visual pathways that project to the central brain, including LHIN-PLP-PN1, a hit found in both neural silencing screens. At first, these visual and olfactory circuits will bias downstream action selection circuits, causing flies to avoid one another **(Fig. 4f, magenta arrows)**. However, chronic exposure to fly pheromones will suppress the baseline and locomotor-related activity of MBON-β’2mp, allowing MBON-β’2mp-bi to strongly influence downstream action selection circuits and drive sociable reactions towards other flies **(Fig. 4f, blue arrows)**.

## Discussion

Here we have shown that *Drosophila melanogaster* become sociable through exposure to conspecific odors and their processing by specific circuits in the mushroom body, the principal center for learning and memory in the insect brain. This establishes sociability as an ethologically foundational learned behavior. But why might conspecific sociability be learned rather than hard-wired? Our experiments following only one day of social isolation suggest that fear may be the default state of the adult fly. This might be an appropriately conservative starting point for socially naive flies to avoid interacting with potentially dangerous animals. Only after sufficiently long exposure to conspecific odors should one’s guard be let down. In addition, we note that insect cuticular hydrocarbons are not immutable but can be considerably modified by environmental factors like temperature and humidity ^69^. Thus, learning may also provide a means for maintaining flexibility that allows animals, upon eclosion, to discover the actual signature of conspecific pheromones in a particular environment.

The precise fly pheromone that drives sociability learning remain unknown. Unlike the specific role for cVA in courtship conditioning ^14^, we believe that it may include a broad palette of odors. In our screens we did not observe a significant effect when silencing OSNs responsive to cVA (expressing OR67d ^37^), or to other pheromones (expressing OR88a, OR47b, or OR65a ^70,71^). Additionally, we did not observe a significant impact of silencing broad GAL4 driver lines targeting large fractions of OSNs expressing the GR63a, ORCO, IR8a, or IR25a co-receptors nor for flies with individually mutated *Gr63a*, *Ir8a*, *Ir25a*, or *Orco* co-receptors. On the other hand, the combined mutation of both *Orco* and *Gr63a* strongly perturbed sociability learning. Thus, because ORCO is a co-receptor for pheromonal ORs and GR63a ^72^ has been implicated in *Drosophila* stress odorant sensing ^73^ the conspecific odor signature in question may be a complex combination of pheromones. Similarly, silencing individual classes of visual projection neurons did not suppress fearful reactions—nevertheless, vision is required as single-housed flies in the dark do not react before being touched. This suggests that fearful reactions are likely driven by a combination of cell classes that encode a variety of visual features. This might explain why *D. mel.* were less fearful of *D. sim.*—a closely related species with some pheromonal overlap and a very similar visual morphology—compared with the more evolutionarily distant and visually distinct beetle species.

Conspecific odors appear to drive sociability learning through the engagement of specific neurons that form a tightly knit network in the mushroom body and that are implicated in associative learning (e.g., MBON-α1 ^74^, PAM-α1 ^74^, MBON-β’2mp ^75,76^, MBON-γ1pedc ^77^ or MBON-γ1pedc*>*α/β ^78,79,80,81^, MBON-γ2α’1^81,82,83^). It has been shown that flies remember the omission of punishment as an experience of positive valence ^84^. Thus, one way in which these networks might drive sociability learning is by associating fly odors with the absence of threatening behaviors (e.g., rapid approach or forceful impact). Learned associations derived from another fly’s behavior have previously been shown in courtship conditioning ^14^.

Notably, many of our mushroom body neuron hits disrupted fearful reactions when silenced. This implies that overlapping circuits regulate both innate reactions and sociability learning. Mushroom body-mediated modulation of innate circuits has previously been suggested as a mechanism for odor-guided behavior ^85^ including innate attraction to yeast when hungry ^86^. This is in line with a prominent ‘valence-balance model’ ^87^ which suggests that MBON populations represent a vector with each element driving a specific behavioral valence—aversion or attraction. Learning (or neural silencing in the case of our screens) biases the influence of MBONs upon downstream action selection circuits (e.g., potentially in the SMP) towards one valence or the other. For example, in the case of courtship conditioning, failed mating is thought to depress circuits that drive attraction to females smelling of cVA ^88^ resulting in indifference or aversion. In the case of sociability, even a slight MBON bias towards attraction could explain the observation that flies cluster into groups ^27^. This is supported by the observation that social attraction requires Kenyon cells in the gamma lobe of the mushroom body ^26^.

This ‘valence-balance model’ is also in line with the relationship we observe between the activities of MBON-β’2mp and MBON-β’2mp-bi and their respective importance in generating fearful reactions and learning sociability (as determined by neural silencing). We found that the activity of MBON-β’2mp, which is crucial for generating fearful reactions, closely tracks bouts of walking. However, it is suppressed after two hours of conspecific exposure. MBON-β’2mp-bi, which is key for learning sociability, is also time-locked to bouts of walking. However, it shows no decay in activity over two hours. From these data we hypothesize that conspecific odors diminish the influence of MBON-β’2mp while leaving unchanged the impact of MBON-β’2mp-bi on downstream action selection centers. We envision that these two MBONs (among many others) compete for behavioral control: MBON-β’2mp initially dominates to drive fearful reactions but following prolonged conspecific odor exposure MBON-β’2mp-bi wrests control to encourage sociable behaviors. Thus, the mushroom body network regulating sociability may represent an ancient learning pathway with a foundational role in gating conspecific interactions, the expression of most social behaviors, and the emergence of communal living.

## Methods

### Animal stocks and husbandry

The majority of experiments were performed on adult *Drosophila melanogaster*. Flies were raised at 25 ^◦^C and 50% humidity on a 12 h light–dark cycle and collected as pupae after stage P14 ^89^. Females were identified by selecting against the presence of sex combs. Pupae were then placed either singly or in groups of 5–10 in vials with standard cornmeal food. The parents of collected pupae were 2–14 days-post-eclosion (dpe). All experiments were conducted during the evening in zeitgeber time (ZT 11–14).

#### Non-transgenic animals

Behavioral experiments were performed on wild-type PR (Phinney Ridge, Seattle) *Drosophila melanogaster*, wild-type *Drosophila simulans.196* (Drosophila Species Stock Center [DSSC]: 14021-0251.196) and wild-type cowpea beetles *Callosobruchus maculatus* (Pocerias, Switzerland). *D. sim* flies were raised and sorted under the same conditions as *D. mel*. *C. mac* were raised at room temperature (22–24 ^◦^C) on a ∼15–9 light-dark cycle (natural light during July in Switzerland) in a plastic container filled with cowpeas. We used both male and female beetles and did not control for age or mating status.

#### Transgenic animals used for neural silencing screens

Behavioral experiments used to identify neurons responsible for fearful reactions and sociability learning were performed on the progeny of crosses between Gal4 driver line males (see **Table 1**) and UAS-TNT.E2 females (Bloomington number: 28837). This allowed us to express tetanus toxin (TNT) and prevent synaptic transmission in neurons of interest. Crosses were flipped into new food vials every 2–3 days for three iterations. Among progeny, female pupae were identified and single-housed for 4–7 dpe. For the sociability learning screen, flies were then grouped for 24 h prior to the experiment.

**Table 1:**
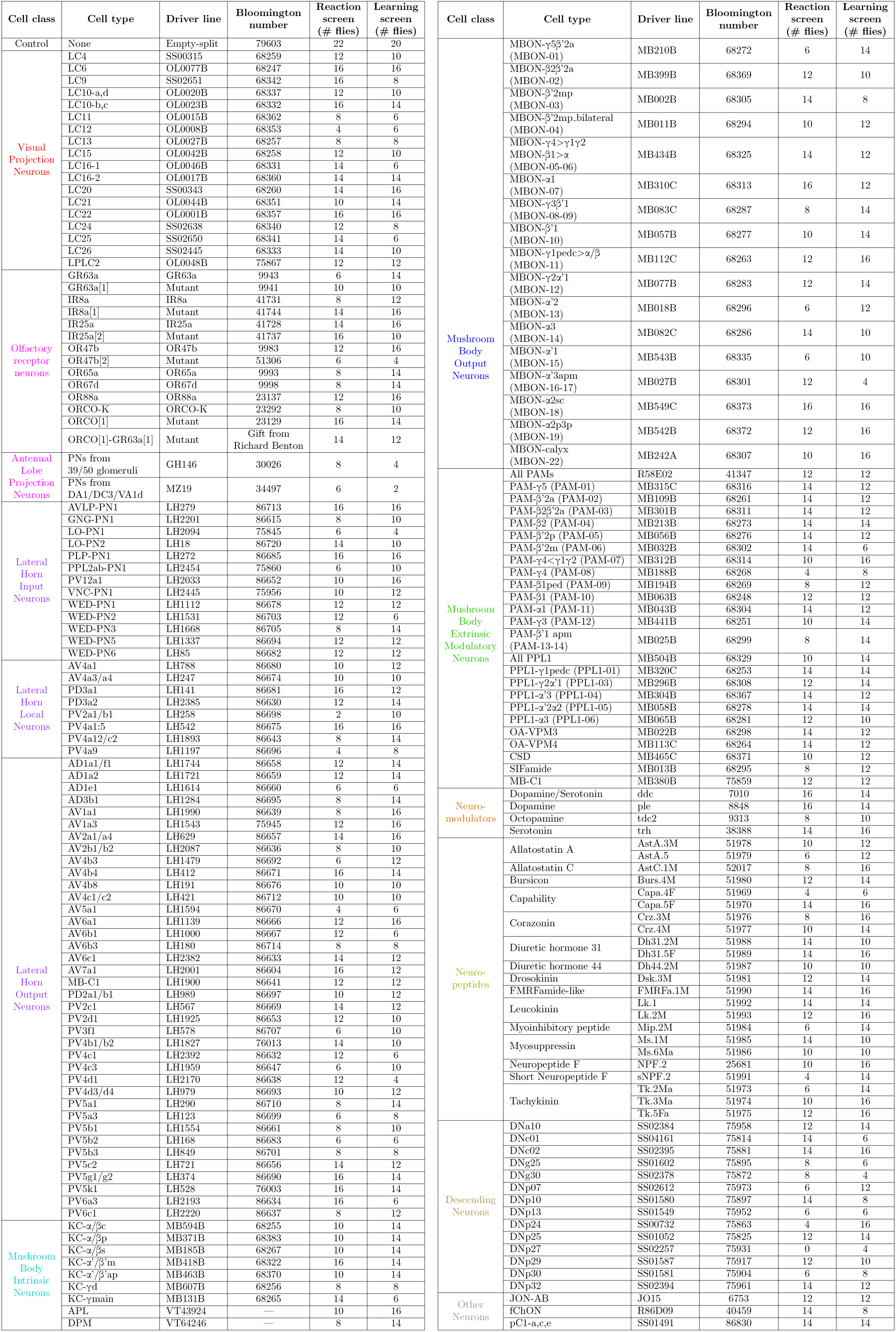
Driver lines used in fearful reaction and sociability learning neural silencing screens. Columns show: the main cell type targeted; the driver line/mutant used; the Bloomington Stock Center number; and the number of flies recorded for the reaction and learning screens.

#### Transgenic animals used for neural recordings

Experiments to record the activity of MBONs of interest were performed on the progeny of crosses between Gal4 driver line males targeting MBON-β’2mp, MBON-β’2mp-bi, or MBON-calyx (see **Table 1**) and fluorescent reporter expression females (*+; UAS-GCaMP6s; UAS-tdTomato*). Crosses were flipped into new food vials every 2–3 days for three iterations. Among progeny, female pupae were identified and either single- or group-housed for 5–8 dpe. Freely moving conspecific flies were group-housed wild-type PR females the same age as tethered, recorded animals.

### Behavioral experiments

Pupae were sorted using a wet paint brush. To group single-housed animals for one day we transferred 5–9 single-housed flies from food vials into 5 mL falcon tubes (Falcon, 352063, polypropylene round-bottom tubes) using a funnel. Then, these flies were transferred together into an empty food vial. To place flies into arenas we transferred them from food vials into falcon tubes and anesthetized them on ice for 2 min. Single-housed flies were kept alone in falcon tubes. Then, flies were poured onto a cooled petri dish over ice and loaded into behavioral arenas using a paint brush **(Extended Data Fig. 1a,c)**. For male-female experiments, during cold anesthesia, we removed males’ wings under a microscope to prevent the production of courtship song and wing displays. Flies were left in arenas for 5 min to recover from cold anesthesia before beginning each experiment. After this recovery time, experiments were initiated by manually opening the arena gates **(Extended Data Fig. 1b,d)**. For the odor-enriched arena experiments, flies were left for 2 h with the gates closed prior to recording. The specific details of each behavioral experiment are summarized in **Table 2**. We discarded flies that showed any physical defect including broken or missing legs, twisted or folded wings, or if they were abnormally small.

**Table 2:** Details for behavioral experiments. Columns show: the behavioral experiment; genotype; sex; social condition of the flies used; the duration of experiments; the group used as a control to compute metric thresholds; and important notes for each experiment.

| Experiment | Genotype | Sex | Social condition (days) | Recording time (minutes) | Used for thresholds | Notes |
| --- | --- | --- | --- | --- | --- | --- |
| Sociability control | Wild-type (PR) | Female-Female<br>Male-Male<br>Female-Male | Single-housed (5-8)<br>Group-housed (5-8) | 10 | Female-Female group-housed | We removed the wings of males in Female-Male experiments. |
| 1 dpe sociability | Wild-type (PR) | Female-Female | Single-housed (1)<br>Group-housed (1) | 15 | Female-Female group-housed |  |
| Learning over 24 h | Wild-type (PR) | Female-Female | Single-housed (5)<br>Group-housed (5)<br>Single-housed (4) then grouped (1) | 15 | Female-Female group-housed |  |
| Forgetting | Wild-type (PR) | Female-Female | Single-housed (5, 8, 12)<br>Group-housed (5, 8, 12)<br>Group-housed (4) then isolated (1, 4, 8) | 15 | Female-Female group-housed (8) |  |
| Learning over 2 h | Wild-type (PR) | Female-Female | Single-housed (5-8)<br>Group-housed (5-8) | 120 | Female-Female group-housed |  |
| Learning in the dark | Wild-type (PR) | Female-Female | Single-housed (5-8)<br>Group-housed (5-8) | 120 | Female-Female group-housed | We turned off the cool white LEDs inside the behavioral system enclosure. |
| Odor enriched | Wild-type (PR) | Female-Female | Single-housed (5-8)<br>Group-housed (5-8) | 15 | Female-Female group-housed | Flies were mounted in the arenas 2 h before recordings. Flies used to enrich the odor were group-housed. |
| Anosmic mutants | Wild-type (PR)<br>*IR8a<sup>1</sup>*, *IR25a<sup>2</sup>*,<br>*GR63a<sup>1</sup>*, and *ORCO<sup>1</sup>* | Female-Female | Single-housed (5-8)<br>Group-housed (5-8) | 15 | Wild-type female-female group-housed | Anosmic animals can still smell using the co-receptor *IR76b* which detects polyamines<sup>90</sup>. |
| Cowpea beetle | Wild-type (PR, *D. mel*)<br>Wild-type (*C. mac*) | Female (*D. mel*)-<br>Female/male (*C. mac*) | Group-housed with conspecifics (5-8)<br>Single-housed (4-7) then grouped with heterospecifics (1) | 15 | *D. mel* female-female group-housed | Beetles were mounted into the behavior arenas using the same procedure used for flies. Beetles were always placed on the right side of the arena. |
| *Drosophila simulans* | Wild-type (PR, *D. mel*)<br>Wild-type (*D. sim*, 196) | Female-Female | Group-housed with conspecifics (5-8)<br>Single-housed (4-7) then grouped with heterospecifics (1) | 15 | *D. mel* female-female group-housed | We sorted *D. mel* from *D. sim* using a microscope while flies were anesthetized before mounting them into the arenas. We always placed *D. sim* on the right side of the arena. |
| Screens | Table 1 | Female-Female | Single-housed (5-8)<br>Single-housed (4-7) then grouped (1) | 15 | Empty-splitGal4 female-female group-housed | We recorded at least 6 pairs from each genotype. Group-housed flies were paired with conspecifics from the same or different vials. We only analyzed lines were more than 8 flies showed valid proximity events. |

#### Experimental system

Images were recorded using a PCO.panda 4.2 camera (GLOOR Instruments, CH) with a Milvus 2/100M ZF.2 lens (ZEISS, DE) from the bottom of the arena and through a protected silver mirror at a 45° angle (PF20-03-P01, and H45CN, Thorlabs, Germany). An infrared LED ring light (CCS Inc. LDR2-74IR2-850-LA) was used to illuminate the arena from above. The complete behavioral system was enclosed and illuminated internally with cool white LED strips (FP1-W01D-500, Sloan). The camera was triggered externally using an Arduino UNO to capture frames at 80 fps. Custom Python software was used to acquire 832 × 832 pixel images. These images were processed using adaptive histogram equalization with the CLAHE method in OpenCV. Images were then compiled into videos using FFmpeg with an x265 encoder at a constant rate factor of 15 to make them visually lossless.

Experiments were performed in a temperature and humidity controlled room (25 ^◦^C and 50%). However, the temperature measured at the arenas was 29–31 ^◦^C due to heating by IR illumination. Behavioral arenas are composed of 2 mm thick acrylic glass floors and ceilings and POM static and movable walls. In its entirety, the whole arena is 50 mm × 50 mm. The sub-arenas for each fly pair are 12 mm × 13 mm. Arenas for odor-enriched experiments are made of the same materials except that, in addition, a 2 mm stripe of fine mesh was used to create walls dividing the main arena from six flies used to provide odors. 2D and 3D CAD models to build or customize these arenas are available on Dataverse (See Data availability).

### Behavioral analysis

Because the main arena includes four sub-arenas, we first used custom Python code to crop videos and obtain individual sub-arenas each with a pair of experimental animals. To do this, we first summed the last 50 frames of each recording and calculated its histogram. From this, we obtained grayscale values representing the arenas. These values were then used to threshold the image and obtain a mask of the arena’s walls. Finally, we used the connected components function from OpenCV to obtain the coordinates of each arena.

#### 2D tracking

From these individual sub-arena videos, we tracked the movements of flies using a deep neural network trained with SLEAP ^45^ (v1.2.8). We trained the SLEAP model using a bottom-up approach over a U-Net network. We used 300 manually labeled frames with 11 landmarks (thorax, head, abdomen, left and right wings, and the pretarsi of the six legs). We augmented the training data with rotations (−180^◦^ to 180^◦^), scaling (0.9 to 1.1), Gaussian noise (5 ± 1), contrast (0.5 to 2.0), and brightness (0 to 10). When annotating frames, we did not distinguish between the ventral and dorsal views of the fly (e.g., an annotated left leg will always be a leg to the left of the head but not necessarily the anatomical left leg of the fly). When flies were walking on the walls of the arena we annotated the proximal side of the body only. Once the model was trained we ran 2D pose predictions and the tracker using the Hungarian algorithm to keep track of the identities of multiple animals. We set the instance count in the tracker to two instances unless otherwise explicitly specified.

To track the beetles we first computed and subtracted the median of the images to remove the background. Then, we binarized images using an adaptive mean thresholding implemented in OpenCV. Next, we adapted the pixel neighborhood size until we obtained three large objects: the fly and both sides of the beetle. We used both sides of the beetle because typically the midline of the beetle was always too dark to be recognized as foreground. Thus, we applied a closing morphological transformation to join both parts of the beetle and to clean up the image, leaving only the fly and the beetle. Finally, we used the Hungarian algorithm to keep track of the animals’ identities, and *k*-means clustering if during the thresholding more than two objects were recognized. To distinguish the fly from the beetle we computed the distance between the SLEAP tracking of the fly and this custom tracking of both objects. We assigned the identity of ‘beetle’ to the object farthest from the SLEAP-tracked object.

After obtaining these tracking predictions, data were exported to HDF5 files using SLEAP functions. The prediction, tracking, and export of data were done using the command line interface provided by SLEAP. Finally, to load and use these data, we first replaced missing values with the closest non-NaN values. Then, we checked for possible identity swaps in the tracking by comparing the position of each animal two frames before and after the analyzed frame. If the distance between the time-points ‘after’ and ‘before’ for the same fly was larger than the distance between the time-point ‘after’ for one fly and time-point ‘before’ for the other fly, and this was true for both flies, we assumed there was an identity swap. To fix this we swapped the tracking data for both flies from this time point onward.

#### Proximity events

We define proximity events as when flies were closer than 5 mm apart. The distance is calculated as the mean of the distances across all possible pairings of the head, thorax, and abdomen of both flies. This prevents outliers from distorting the calculated distance. For beetle experiments, we only consider the distance between the thorax of both animals. Then, we define several dynamic conditions to consider an event valid. First, events must last more than 0.5 s. Then, if they last more than 10 s, they are trimmed to 10 s centered on the frame with the closest distance between the flies. If flies exit the 5 mm distance for less than 0.25 s it is still considered the same event. However, at least 75% of the frames need to be within the 5 mm distance. Finally, the closest distance between the flies must be less than 4 mm. If flies had less than one proximity event every 2 min, on average, the experiment was discarded. Furthermore, we did not consider any proximity event during the first 2 min of recording since flies typically begin by exploring the arena.

Next, we divided the proximity events into two classes depending on the flies’ reactions. If the mean velocity of the fly was higher than 2 mm*/*s after the closest distance to the other fly then it was considered a ‘moving’ event. If, on the hand, the mean velocity of the fly was less than 2 mm*/*s then it was considered a ‘standstill’ event. We calculated the distancing efficiency metric for moving events and the freezing immobility metric for standstill events.

#### Generation of behavioral space and clustering of proximity events

To generate a low-dimensional embedding space for proximity events, we first compute the time series describing how flies are positioned and move relative to one another. These time series include: 1. *d*: the distance between two flies (each fly’s position is determined by averaging the coordinates of its head, thorax, and abdomen); 2. *v*^to.^: the velocity component of a fly towards the other fly; 3. *v*^⊥^: the absolute value of the velocity component of a fly perpendicular to the line connecting the two flies; 4. *θ*: the magnitude of the smaller angle between the heading vector of a fly and the line connecting the two flies. The velocity components *v*^to.^, *v*^⊥^, and the angle *θ*, were computed separately for each fly, yielding a total of seven variables (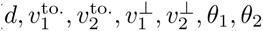).

For each proximity event *i*, we extract from each time series a 40-frame (0.5 s) time window centered at the frame index *t_i_* when the two flies are closest to each other, and concatenate the time windows together. This results in an expansion of the number of dimensions from 7 to 280:

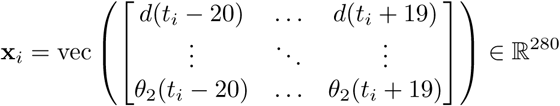

We then reduce the dimensionality from 280 to 2 using UMAP with 100 neighbors ^46,47^. To account for the difference in scales of the distance, velocity, and angle variables, we normalize each subpart of the input data matrix **X**, ensuring that elements corresponding to the same type of variable have a mean of 0 and a standard deviation of 1.

To make the behavioral space more compact, we use a custom distance metric for UMAP:

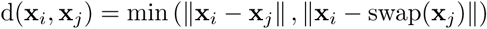

Here, swap(**x**) represents the input vector **x** with the identities of the two flies interchanged (i.e., swapping the values of 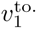. and 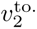, etc.). This adjustment ensures that reciprocal events (such as fly 1 chasing fly 2 and fly 2 chasing fly 1) are treated similarly.

Finally, to cluster proximity events, we performed *k*-means clustering (*k* = 20) on the 2D points in behavioral embedding space, resulting in clusters of visually similar events.

#### Calculating the distancing efficiency metric

Distancing efficiency is calculated using Equation 1 for proximity events in which flies react by moving in any direction. In this equation *AUC_d_* is the integral of the derivative of the distance between the animals (**s**_A_(*t*) − **s**_B_(*t*)) multiplied by the absolute acceleration (|**a**_A_(*t*)|) of the analyzed animal. This integral was estimated by Simpson’s rule over 0.5 s centered on the frame with the closest distance between flies (*t*_0_). *d*_min_ is the value of the closest distance; *th*_min_ and *th*_max_ are the 15th and 85th percentiles, respectively, of the distribution of the closest distance for all the proximity events in the control group. 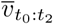 is the mean velocity after the closest distance and 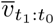 is the mean velocity prior to the closest distance.

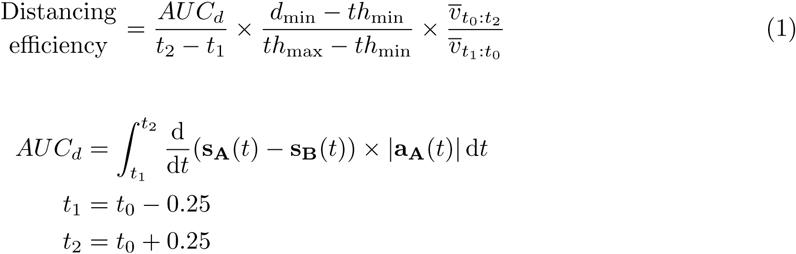

#### Calculating the freezing immobility metric

Freezing immobility is calculated using Equation 2 for proximity events in which the flies react by staying approximately at the same position. In this equation *AUC_f_*is also the integral of the derivative of the distance between the animals multiplied by the acceleration of the analyzed animal. However, here we subtract high frequency movements of the legs (*H*(*s, t*)). This time we also estimate the integral using Simpson’s rule but over 0.25 s starting at the closest distance between the flies (*t*_0_). *H*(*s, t*) is obtained using a Butterworth digital high-pass filter of fifth order with a cutoff frequency of 7 Hz which corresponds to approximately the frequency of leg movements during grooming ^91^. We apply this filter to the mean position of both front and hind legs and then smooth the resulting signal using a Savitzky–Golay filter of second order with a time window of 0.25 s. *v_t_*_0 :*t*2_ is the mean velocity after the closest distance and *v_t_*_1 :*t*0_ is the mean velocity prior to the closest distance.

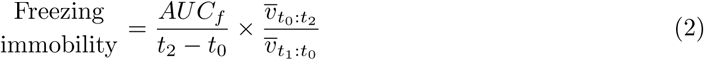

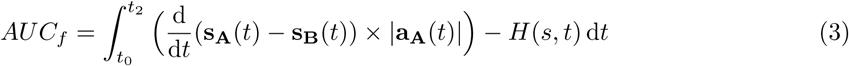

#### Sociability index

To calculate the sociability index we defined thresholds based on associated control experiments. We note that, because each experiment has its own controls, thresholds can be slightly different and the sociability level of controls will always be nearly the same. We chose thresholds on distributions and did not focus on raw ‘distancing efficiency’ and ‘freezing immobility’ values because we were looking to distinguish between control and experimental groups, not individual proximity events. To calculate the threshold values we computed both metrics for control data and identified the 85th percentile value of each metric. Then, we applied these thresholds to distributions from the experimental groups by quantifying the proportion of proximity events from each of the two classes that were below their respective thresholds. In other words, we compared what proportion of the data from each experimental group was within 85% of the control data. Importantly, the statistically significant difference between single and group-housed sociability indices were not substantially impacted by the percentile threshold chosen **(Extended Data Fig. 3)**. We made pairwise comparisons using a Mann–Whitney statistical test. We made multiple comparisons using a Kruskal–Wallis test followed by a post hoc Conover’s test with a Holm correction.

### Connectomics analysis

We used the female adult fly brain (FAFB) connectomics dataset ^31,32^ from Codex (v783) and the hemibrain dataset ^92^ from NeuPrint ^93^ (v1.2.1) to explore the connections between neural silencing screen hits. After loading the data we used a custom Python script to obtain neuron IDs from cell type names. We used the equivalent cell type names in Table 3 to find neurons from our screens that had been identified and labeled in connectomics datasets. We searched for the name in the “hemibrain type”, “label”, and “cell type” columns, in that order of priority. Then, for each cell we searched for every downstream partner with at least 10 synaptic connections. Next, to obtain every direct connection in our network, we identified which of those downstream partners were also hits, excluding self-connections. Then, we did the same analysis but one level further downstream. Finally, we took the list of all downstream partners of our hits. These were ‘neurons of interest’ for which we then searched for second-level downstream partners connected to our hits. Thus, we obtained every possible connection within our network via one hop / interneuron. We used the exclusion criteria that interneurons should not be from our screen as ‘non-hits’ and that they should not connect neurons that already have direct connections. Furthermore, we prioritized connecting neurons that were not directly in the network already.

**Table 3:**
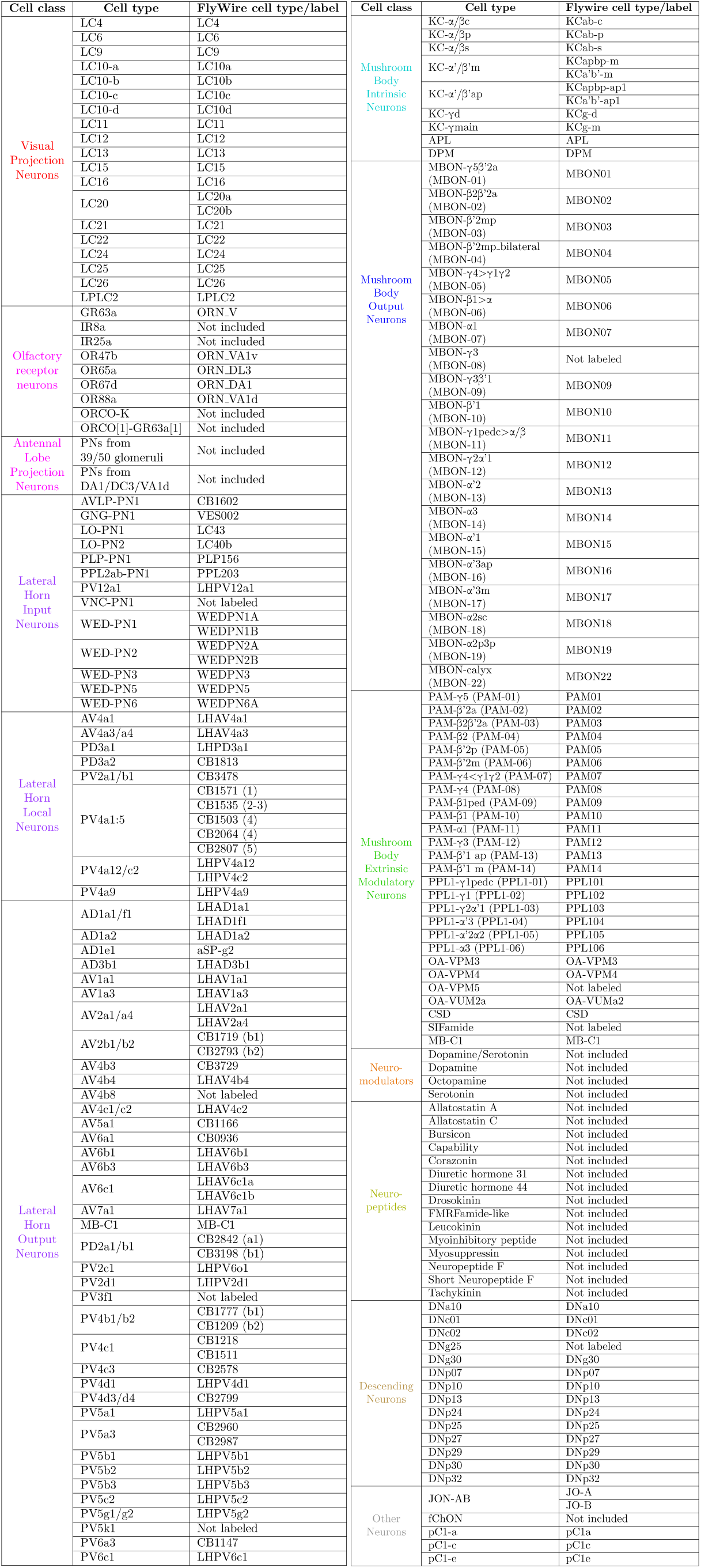
The correspondence of cell types between driver lines and the FlyWire brain connectome. Columns show: main cell types targeted and FlyWire labels for the same neurons.

For statistical analyses of connectivity, we either randomly selected neurons from those tested in our screens that were identified in the connectome (Table 3), or we selected randomly and added the additional constraint that the proportions of neurons from each brain region in our hits should be maintained (i.e., if we had 8 MBONs among our hits and 2 lateral horn neurons (LHNs), to maintain this proportion we would randomly select 8 MBONs and 2 LHNs).

### *In vivo* two-photon calcium imaging

#### Brain dissection

To gain optical access to the brain for functional imaging, we dissected the dorsal head of the fly. First, we anesthetized animals using a cold plate at 6 ^◦^C. Then, we used forceps to remove their wings close to their joints on the thorax. We next mounted the fly onto the imaging stage ^94^ by gluing its scutellum to a metal shim **(Extended Data Fig. 9a)** with UV-curable glue (Bondic, Aurora, ON Canada) and an LED-200 (Electro-Lite Co. Bethel, CT USA). Because proboscis extensions cause the brain to move, we glued the proboscis in place **(Extended Data Fig. 9b)**. Finally, we fixed the head of the fly to the shim by gluing around the dorsal perimeter of the eyes. We carefully spread UV glue around the neck of the fly to prevent leakage of physiological saline **(Extended Data Fig. 9c)**.

For dissections, we filled the stage with extracellular saline ^95^ and cut the vertex using a hypodermic needle (30 G, BD PrecisionGlide, Franklin Lakes, NJ USA). The cut began at the anterior head and followed the lines of bristles along the compound eyes. We completed the cut by carefully sliding the needle below the cuticle and reaching towards the posterior head **(Extended Data Fig. 9d)**. After the cuticle was cut away, we used forceps to remove the fat bodies **(Extended Data Fig. 9e)**, trachea, and ocelli **(Extended Data Fig. 9f)**. This exposed the brain, allowing us to gain optical access to the medial lobe of the mushroom bodies **(Extended Data Fig. 9g)**. We mounted the fly onto our experimental setup and discarded flies that showed any physical defect including broken or missing legs or if arenas leaked saline solution **(Extended Data Fig. 9h)**.

#### Experimental system

We used a Bergamo II (Thorlabs, Germany) two-photon microscope to record mushroom body neural activity. This system performs galvo-galvo scanning at 8 Hz of a 930 nm Ti:Sapphire laser (Mai Tai DeepSee, Spectra Physics) at 12.5 mW to record from a single imaging plane. The fly was tethered and placed upon a spherical treadmill: a foam ball (Last-A-Foam FR-7106, General Plastics, Burlington Way, WA USA, density: 96.11 kg*/*m^3^) of 9.9 mm diameter supported by air flowing at 0.9 L*/*min. Airflow was controlled using a mass flow controller (Vögtlin Instruments, Switzerland). Surrounding the imaging stage, we built a social arena that allows a second freely moving fly to behave around the tethered fly. We used two IR-sensitive cameras (Basler, acA1920-150µm, Germany) to record the behavior of both flies at 100 fps. One camera was positioned to the front and another to the right of the arena. Both cameras were outfitted with lenses (Computar, MLM3X-MP) at ×0.3 Zoom. We illuminated flies using an infrared LED ring light (CCS Inc. LDR2-74IR2-850-LA). Additionally, we placed a UV light source (M375F2, M93L01, and F810SMA-405, Thorlabs, Germany) 5 cm below the front of the arena. This allowed animals to see one another during imaging without PMT bleed through.

The social arena housing the freely moving fly is composed of an assembly of five acrylic pieces. This avoids the use of glue and facilitates cleaning. Each piece is 2 mm thick except for the floor which is 0.75 mm thick. We did a chamfer on the exterior edges of the floor using a conical countersink (DIN 335C-HSS-15mm, NERIOX, Germany). This allowed the spherical treadmill to rise higher and to roll more smoothly. Moreover, we sanded the floor’s exterior to make it translucent. If the floor was fully transparent the freely moving flies would typically freeze, likely due to perception of movements of the foam ball. Finally, a second component consisting of two 4 mm thick acrylic pieces glued together is used to load the freely moving fly into the arena. 2D and 3D CAD models to build or customize this arena are available on Dataverse (See Data availability).

#### Experimental protocol

We recorded three MBON cell types: MBON-β’2mp, MBON-β’2mp-bi, and MBON-calyx. We used ThorImage 4.1 to acquire 352 × 128 pixel images from both the green and red PMT channels. We used ThorSync 4.1 to synchronize imaging data with behavioral data. Pixel sizes were 0.311 µm for MBON-β’2mp, and MBON-β’2mp-bi imaging, and 0.386 µm for MBON-calyx imaging. We recorded 2,400 microscopy frames (around 5 min) for five different experimental epochs: 10 min after the fly was mounted (‘Alone’), when the second fly was placed in the waiting room (‘Before’), when the second fly was introduced into the arena to permit social interactions (‘During’), one hour after interactions began (‘After 1 h’), and two hours after interactions began (‘After 2 h’). We waited 2 min between ‘alone’ and ‘before’ recordings, as well as between ‘before’ and ‘during’ recordings. We selected imaging planes by starting at the ventral-most part of the recorded neurons and then moving between 10–13 µm dorsally where the fluorescence was typically more intense. We readjusted the imaging plane prior to the ‘After 1 h’ and ‘After 2 h’ recordings. Although sometimes the brain did not shift at all or only very little, in some cases it could sink around 7 µm*/*h when the fat bodies and trachea around (not only on top of) the brain were also removed.

#### Behavior analysis

We used two orthogonally positioned IR-sensitive cameras to track the position of the free-moving fly. We trained a different SLEAP (v1.2.8) model for each camera but labeled the same three landmarks: head, thorax, and abdomen **(Supplementary Video 13)**. Both models use a bottom-up approach over a U-Net network. We labeled 150 frames for the front camera and 200 for the side camera. We augmented the training data with rotations (−180^◦^ to 180^◦^), scaling (0.9 to 1.1), Gaussian noise (5 ± 1), contrast (0.5 to 2.0), and brightness (0 to 10). Then, we used the same custom Python code from our behavioral experiments to predict, track, and fill in missing values.

To obtain the position of the tethered fly we first selected a random sample of 50 frames. Then, for each of those frames we binarized the image using Otsu’s method, and applied an opening morphological transformation with a rectangular structuring element of size 50 × 4 for the front camera, and 4 × 70 for the side camera. Next, we obtained the coordinates of the connected component closest to the image’s center. Finally, we calculated the mean position of the fly from all the frames in the random sample and used these coordinates as the origin of the freely moving fly tracking data.

Spherical treadmill rotational speeds were obtained by running FicTrac ^66^ on front camera images recorded at 100 fps. FicTrac was configured as in **Table 4**.

**Table 4:** FicTrac configuration parameters. Columns show parameters and their values as used to calculate ball rotations with FicTrac software.

| Parameter | Value |
| --- | --- |
| Working plane | yz |
| max_bad_frames | 5 |
| opt_bound | 0.35 |
| opt_do_global | yes |
| opt_max_err | 25,000 |
| opt_max_evals | 50 |
| q_factor | 10 |
| thr_ratio | 1.1 |
| thr_win_pc | 0.2 |
| vfov | 3.5 |

#### Functional imaging data post-processing

Even when the proboscis was glued we observed motion artifacts in the brain including large translations and smaller non-affine deformations. Calcium indicators have low baseline fluorescence by design, therefore, it becomes challenging to use them for motion correction. Nevertheless, we could use the signal from the co-expressed red fluorescent protein, tdTomato, to register both the red and green channels. First, we applied a center-of-mass registration to center images. Then, we calculated the motion field from the red channel and applied an optic flow motion correction using bi-linear interpolation as described in ^94^. Finally, we applied a median filter with a kernel size of 3 to remove noise.

#### ΔF/F calculation

Different cells can have different calcium indicator expression levels. Therefore, GCaMP fluorescence is usually analyzed as the change in fluorescence relative to baseline fluorescence. Here we were interested in investigating how social experience might modify the overall activity of individual MBONs. Thus, movements in the z-axis resulting in changes in the imaging plane were problematic. To overcome this issue we used adaptive thresholding with a neighborhood size of 31 pixels to obtain a mask in the red channel. This allowed us to identify MBON neurites in each frame. Then, we applied these image masks to the green channel for every trial and t-stacked the frames for a given epoch. One t-stack from each of the five experimental epochs (alone, before, etc.) was used to calculate the fluorescence baseline as the 5th percentile for each pixel. Pixels outside the mask were assigned their maximum value causing them to have very small values when used to normalize changes in fluorescence. We selected the fluorescence baseline as the 5th percentile of each pixel over all five recording epochs.

With this baseline computed separately for each fly we calculated the ΔF/F as follows: ΔF/F = 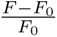; where *F* is the pixel fluorescence intensity at a given frame and *F*_0_ is its corresponding baseline intensity. Finally, we computed the mean ΔF/F for each frame across pixels within the mask. We discarded the first minute of each experiment because fluorescence appeared to be affected by the initiation of laser scanning.

#### Sociability analysis

We used the proxy of spherical treadmill rotations to estimate the tethered fly’s behavior. Thus, it was not possible to differentiate between freezing and relaxed standing. Because of this, we focused our analysis of tethered sociability on the production of fear-like locomotor reactions. First, we defined proximity events when the freely moving fly was less than 6 mm away from the tethered fly. The number of proximity events was comparable for experiments with single- and group-housed flies **(Extended Data Fig. 11a)**. Then, for each fly with more than two proximity events per epoch, we computed the ratio of the median acceleration between the 2.5 s after and 2.5 s before the start of the proximity event. In this case, values higher than 1 would indicate increased acceleration when the freely behaving fly was in close proximity. We observed that, indeed, single-housed flies were more reactive to the free-moving fly in the ‘During’ epoch compared with group-housed animals. Moreover, this reactivity was not present in the ‘After 1 h’ and ‘After 2 h’ epochs in single-housed animals. This metric also remained constant for group-housed animals across epochs **(Extended Data Fig. 11b)**. These data suggest that tethered, dissected animals can also generate fear-like reactions that are suppressed over the course of hours of social exposure.

#### Comparing neural and locomotor dynamics

Spherical treadmill speeds were computed at 100 Hz and neural imaging data at 8 Hz. Therefore, we first interpolated the behavioral data to the lower imaging data frame rate. Then we smoothed all signals (ΔF/F, treadmill speed, and inter-fly distance) using a second-order low-pass Butterworth digital filter with a cutoff frequency of 0.25 Hz. Next, we detected the baselines of these signals using asymmetric least squares fitting with a smoothing parameter of 1*e*7 and a penalizing weighting factor of 0.05 and 0.005 for the treadmill speed and ΔF/F, respectively. We subtracted these baselines from ΔF/F and treadmill speed data. We clipped negative values to 0.

Simply correlating the rotational speed of the spherical treadmill and neural activity could be misleading due to the slow decay of our calcium indicator. Therefore, to quantify neural activity’s association with walking speed, we identified behavioral events in which the fly generated fictive locomotion at greater than 1 mm*/*s for at least 1 s. Events were at least 1.5 s apart. We then set the time window of interest from 1 s before the event began until the end of the event (i.e., when the speed of the treadmill became less than 1 mm*/*s for more than 1.5 s).

We next obtained ΔF/F traces during the same time windows of behavioral events. As a control, for the same number and duration of events we selected a random set of time points and their associated ΔF/F traces. We calculated the mean ΔF/F from real events and compared them to those obtained from random control events. We computed this comparison for sets of random events per condition and calculated the time points where neural activity was significantly different in more than 70% of these random sets. Specifically, *P <* 0.05 for a test with 1,000 permutations with a false discovery rate for positively correlated tests corrected for multiple comparisons. Because the random selection of events may impact the results of this comparison, we obtained 33 sets of random events per fly to compare with the mean signal from real events. For every condition we have three animals. Thus, we ran 99 comparisons per condition. We performed the same analysis to compare the distance between animals with neural activity dynamics. In this case, we defined proximity events when the freely moving fly was less than 6 mm away from the tethered fly for at least 0.5 s. Events were at least 1 s apart. We then set the time window of interest from 1 s before the event began until the end of the event. Then, the selection of random sets and comparisons with neural activity were performed as described above.

#### Analyzing baseline neural activity

To examine the overall baseline activity of MBONs, we obtained segments of the original ΔF/F traces that were equal to or less than the fit by asymmetric least squares smoothing described above. Then, we calculated the mean baseline for each segment detected, and finally, obtained the mean of all mean baselines for each epoch for each fly. As before, we discarded the first minute of every experiment. We performed a similar analysis to calculate each neuron’s average activity, but the segments selected were instead those greater than 0 in the ΔF/F traces where the baseline had been subtracted. We compared different groups using a Kruskal–Wallis test followed by a post hoc Conover’s test. We corrected for multiple comparisons using a false discovery rate for positively correlated tests.

## Data availability

Data are available at: https://dataverse.harvard.edu/dataverse/sociability_learning/. This repository does not include raw behavioral videos due to data storage limits. They are available upon reasonable request.

## Code availability

Analysis code is available at: https://github.com/NeLy-EPFL/Sociability-Learning.git

## Extended data

**Extended Data Fig. 1:**
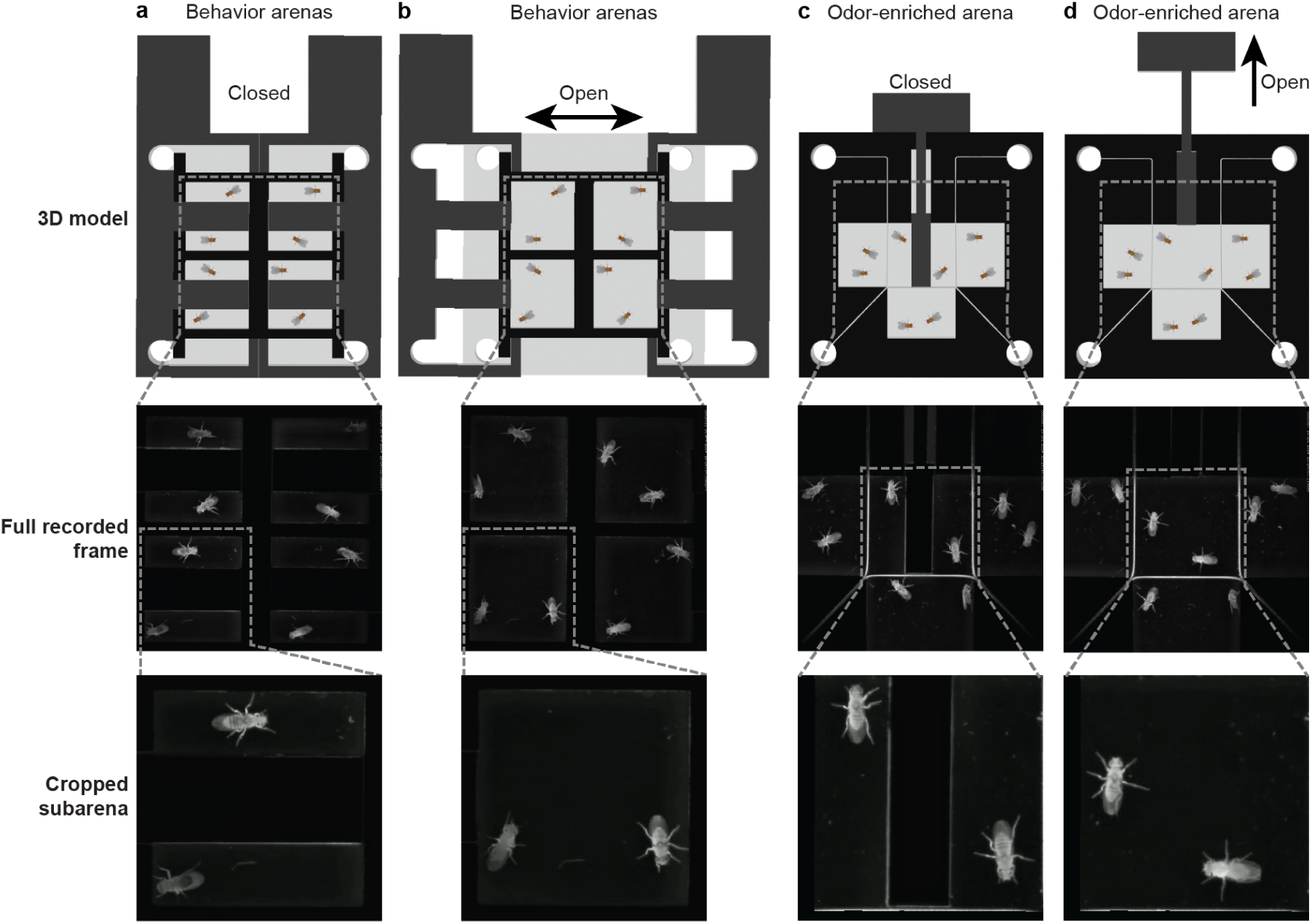
Custom designed arenas for sociability experiments. (a–b) For typical sociability experiments two flies were paired with one another. Then their behavior was tracked and analyzed. To achieve this, we designed an arena with four sub-arenas (top). **(a)** Initially a gate separated the four pairs of animals to prevent interactions. **(b)** These gates could then be manually opened for all four arenas, to enable inter-animal interactions and initiate the experiment. **(c–d)** To test the relative contributions of touch (taste), and smell in sociability learning, we designed another ‘odor-enriched’ arena. This arena has only one pair of experimental flies surrounded by six flies which generate a fly-odor enriched environment. These animals are out of reach due to a mesh wall. **(c)** Initially a gate prevents interactions between the pair of experimental flies. Thus, they can be exposed for hours to fly odors without being given the opportunity to touch or taste another fly. **(d)** Afterwards, the gate is opened to enable inter-animal interactions and initiate the experiment. For all figure panels, the top row shows a 2D projection of a 3D CAD model of the arena, the middle row shows a real recorded camera image, and the bottom row shows a single experimental sub-arena cropped from the original image. This cropped image was used to track and analyze fly behaviors.

**Extended Data Fig. 2:**
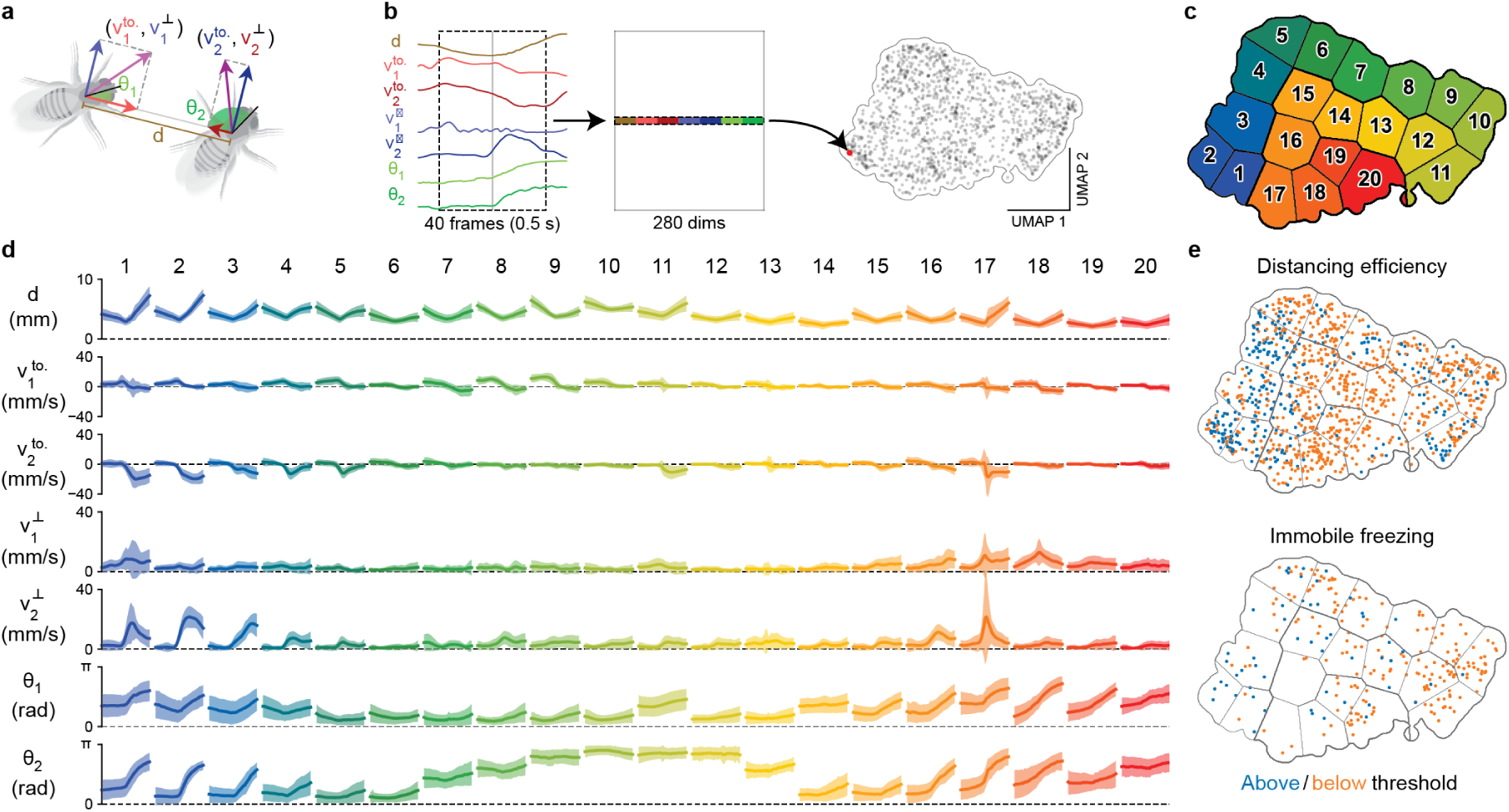
Low-dimensional embedding and clustering of behavioral proximity events. **(a)** Variables used to represent the relative positions and movements of a pair of flies. These include the distance (*d*), velocity components towards and perpendicular to the orientation of the other fly (*v*^to.^*, v*^⊥^), and angles between the headings and the line connecting a pair of flies (*θ*). **(b)** Pipeline for generating a behavioral map of proximity events. For each proximity event, we extracted traces for the variables within a time window of ±0.25 s around the moment when flies are closest to one another. We then flattened these traces into a vector. Vectors from all proximity events were then subjected to nonlinear dimensionality reduction to two dimensions using UMAP. **(c)** The behavioral map was segmented into distinct regions using *k*-means clustering (*k* = 20). Each of the 20 regions represents a different type of proximity event based on the selected variables. **(d)** Time-series traces of the selected variables within the selected time windows for each of the 20 clusters. These traces provide a detailed view of how each variable changes over time within each cluster. Solid lines and shaded regions represent the means and ±1 standard deviations, respectively. **(e)** Proximity events above (blue) and below (orange) thresholds for the distancing efficiency (top), or immobile freezing (bottom) metrics. Data are visualized in a low-dimensional behavioral embedding generated using UMAP. Each dot represents one proximity event. Regions separated by lines are the 20 clusters identified using *k*-means clustering.

**Extended Data Fig. 3:**
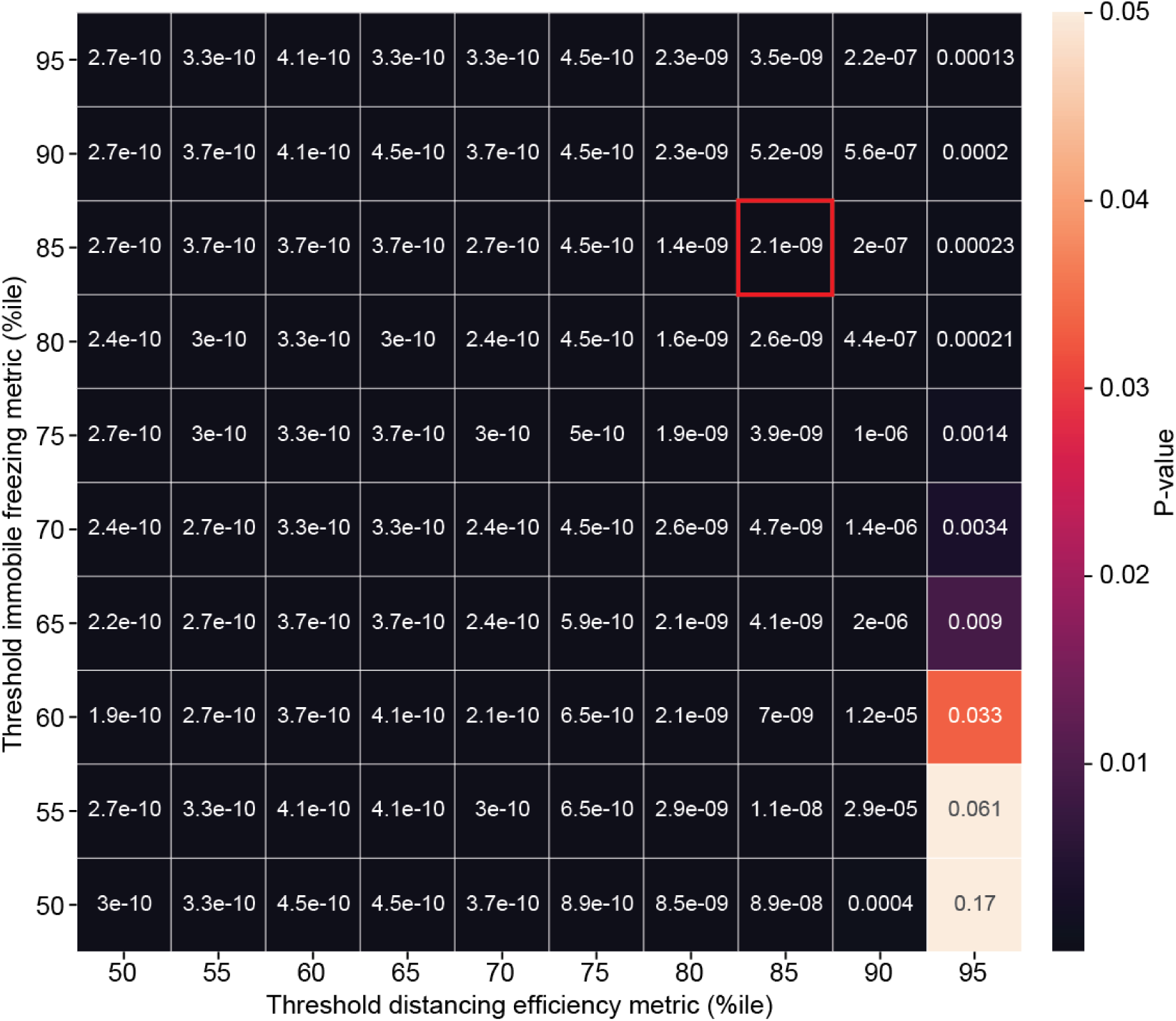
Sensitivity analysis of the thresholds applied to immobile freezing and distancing efficiency data. Shown are *P* values (color-coded) resulting from comparing single and group-housed females in Fig. **1**h. We performed a Mann–Whitney statistical test while varying the thresholds (the 50th to the 95th percentile) applied to distancing efficiency and immobile freezing data. Indicated is the pair of thresholds used throughout the manuscript (square with red borders).

**Extended Data Fig. 4:**
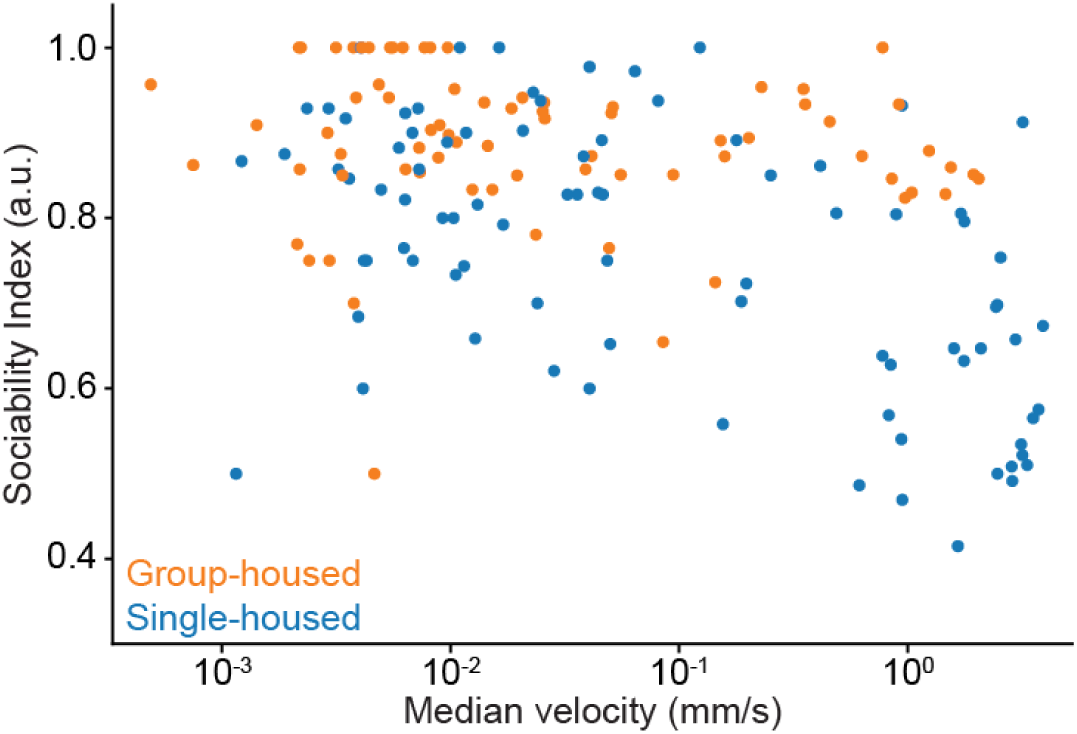
The relationship between an individual’s locomotor activity level and sociability index. Plotted for each single-(blue) or group-housed (orange) animal from female-female and male-male experiments (Fig. **1**h) are their sociability index as a function of median velocity. The Spearman’s correlation coefficient is −0.455. Notably, it is possible for flies to be quite active and yet have a high sociability index (top-right) and for inactive flies to have a low sociability index (bottom-left).

**Extended Data Fig. 5:**
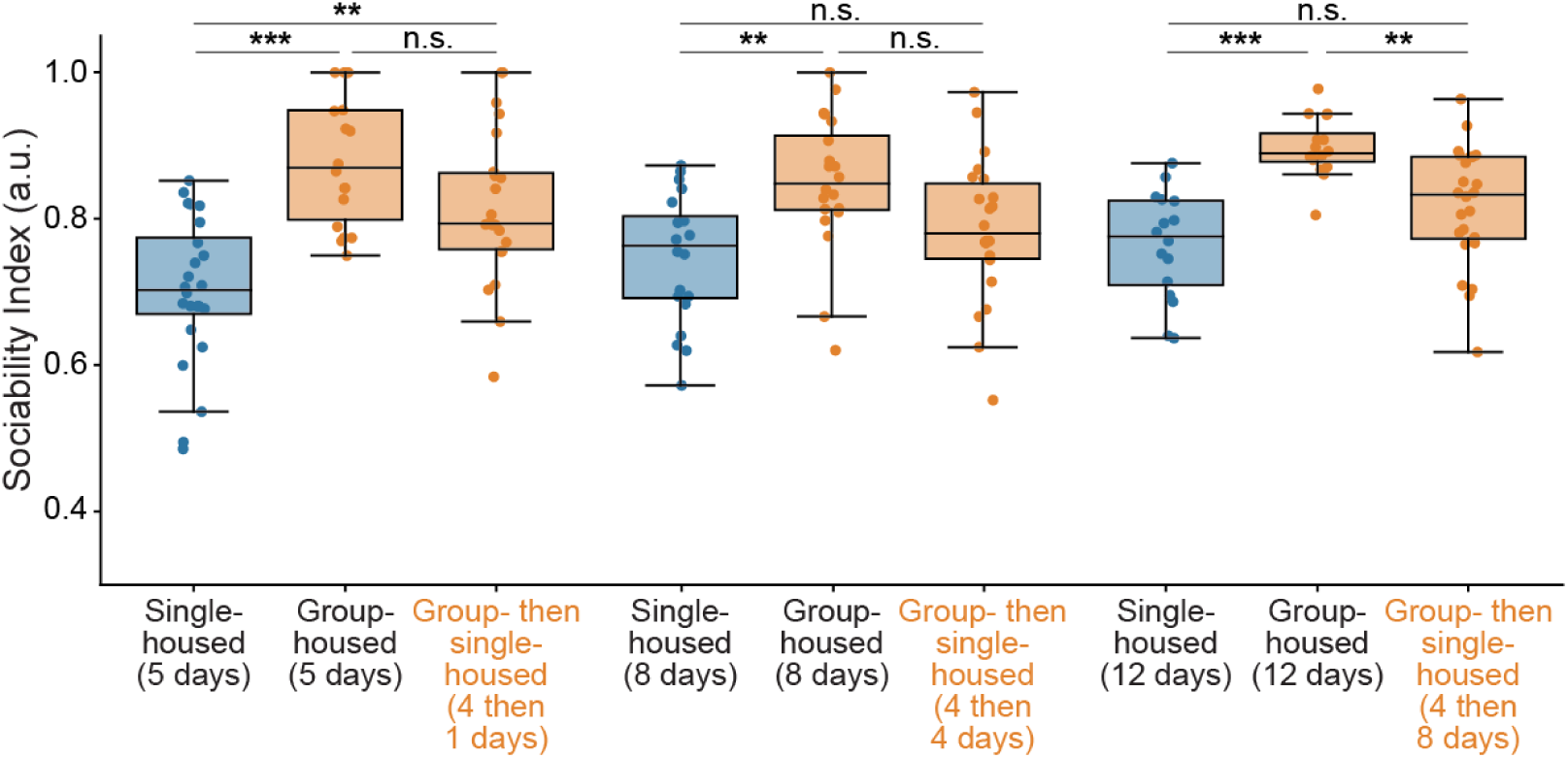
The duration of isolation required to forget sociability. A comparison of sociability across single-housed, group-housed, or group-housed animals that were then isolated for 1 (left), 4 (middle), or 8 (right) days. From left to right, the number of animals analyzed are: *n* = 24, 18, 22, 20, 20, 22, 16, 16, and 24. Flies were compared across conditions to animals of the same age (5, 8, or 12 dpe) using a Kruskal–Wallis test followed by a post hoc Conover’s test with a Holm correction for multiple comparisons. *** *P <* 0.0001, ** *P <* 0.001, n.s. *P* ≥ 0.01.

**Extended Data Fig. 6:**
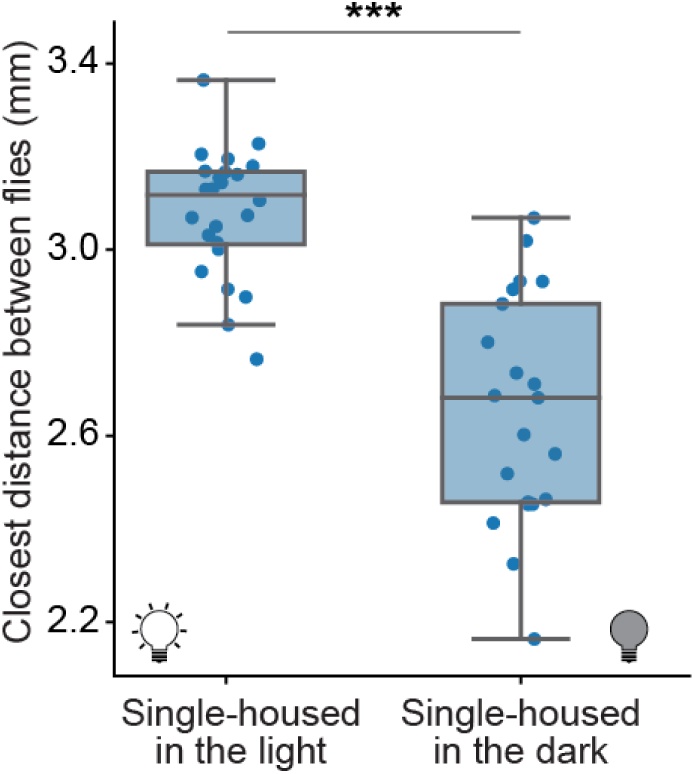
The closest distance between single-housed flies as a function of visual context. The closest distance of single-housed flies during proximity events in the first 30 min of interactions in illuminated arenas (*n* = 24) or in the dark (*n* = 23). Conditions were compared using a Mann–Whitney statistical test. *** *P <* 0.0001.

**Extended Data Fig. 7:**
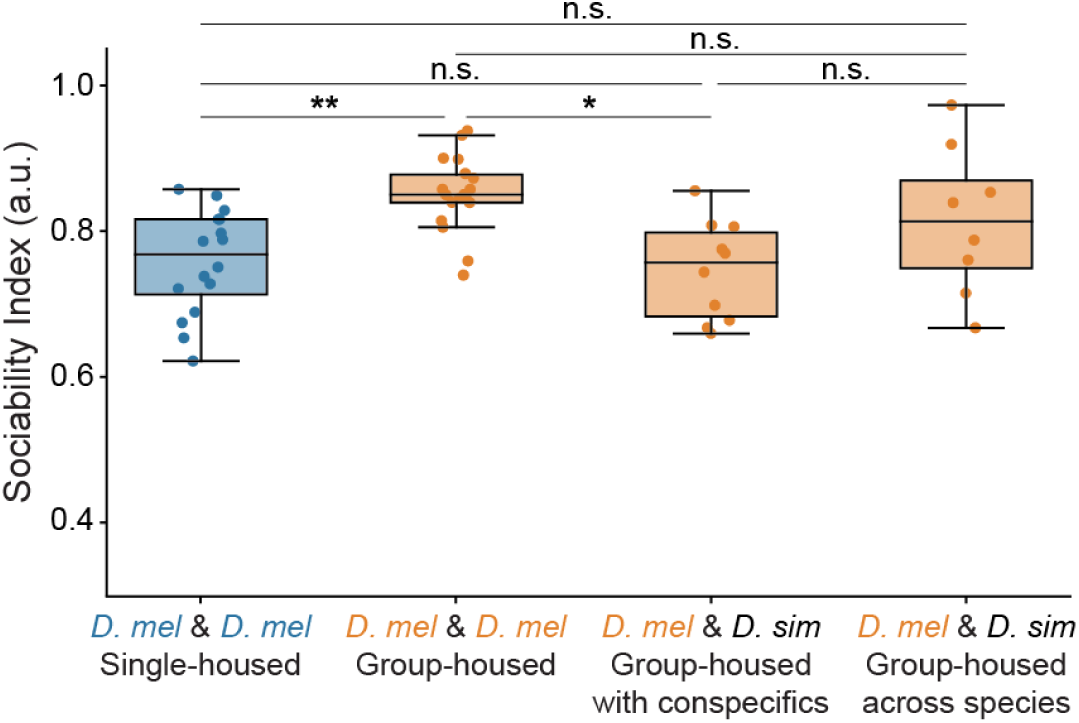
Sociability learning between *Drosophila* species. We assessed whether odors from *Drosophila simulans* could drive sociability learning in *Drosophila melanogaster*. As a baseline, we compared either single-housed *D. mel* (*n* = 16) or group-housed *D. mel* (*n* = 18). We also examined the sociability index of *D. mel* towards *D. sim.* when they were either raised housed with conspecifics only (*n* = 10) or mixed together (*n* = 8). Data points are from *D. mel* behavioral events only. For statistical comparisons, we used a Kruskal–Wallis test followed by a post hoc Conover’s test with a Holm correction for multiple comparisons. ** *P <* 0.001, * *P <* 0.01, n.s. *P* ≥ 0.01.

**Extended Data Fig. 8:**
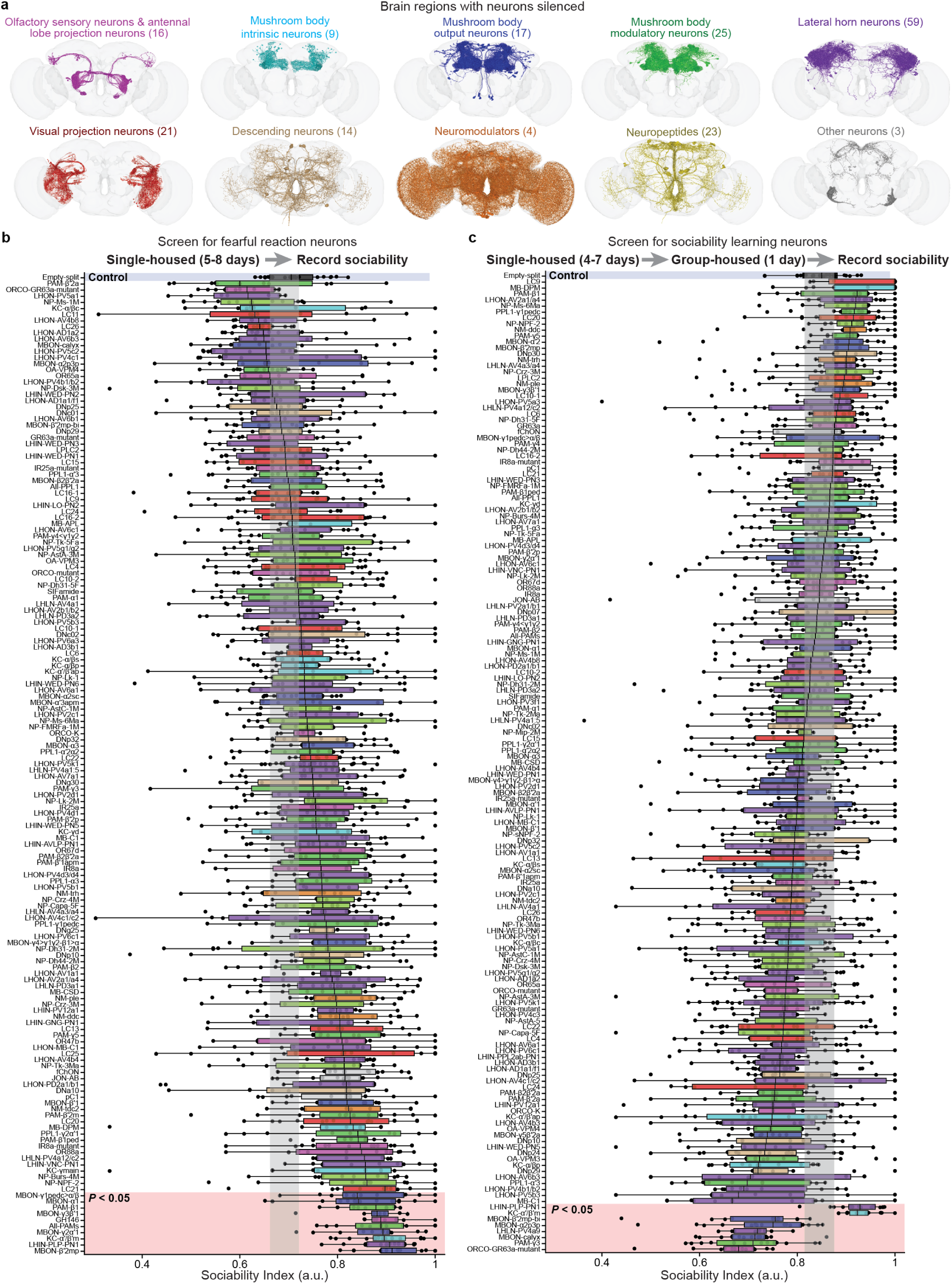
Neural silencing screens for fearful reactions and sociability learning. **(a)** Fly brain connectome renderings of up to 100 neurons in brain regions selected for neural silencing. Neurons were targeted for silencing in the olfactory system (peripheral olfactory sensory neurons, antennal lobe projection neurons, mushroom body intrinsic neurons, mushroom body output neurons, mushroom body modulatory neurons, and lateral horn neurons). As well, neurons were targeted in the visual, descending, neuromodulatory, neuropeptide, and other systems. **(b–c)** We performed two complementary neural silencing screens. **(b)** In the first screen, we tested the roles of neurons in the production of fearful reactions. We single-housed flies and then recorded their pairwise sociability. **(c)** In the second screen, we tested the roles of neurons in the learning of sociability by initially single housing flies and then group housing them for 1 day. We then recorded their pairwise sociability. Shown are the sociability indices for 159 and 163 of the driver lines screened for the reaction and learning screens, respectively (some lines had too few proximity events). The number of flies tested are enumerated in Table 1. Driver lines are color-coded by their respective brain region as in panel **a**. The vertical gray region indicates the 95% bootstrap confidence interval for the control flies (empty-split-Gal4, blue transparency) acquired from 1,000 bootstrap samples. Driver lines that are statistically significantly different from the control (*P <* 0.05) are highlighted (pink transparency). A Mann–Whitney statistical test was performed comparing the control line with every tested line, the Holm method was used to correct for multiple comparisons.

**Extended Data Fig. 9:**
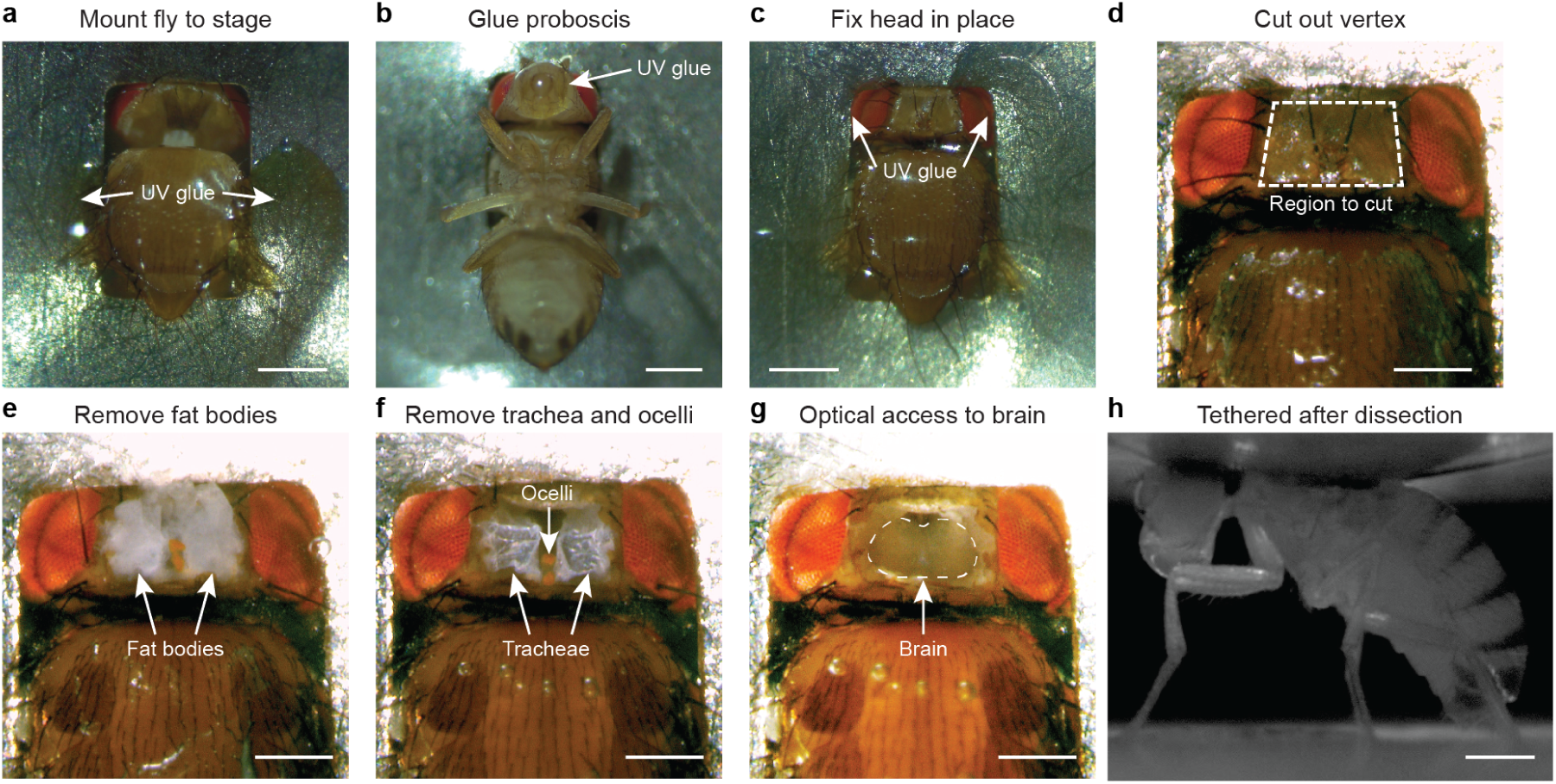
Dissection approach used to record MBON activity during sociability learning. **(a)** A fly is mounted onto a behavioral stage and its thorax is glued in place using UV curable glue. Scale bars are 0.3 mm for panels **(a-c)**. **(b)** The proboscis and maxillary palps are glued in place. **(c)** The head is then glued to the shim. **(d)** The dashed lines indicate where the dorsal head will be cut to remove the vertex. Scale bars are 0.15 mm for panels **(d-g)**. **(e)** After the vertex cuticle is pulled away, fat bodies covering the brain are then also removed. **(f)** Subsequently, the trachea and ocelli are removed. **(g)** We then have optical access to the brain. Dashed lines show the contour of the central brain. **(h)** Fly mounted under the two-photon microscope on a spherical treadmill within the social arena. Scale bar is 0.5 mm.

**Extended Data Fig. 10:**
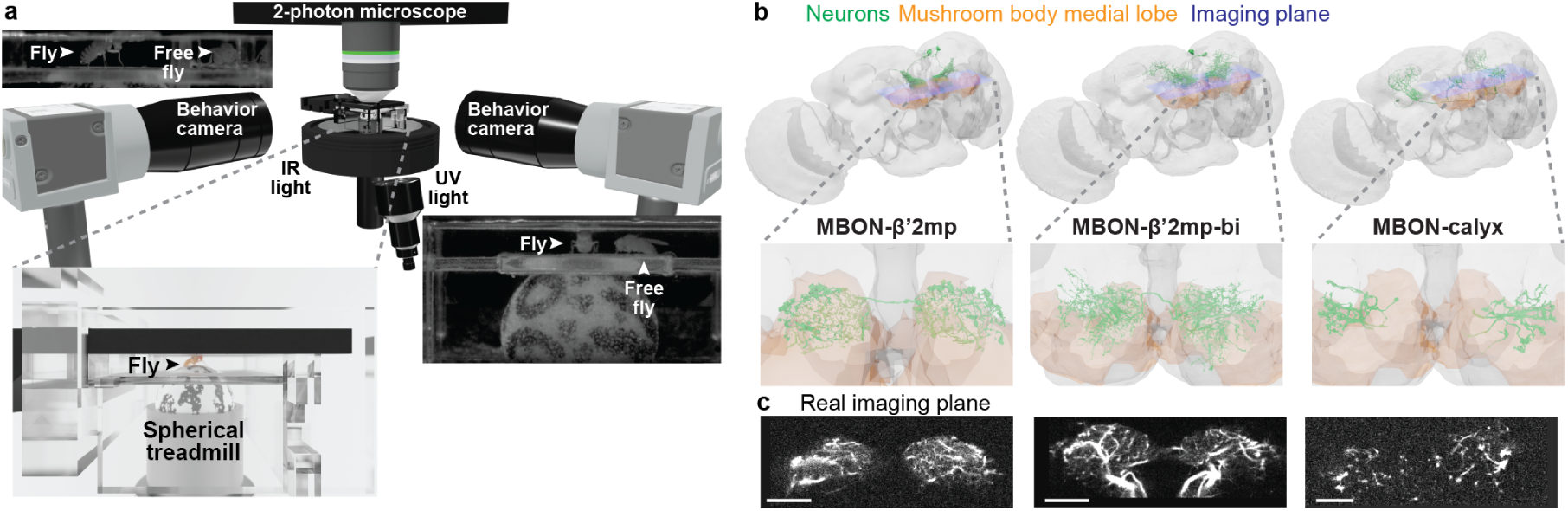
Two-photon microscope system used to record MBON activity during sociability learning. **(a)** Experimental system used to record brain activity during inter-fly social interactions. A two-photon microscope records MBON activity in a tethered animal walking on a spherical treadmill whose rotations are quantified using a front-facing camera. Simultaneously, the movements of a freely-behaving fly are tracked. A UV LED is used to illuminate the arena. **(b–c)** Brain imaging during social interactions. **(b)** A connectome-based rendering of the brain. The mushroom body’s medial lobe is shown for reference (orange). The morphologies of three selected MBONs—MBON-β’2mp (left), MBON-β’2mp-bi (middle), and MBON-calyx (right)—are also shown (green). Below is a cross-section of the rendering corresponding to the imaging plane as well as **(c)** recordings of this imaging plane from real neural recordings in which MBONs express the calcium indicator, GCaMP6s. Scale bars are 20 µm.

**Extended Data Fig. 11:**
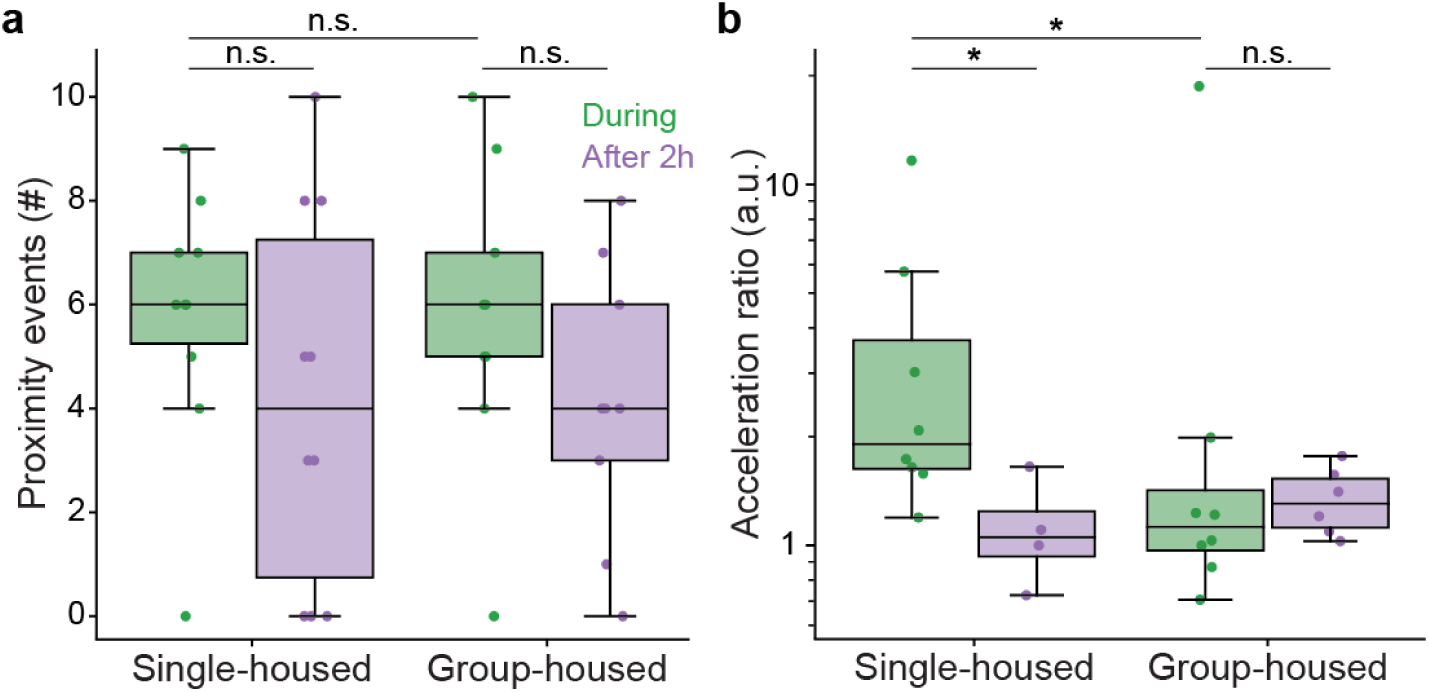
Proximity events and fear-like reactions in tethered flies. **(a)** The number of proximity events remains relatively constant throughout the duration of an experiment. This is true regardless of the social experience of the tethered animal and the time since the experiment began (n=9 for all conditions). **(b)** Single-housed tethered animals show larger increases in acceleration at the onset of proximity events than group-housed animals. This is reduced after 2 h of social exposure although it remains constant for group-housed flies. The acceleration ratio was calculated for flies with more than two proximity events per epoch (single-housed ‘During’, n=8 flies; single-housed ‘After 2h’, n=4 flies; group-housed ‘During’, n=9 flies; group-housed ‘After 2h’, n=7 flies. We performed pairwise comparisons using the Mann–Whitney statistical test and corrected for multiple comparisons with the false discovery rate for positively correlated tests. * *P <* 0.05, n.s. *P* ≥ 0.05.

**Extended Data Fig. 12:**
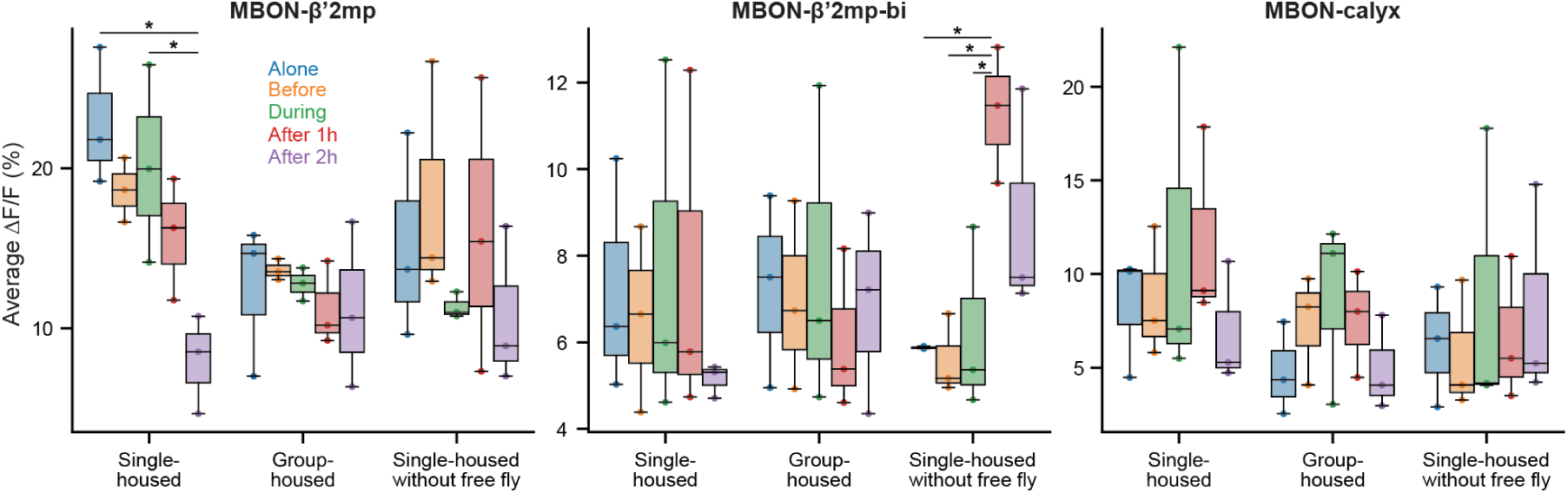
MBONs mean neural activity during behavior. Mean neural activity during behavior (i.e., after removing the baseline) across epochs and social conditions for MBON-β’2mp (left), MBON-β’2mp-bi (middle), and MBON-calyx (right). Conditions were compared using a Kruskal–Wallis test followed by a Conover’s post hoc test corrected for multiple comparisons with the false discovery rate for positively correlated tests. * *P <* 0.05, n.s. *P* ≥ 0.05.

## Supplementary Information

**Supplementary Information File - Neural activity traces for all MBONs across all conditions.**

[Link to Supplementary Information File 1](https://www.dropbox.com/scl/fi/o1wk3zhakw3sg31cjms3d/Lobato_SociabilityLearning_SupplementaryInformationFile2024.pdf?rlkey=028p0kmv1ri5o1nt05vdfc7hn&dl=0)

## Supplementary Videos

**Video 1: Examples of fearful reactions between single-housed animals.** Shown sequentially are four examples of monitoring behaviors, repulsion, and freezing of one fly (cyan trajectories) in response to a second fly (magenta trajectories). Indicated are proximity events (red circles).

[Link to Supplementary Video 1](https://www.dropbox.com/scl/fi/bpltf7v1pm46ipu05rreb/Video1.mp4?rlkey=q6wz75ots3lyen5n9vnybcs7l&dl=0)

**Video 2: Examples of sociable behaviors between group-housed animals.** Shown sequentially are four examples each of walking alongside one another, interactions via leg contact, and undisturbed reactions of one fly (cyan trajectories) in the presence of a second fly (magenta trajectories). Indicated are proximity events (red circles).

[Link to Supplementary Video 2](https://www.dropbox.com/scl/fi/3yy9aur72cz3hzld53oab/Video2.mp4?rlkey=wcatl7leaq8l7vje64mj241jx&dl=0)

**Video 3: Examples of behaviors from embedding clusters of proximity events.** Shown in sequence are 35 randomly sampled examples of behaviors from each cluster (highlighted in white in the 2D UMAP space, top-left). Examples taken from single- or group-housed animals are outlined in blue and orange, respectively. The ratio of single and group-housed flies reflects the cluster’s composition as closely as possible. Events are shown within a 0.5 s time window centered on the moment when the flies are closest to one another. In each example, a single fly (cyan trajectories) reacts to a second fly (magenta trajectories). Shown as well (bottom-left) are traces for key variables: distance (*d*), velocity components (*v*^to.^*, v*^⊥^), and relative headings (|*θ*|). Indicated are the mean trace (solid line) and individual event traces (translucent lines) for each cluster.

[Link to Supplementary Video 3](https://www.dropbox.com/scl/fi/5uhs6y1vg78h90hpbudyp/Video3.mp4?rlkey=nyekumz38g4qh3eob0omxl08h&dl=0)

**Video 4: Examples of behaviors across a range of distancing efficiency values.** Eight examples of proximity events from high (top-left) to low (bottom-right) distancing efficiency values. Indicated on the top-right is the distancing efficiency value for the fly in question (cyan trajectory). Those in the top and bottom rows are above and below the control threshold, respectively.

[Link to Supplementary Video 4](https://www.dropbox.com/scl/fi/1ph6nie3yx00wiql1lo9g/Video4.mp4?rlkey=5dvrx7gs451pbg6s0l4se4yjw&dl=0)

**Video 5: Examples of behaviors across a range of immobile freezing values.** Eight examples of proximity events from high (top-left) to low (bottom-right) immobile freezing values. Indicated on the top-right is the immobile freezing value for the fly in question (cyan trajectory). Those in the top and bottom rows are above and below the control threshold, respectively.

[Link to Supplementary Video 5](https://www.dropbox.com/scl/fi/caojm9lyz9fmks1bpb0jn/Video5.mp4?rlkey=dm3glyvuraeoah7tfhou9aad5&dl=0)

**Video 6: Examples of inter-fly interactions in the dark.** Shown are four proximity events for single-housed (left) and four events for group-housed flies (right) in complete darkness.

[Link to Supplementary Video 6](https://www.dropbox.com/scl/fi/0iuam1ze6fakxueoaj86m/Video6.mp4?rlkey=l9zsgrcpy00vxfh9gt2pa8amf&dl=0)

**Video 7: Example of a pair of single-housed flies in the fly-odor enriched arena.** The behavior of a pair of flies (center) just after the gate separating them, which had been closed for 2 h, is opened. The central chamber is separated from three adjacent chambers by an odor permeable mesh. Each of the adjacent chambers contains another pair of flies.

[Link to Supplementary Video 7](https://www.dropbox.com/scl/fi/0e8b8cuk094py4tbbildr/Video7.mp4?rlkey=8p31c0aeqiafjcw8acw51jx95&dl=0)

**Video 8: Examples of fly-beetle interactions.** Shown are four sets of interactions between a fly and beetle that were (left) raised only with their own species or (right) raised mixed together.

[Link to Supplementary Video 8](https://www.dropbox.com/scl/fi/4bfxmzpha7t79zasm0uxx/Video8.mp4?rlkey=y1p62jon7zv0j4qqf67w5u560&dl=0)

**Video 9: MBON-β’2mp reaction and learning screen behaviors.** The behavior of eight pairs of flies with MBON-β’2mp silenced. Shown are behaviors from the fearful reaction (left) and sociability learning (right) screens. Indicated are each fly’s trajectories (cyan and magenta).

[Link to Supplementary Video 9](https://www.dropbox.com/scl/fi/yqmvr6lwt4b09zeg5wq07/Video9.mp4?rlkey=tm6w0xs6zmft6rh1yc3226tai&dl=0)

**Video 10: MBON-β’2mp-bi reaction and learning screen behaviors.** The behavior of eight pairs of flies with MBON-β’2mp-bi silenced. Shown are behaviors from the fearful reaction (left) and sociability learning (right) screens. Indicated are each fly’s trajectories (cyan and magenta).

[Link to Supplementary Video 10](https://www.dropbox.com/scl/fi/uyxeezeq0ptjciz3q3vhs/Video10.mp4?rlkey=0y5xggkvucsqeaxxao5zb3h4w&dl=0)

**Video 11: MBON-calyx reaction and learning screen behaviors.** The behavior of eight pairs of flies with MBON-calyx silenced. Shown are behaviors from the fearful reaction (left) and sociability learning (right) screens. Indicated are each fly’s trajectories (cyan and magenta).

[Link to Supplementary Video 11](https://www.dropbox.com/scl/fi/icplnadzuf1481fsz3x5g/Video11.mp4?rlkey=7t6h6zp1dhm28puwmay5aoz3r&dl=0)

**Video 12: Animation of the system used for two-photon microscopy during social interactions.** Flies have their wings removed and are then mounted onto a custom acrylic-based social arena. The fly is positioned above an air-suspended spherical treadmill. Prior to this, the cuticle, fat bodies, and trachea above the brain were removed to gain optical access to the mushroom body. This arena design enables a second, freely behaving fly to enter and explore the arena. Outside the arena, two front and right facing cameras record the behavior of the freely moving fly and rotations of the spherical treadmill. The arena is illuminated by 375 nm UV light that allows flies to see one another and 850 nm LEDs that enable near-infrared camera recordings. Simultaneously, a microscope objective is placed above the fly brain to enable two-photon calcium imaging of MBON activity.

[Link to Supplementary Video 12](https://www.dropbox.com/scl/fi/00wm6rg1b9aq9w3atail2/Video12.mp4?rlkey=5rmhgi288bm733vefnclvn5pl&dl=0)

**Video 13: Example data from MBON-β’2mp functional imaging.** Shown are views from behavioral cameras placed to the right and in front of the tethered animal during neural recordings (top-left). Tracking (blue lines) of the freely moving fly is overlaid in both camera images. These tracking data are then used to reconstruct the relative birds-eye position of the freely behaving fly with respect to the tethered fly (top-right). Shown as well (bottom-left) are registered GCaMP6s (cyan) and tdTomato (red) fluorescence imaging data from MBON-β’2mp neurons in the ‘During’ epoch of the experiment. These data were processed (middle-left) to generate a pixel-wise ΔF/F image in which neurons range from being silent (blue) to being active (red).

[Link to Supplementary Video 13](https://www.dropbox.com/scl/fi/q96r73xlif9agptdgkzpl/Video13.mp4?rlkey=pf270r7qucbon8akqscuuh2t3&dl=0)

## Acknowledgments

We thank S. Boy-Röttger for assistance in animal preparation; M. Durrieu for valuable insights; R. Benton for sharing olfactory mutant *D. mel.* and wild-type *D. sim.*; the members of the Neuroengineering Laboratory for helpful discussions and comments on an earlier version of the manuscript. Stocks obtained from the Bloomington Drosophila Stock Center (NIH P40OD018537) were used in this study. PR acknowledges support from an SNSF Project Grant (175667), and an SNSF Eccellenza Grant (181239). VLR acknowledges support from the Mexican National Council for Science and Technology, CONACYT, under the grant number 709993. TKCL acknowledges support from the Croucher Foundation.

## Author Contributions

V.L.R. - Conceptualization, Methodology, Software, Validation, Formal Analysis, Investigation, Data Curation, Validation, Writing – Original Draft Preparation, Writing – Review & Editing, Visualization.

T.K.C.L. - Methodology, Software, Validation, Formal Analysis, Investigation, Validation, Writing – Original Draft Preparation, Writing – Review & Editing, Visualization.

P.R. - Conceptualization, Methodology, Resources, Writing – Original Draft Preparation, Writing - Review & Editing, Supervision, Project Administration, Funding Acquisition.

## Ethical compliance

All experiments were performed in compliance with relevant national (Switzerland) and institutional (EPFL) ethical regulations.

