## Supplementary Information File for "Conspecific sociability is regulated by associative learning circuits"

Victor Lobato-Rios[1], Thomas Ka Chung Lam[1], and Pavan Ramdya*[1]

[1]Neuroengineering Laboratory, Brain Mind Institute & Interfaculty Institute of Bioengineering, EPFL, Lausanne, Switzerland

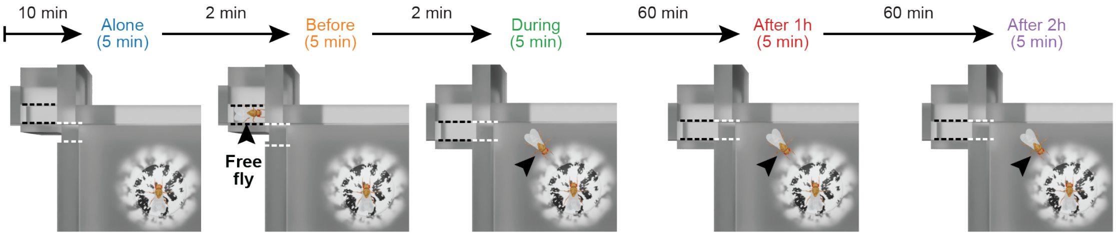


**Supp. File Fig.** 1: **Experimental protocol used to assess the degree to which MBON activities reflect the presence or position of another freely behaving fly, locomotor reactions of the tethered fly, or long-term sociability learning.** Experimental protocol used to assess the degree to which MBON activities reflect the presence or position of another freely behaving fly, locomotor reactions of the tethered fly, or long-term sociability learning. After flies habituated on the spherical treadmill for 10 minutes, MBONs were recorded for five minutes to capture their spontaneous activity in the absence of another fly ('Alone'). After 2 min, a second fly (black arrowhead) was placed into a waiting room (black dashed lines), out of sight but potentially detectable through olfaction ('Before'). Then, 2 min later, this second fly is allowed to enter the arena through an entrance (white dashed lines) and to interact with the tethered animal ('During'). The fourth and fifth epochs of 5 min recordings are one hour ('After 1 h') and two hours ('After 2 h') after the second fly was introduced into the arena. This allowed us to identify potential correlates of sociability learning.

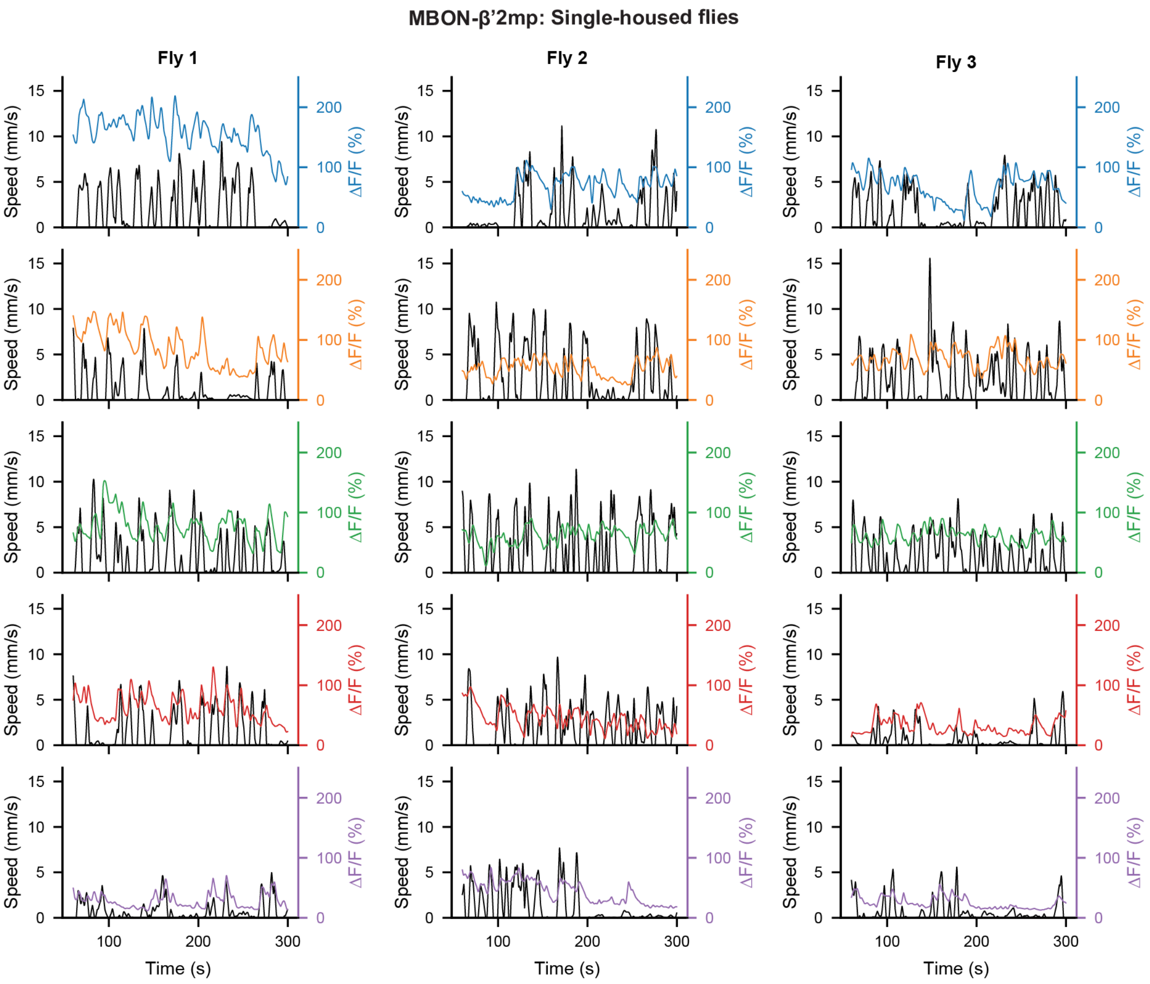


**Supp. File Fig.** 2: **MBON-$\beta$’2mp: Single-housed flies neural activity and spherical treadmill speed.** Treadmill-derived locomotor speed (black) and ΔF/F neural activity traces for each single-housed fly recorded across the five experimental epochs (color-coded as in **Supp. File Fig.** 1).

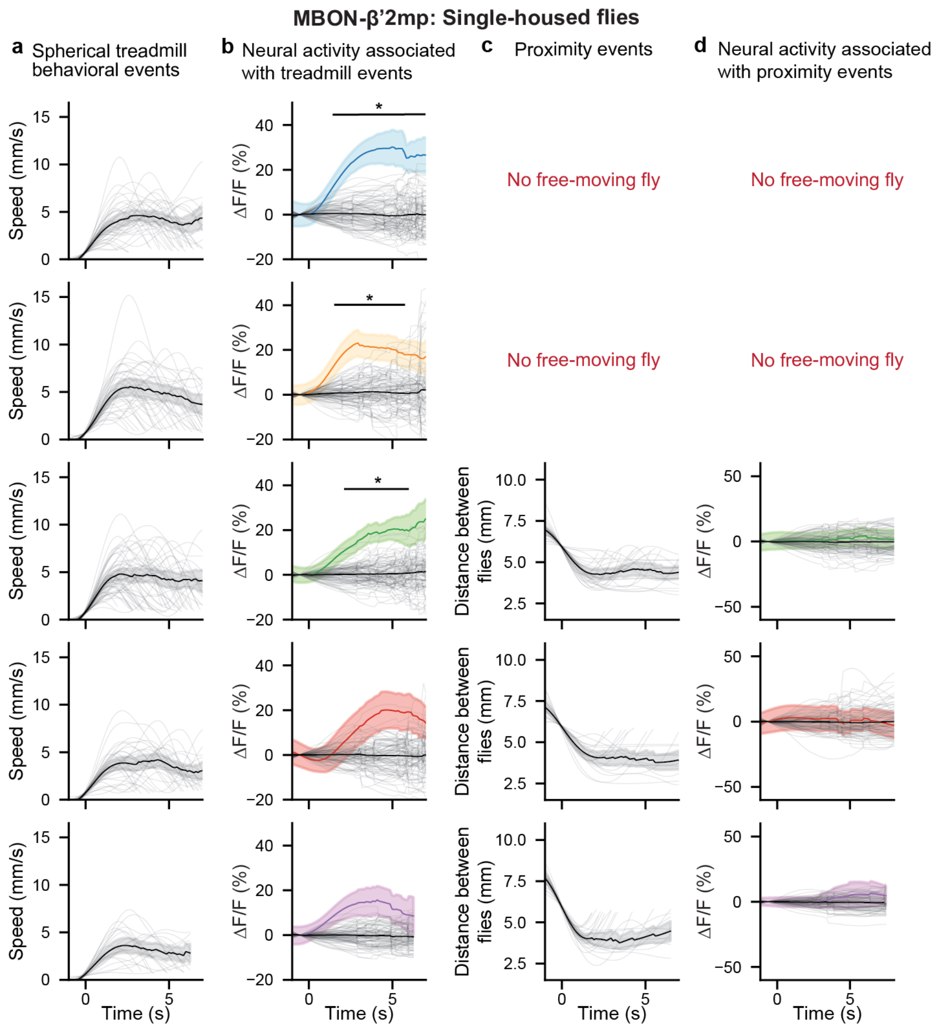


**Supp. File Fig.** 3: **MBON-$\beta$'2mp: Single-housed flies spherical treadmill speed, and proximity events.** **(a)** Behavioral events were selected for each epoch by thresholding treadmill speed and time-locked to event onset (0 s). Overlaid are individual events (translucent traces) and the mean of all events for all three animals for a given epoch (solid black traces) along with the 95% confidence interval (gray transparency). **(b)** Mean neural activity and 95% confidence interval associated with the spherical treadmill events (traces color-coded by experimental epoch **Supp. File Fig.** 1). Overlaid are traces (translucent gray) representing the mean activity of each set of randomly selected events. The means and confidence intervals of all randomly selected events are also shown for comparison (black traces). Mean traces were compared at each time point using a permutation test followed by a correction for multiple comparisons using the false discovery rate for positively correlated tests. * $P < 0.05$. All other comparisons were statistically similar to one another (i.e., $P \geq 0.05$). **(c)** Behavioral events were selected for each epoch by thresholding the distance between flies and time-locked to event onset (0 s). Overlaid are individual events (translucent traces) and the mean of all events for all three animals for a given epoch (solid black traces) along with the 95% confidence interval (gray transparency). **(d)** Mean neural activity and 95% confidence interval associated with the proximity events (traces color-coded by experimental epoch). Overlaid are traces (translucent gray) representing the mean activity of each set of randomly selected events. The means and confidence intervals of all randomly selected events are also shown for comparison (black traces). Mean traces were compared at each time point using a permutation test followed by a correction for multiple comparisons using the false discovery rate for positively correlated tests. All comparisons were statistically similar to one another (i.e., $P \geq 0.05$).

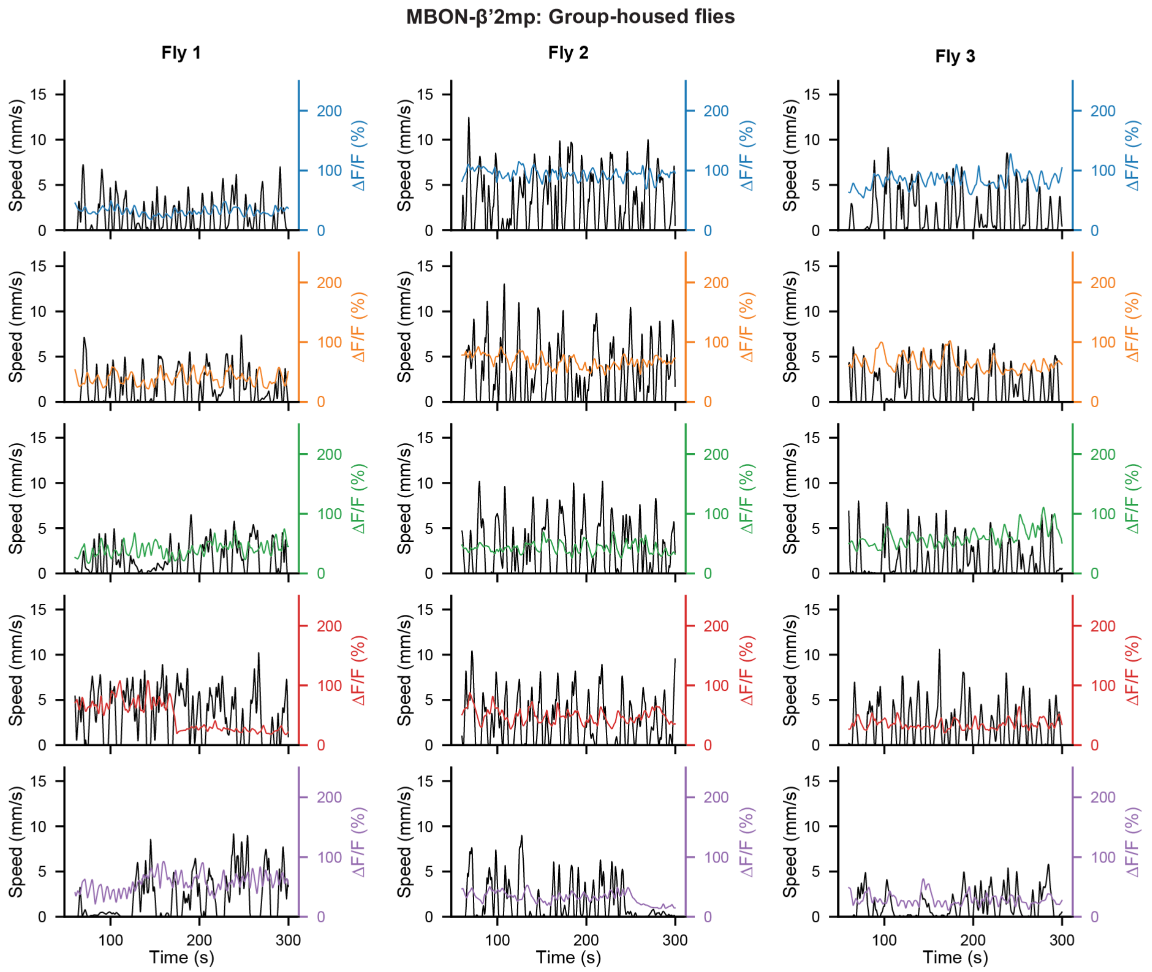


**Supp. File Fig.** 4: **MBON-$\beta$'2mp: Group-housed flies neural activity and spherical treadmill speed.** Treadmill-derived locomotor speed (black) and ΔF/F neural activity traces for each group-housed fly recorded across the five experimental epochs (color-coded as in **Supp. File Fig.** 1).

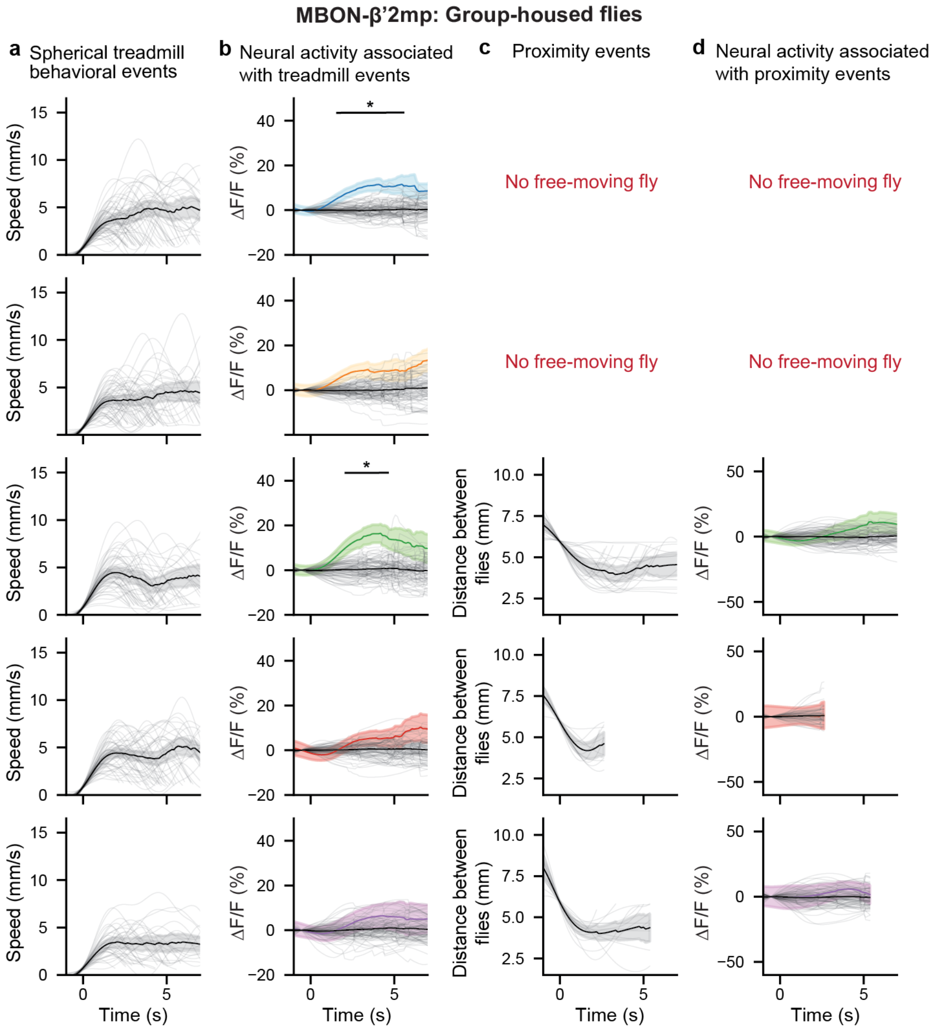


**Supp. File Fig.** 5: **MBON-$\beta$'2mp: Group-housed flies spherical treadmill speed, and proximity events.** **(a)** Behavioral events were selected for each epoch by thresholding treadmill speed and time-locked to event onset (0 s). Overlaid are individual events (translucent traces) and the mean of all events for all three animals for a given epoch (solid black traces) along with the 95% confidence interval (gray transparency). **(b)** Mean neural activity and 95% confidence interval associated with the spherical treadmill events (traces color-coded by experimental epoch **Supp. File Fig.** 1). Overlaid are traces (translucent gray) representing the mean activity of each set of randomly selected events. The means and confidence intervals of all randomly selected events are also shown for comparison (black traces). Mean traces were compared at each time point using a permutation test followed by a correction for multiple comparisons using the false discovery rate for positively correlated tests. * $P < 0.05$. All other comparisons were statistically similar to one another (i.e., $P \geq 0.05$). **(c)** Behavioral events were selected for each epoch by thresholding the distance between flies and time-locked to event onset (0 s). Overlaid are individual events (translucent traces) and the mean of all events for all three animals for a given epoch (solid black traces) along with the 95% confidence interval (gray transparency). **(d)** Mean neural activity and 95% confidence interval associated with the proximity events (traces color-coded by experimental epoch). Overlaid are traces (translucent gray) representing the mean activity of each set of randomly selected events. The means and confidence intervals of all randomly selected events are also shown for comparison (black traces). Mean traces were compared at each time point using a permutation test followed by a correction for multiple comparisons using the false discovery rate for positively correlated tests. All comparisons were statistically similar to one another (i.e., $P \geq 0.05$).

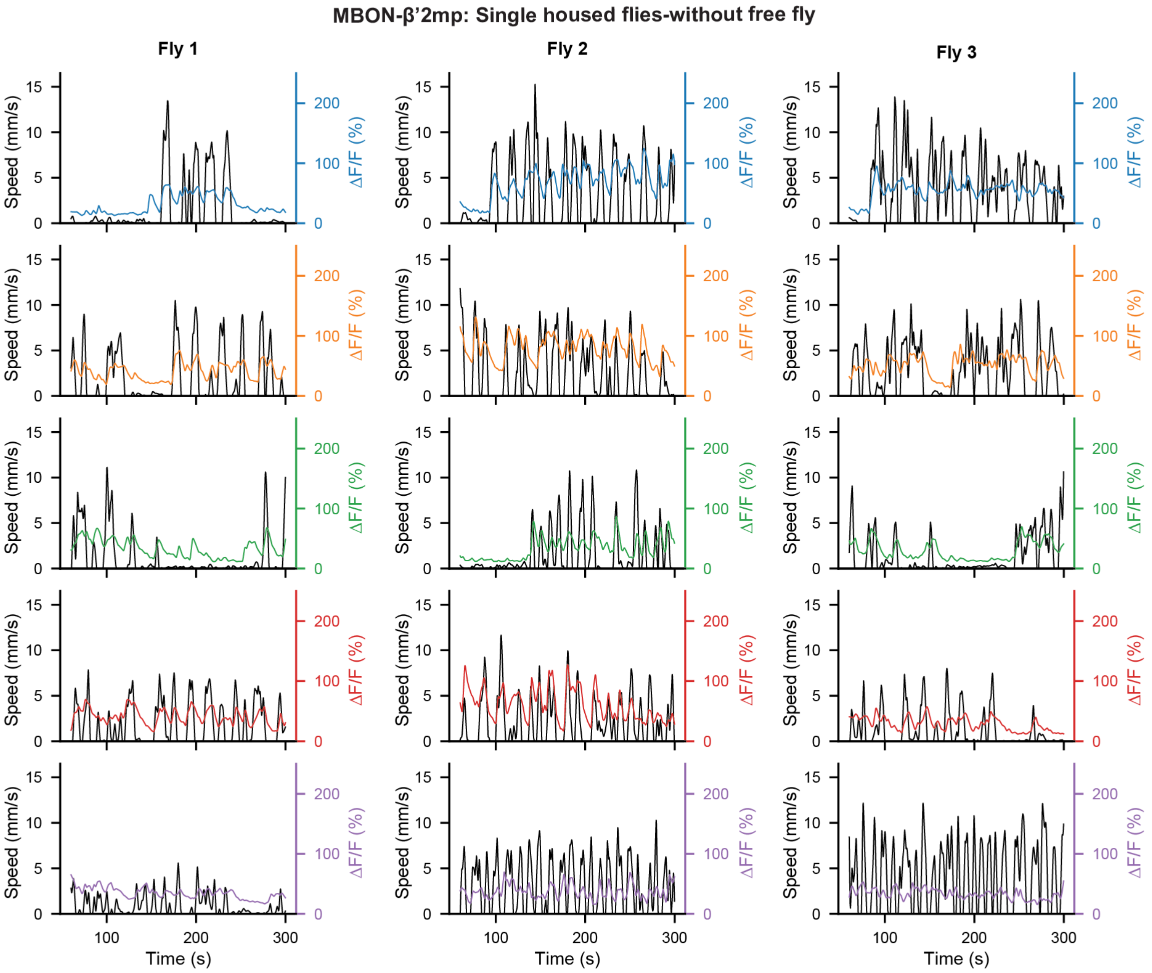


**Supp. File Fig.** 6: **MBON-$\beta$'2mp: Single-housed flies neural activity and spherical treadmill speed without a freely-moving fly in the arena.** Treadmill-derived locomotor speed (black) and ΔF/F neural activity traces for each single-housed fly recorded across the five experimental epochs (color-coded as in **Supp. File Fig.** 1). There was no freely-moving fly during the experiment.

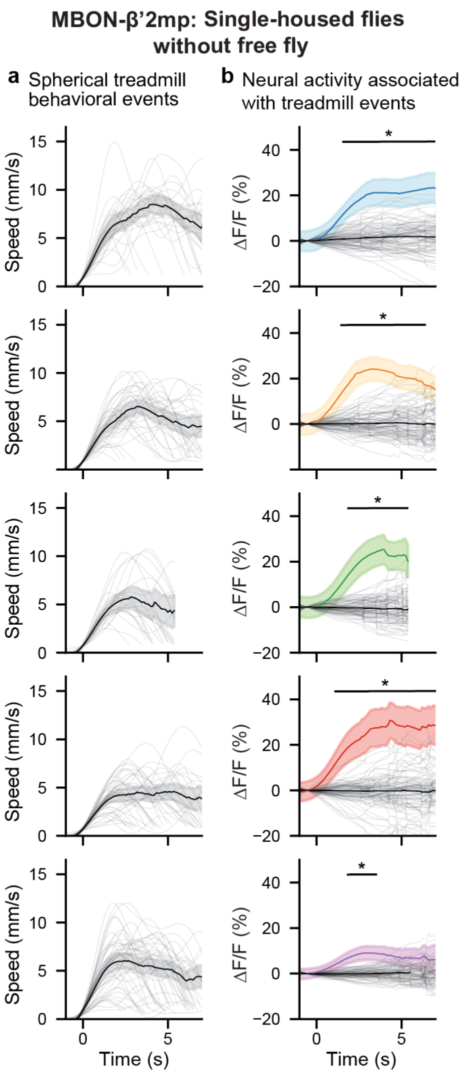


**Supp. File Fig.** 7: **MBON-$\beta$'2mp: Single-housed flies spherical treadmill speed and proximity events without a freely moving fly in the arena. (a)** Behavioral events were selected for each epoch by thresholding treadmill speed and time-locked to event onset (0 s). Overlaid are individual events (translucent traces) and the mean of all events for all three animals for a given epoch (solid black traces) along with the 95% confidence interval (gray transparency). **(b)** Mean neural activity and 95% confidence interval associated with the spherical treadmill events (traces color-coded by experimental epoch **Supp. File Fig.** 1). Overlaid are traces (translucent gray) representing the mean activity of each set of randomly selected events. The means and confidence intervals of all randomly selected events are also shown for comparison (black traces). Mean traces were compared at each time point using a permutation test followed by a correction for multiple comparisons using the false discovery rate for positively correlated tests. * $P < 0.05$. All other comparisons were statistically similar to one another (i.e., $P \geq 0.05$).

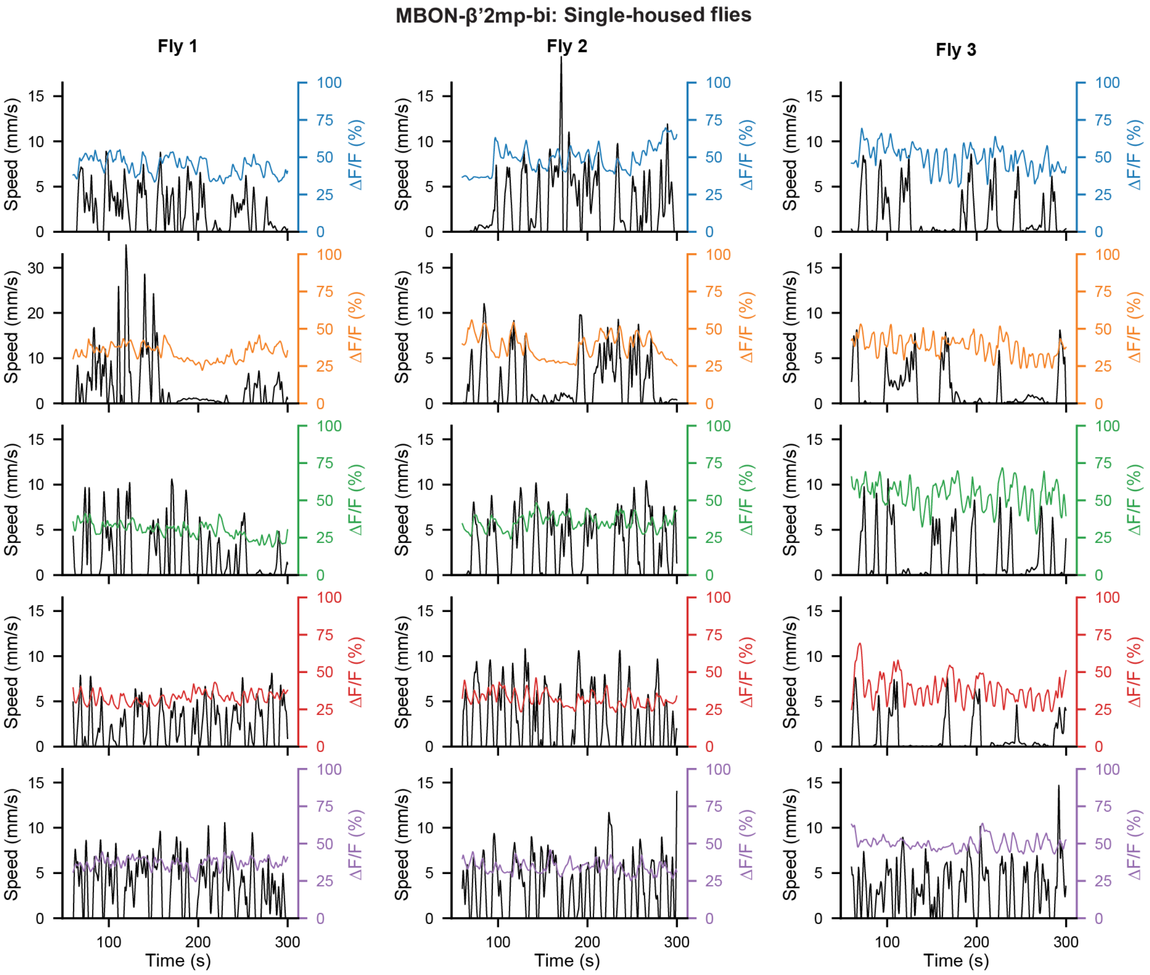


**Supp. File Fig.** 8: **MBON-$\beta$'2mp-bi: Single-housed flies neural activity and spherical treadmill speed.** Treadmill-derived locomotor speed (black) and ΔF/F neural activity traces for each single-housed fly recorded across the five experimental epochs (color-coded as in **Supp. File Fig.** 1).

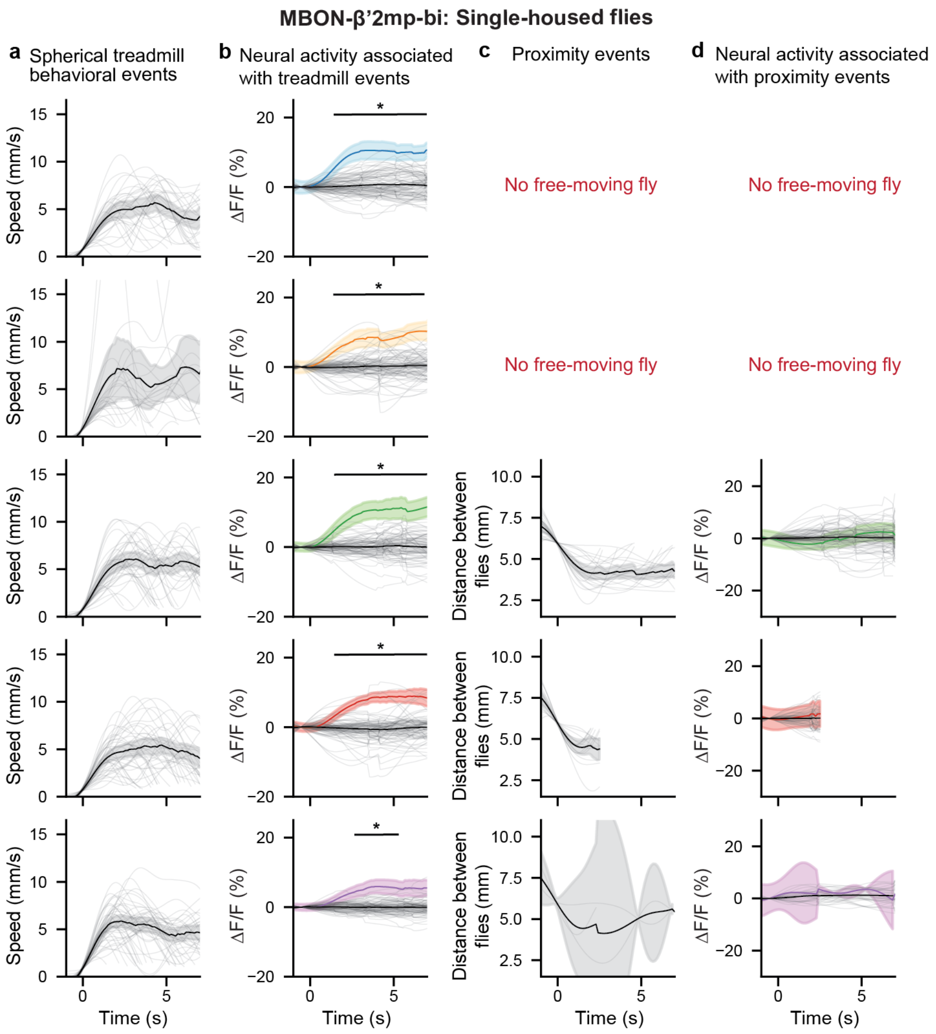


**Supp. File Fig.** 9: **MBON-$\beta$'2mp-bi: Single-housed flies spherical treadmill speed, and proximity events. (a)** Behavioral events were selected for each epoch by thresholding treadmill speed and time-locked to event onset (0 s). Overlaid are individual events (translucent traces) and the mean of all events for all three animals for a given epoch (solid black traces) along with the 95% confidence interval (gray transparency). **(b)** Mean neural activity and 95% confidence interval associated with the spherical treadmill events (traces color-coded by experimental epoch **Supp. File Fig.** 1). Overlaid are traces (translucent gray) representing the mean activity of each set of randomly selected events. The means and confidence intervals of all randomly selected events are also shown for comparison (black traces). Mean traces were compared at each time point using a permutation test followed by a correction for multiple comparisons using the false discovery rate for positively correlated tests. * $P < 0.05$. All other comparisons were statistically similar to one another (i.e., $P \geq 0.05$). **(c)** Behavioral events were selected for each epoch by thresholding the distance between flies and time-locked to event onset (0 s). Overlaid are individual events (translucent traces) and the mean of all events for all three animals for a given epoch (solid black traces) along with the 95% confidence interval (gray transparency). **(d)** Mean neural activity and 95% confidence interval associated with the proximity events (traces color-coded by experimental epoch). Overlaid are traces (translucent gray) representing the mean activity of each set of randomly selected events. The means and confidence intervals of all randomly selected events are also shown for comparison (black traces). Mean traces were compared at each time point using a permutation test followed by a correction for multiple comparisons using the false discovery rate for positively correlated tests. All comparisons were statistically similar to one another (i.e., $P \geq 0.05$).

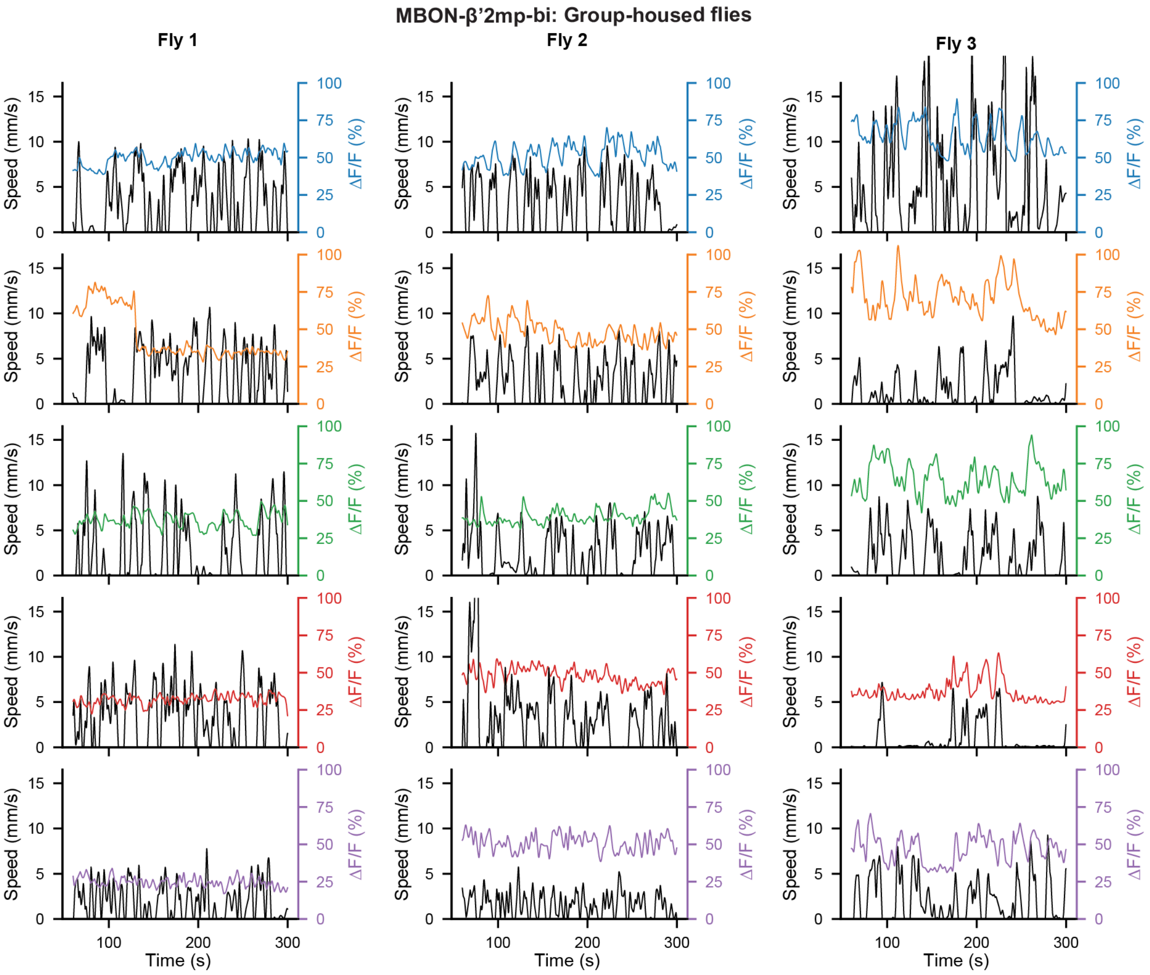


**Supp. File Fig.** 10: **MBON-$\beta$’2mp-bi: Group-housed flies neural activity and spherical treadmill speed.** Treadmill-derived locomotor speed (black) and $\Delta$F/F neural activity traces for each group-housed fly recorded across the five experimental epochs (color-coded as in **Supp. File Fig.** 1).

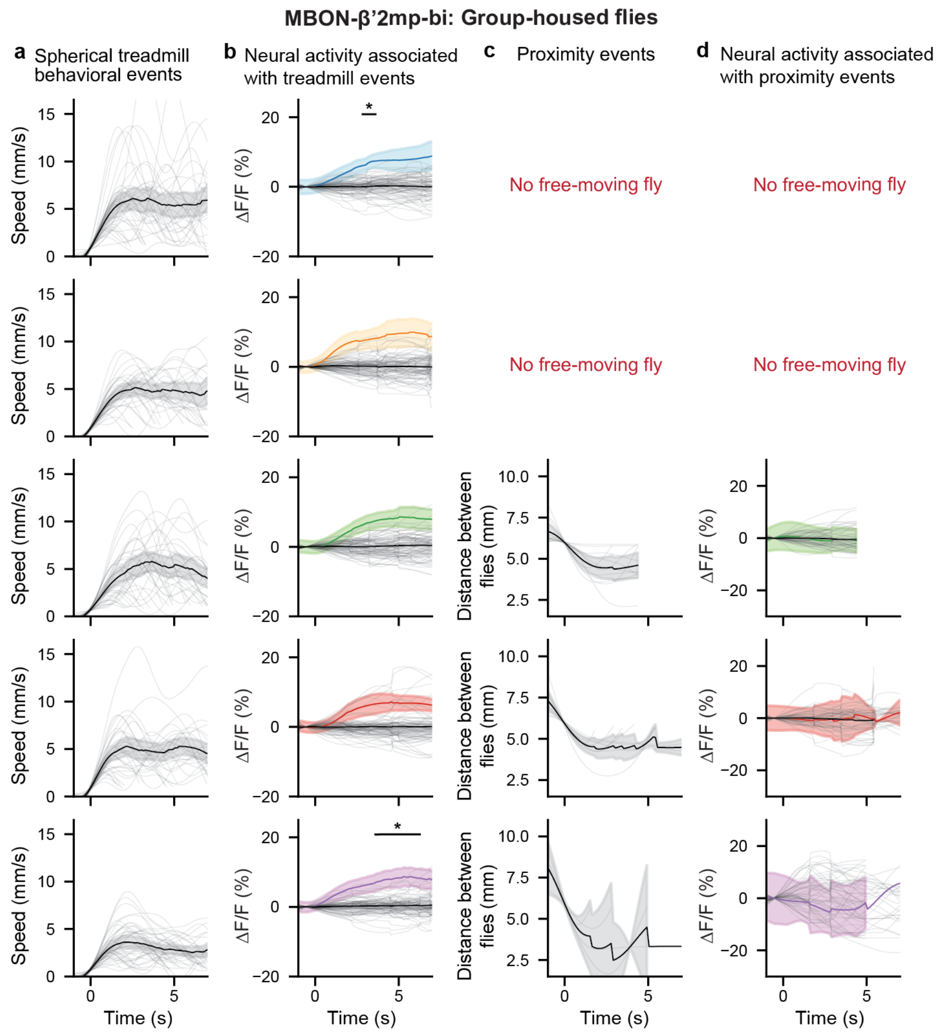


**Supp. File Fig.** 11: **MBON-$\beta$'2mp-bi: Group-housed flies spherical treadmill speed, and proximity events.** **(a)** Behavioral events were selected for each epoch by thresholding treadmill speed and time-locked to event onset (0 s). Overlaid are individual events (translucent traces) and the mean of all events for all three animals for a given epoch (solid black traces) along with the 95% confidence interval (gray transparency). **(b)** Mean neural activity and 95% confidence interval associated with the spherical treadmill events (traces color-coded by experimental epoch **Supp. File Fig.** 1). Overlaid are traces (translucent gray) representing the mean activity of each set of randomly selected events. The means and confidence intervals of all randomly selected events are also shown for comparison (black traces). Mean traces were compared at each time point using a permutation test followed by a correction for multiple comparisons using the false discovery rate for positively correlated tests. * $P < 0.05$. All other comparisons were statistically similar to one another (i.e., $P \geq 0.05$). **(c)** Behavioral events were selected for each epoch by thresholding the distance between flies and time-locked to event onset (0 s). Overlaid are individual events (translucent traces) and the mean of all events for all three animals for a given epoch (solid black traces) along with the 95% confidence interval (gray transparency). **(d)** Mean neural activity and 95% confidence interval associated with the proximity events (traces color-coded by experimental epoch). Overlaid are traces (translucent gray) representing the mean activity of each set of randomly selected events. The means and confidence intervals of all randomly selected events are also shown for comparison (black traces). Mean traces were compared at each time point using a permutation test followed by a correction for multiple comparisons using the false discovery rate for positively correlated tests. All comparisons were statistically similar to one another (i.e., $P \geq 0.05$).

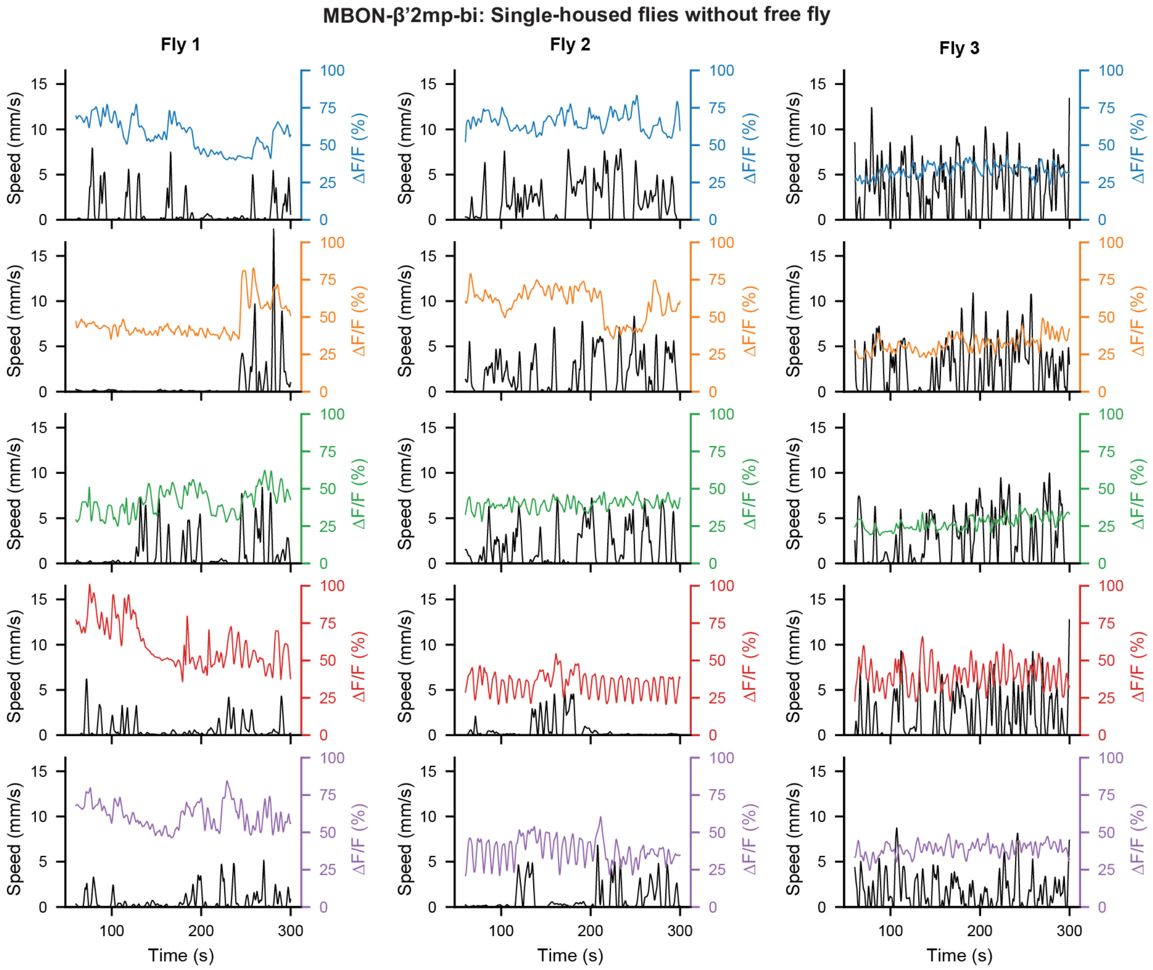


**Supp. File Fig.** 12: **MBON-$\beta$’2mp-bi: Single-housed flies neural activity and spherical treadmill speed without a freely-moving fly in the arena.** Treadmill-derived locomotor speed (black) and ΔF/F neural activity traces for each single-housed fly recorded across the five experimental epochs (color-coded as in **Supp. File Fig.** 1). There was no free-moving fly during the experiment.

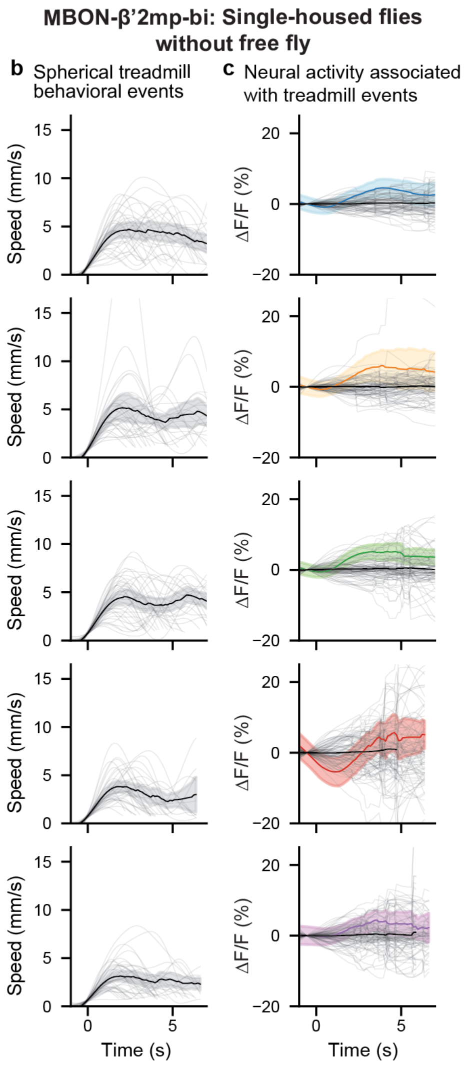


**Supp. File Fig.** 13: **MBON-$\beta$'2mp-bi: Single-housed flies spherical treadmill speed, and proximity events without a freely moving fly in the arena. (a)** Behavioral events were selected for each epoch by thresholding treadmill speed and time-locked to event onset (0 s). Overlaid are individual events (translucent traces) and the mean of all events for all three animals for a given epoch (solid black traces) along with the 95% confidence interval (gray transparency). **(b)** Mean neural activity and 95% confidence interval associated with the spherical treadmill events (traces color-coded by experimental epoch **Supp. File Fig.** 1). Overlaid are traces (translucent gray) representing the mean activity of each set of randomly selected events. The means and confidence intervals of all randomly selected events are also shown for comparison (black traces). Mean traces were compared at each time point using a permutation test followed by a correction for multiple comparisons using the false discovery rate for positively correlated tests. All comparisons were statistically similar to one another (i.e., $P \geq 0.05$).

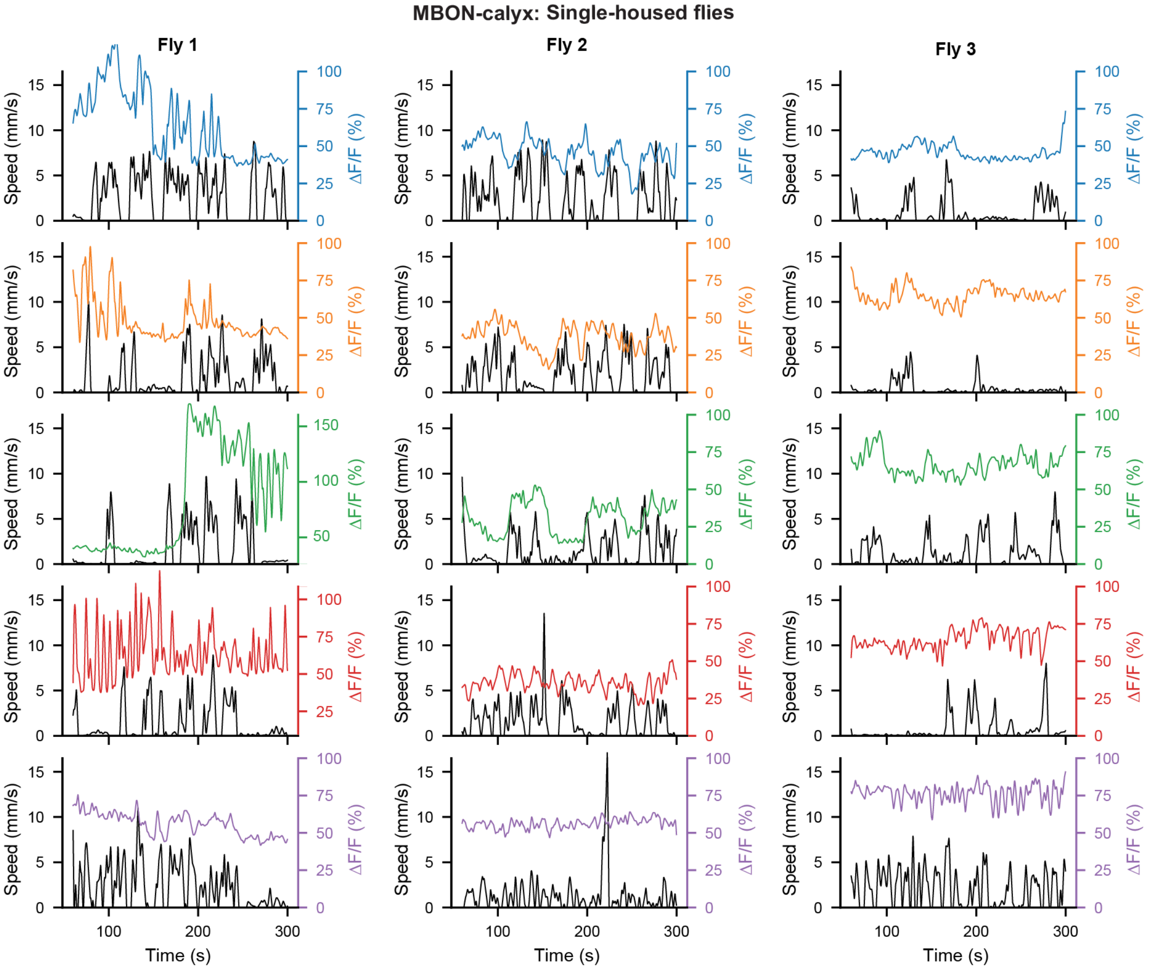


**Supp. File Fig.** 14: **MBON-calyx: Single-housed flies neural activity, spherical treadmill speed, and proximity events.** Treadmill-derived locomotor speed (black) and $\Delta$F/F neural activity traces for each single-housed fly recorded across the five experimental epochs (color-coded as in **Supp. File Fig.** 1).

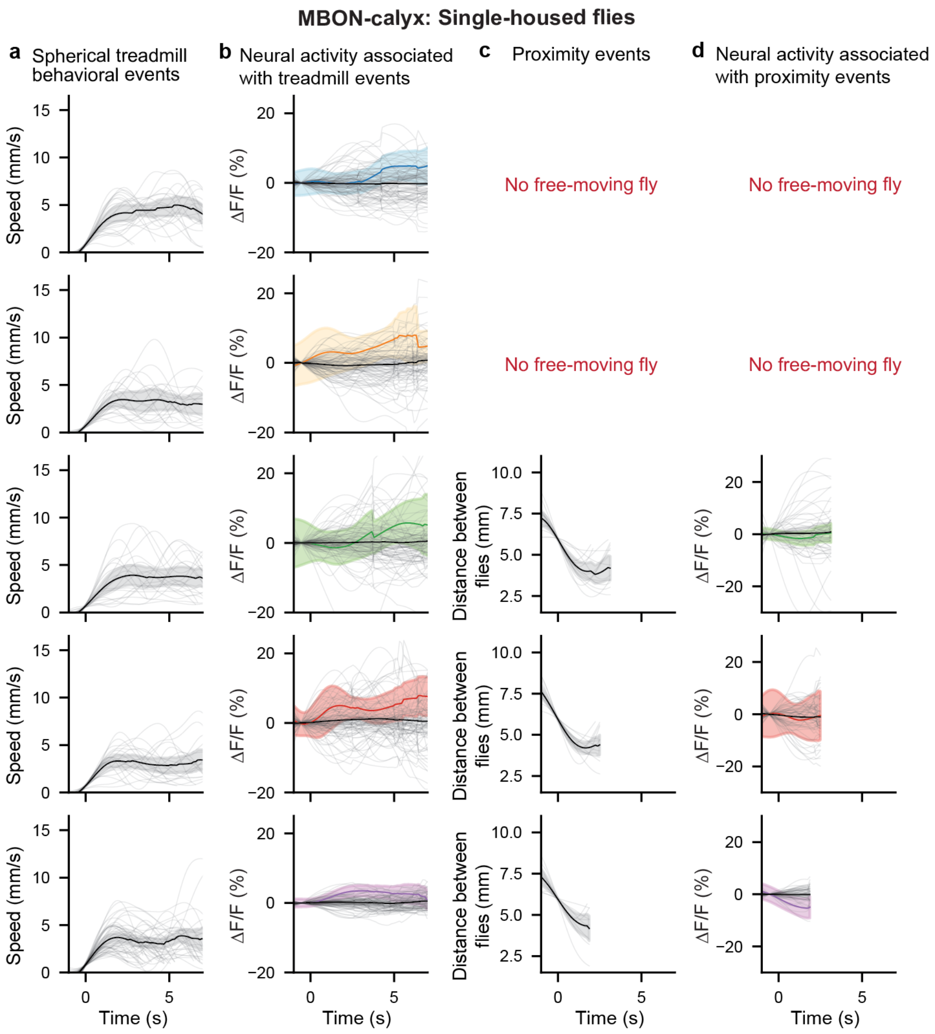


**Supp. File Fig.** 15: **MBON-calyx: Single-housed flies spherical treadmill speed and proximity events.** **(a)** Behavioral events were selected for each epoch by thresholding treadmill speed and time-locked to event onset (0 s). Overlaid are individual events (translucent traces) and the mean of all events for all three animals for a given epoch (solid black traces) along with the 95% confidence interval (gray transparency). **(b)** Mean neural activity and 95% confidence interval associated with the spherical treadmill events (traces color-coded by experimental epoch **Supp. File Fig.** 1). Overlaid are traces (translucent gray) representing the mean activity of each set of randomly selected events. The means and confidence intervals of all randomly selected events are also shown for comparison (black traces). Mean traces were compared at each time point using a permutation test followed by a correction for multiple comparisons using the false discovery rate for positively correlated tests. All comparisons were statistically similar to one another (i.e., $P \geq 0.05$). **(c)** Behavioral events were selected for each epoch by thresholding the distance between flies and time-locked to event onset (0 s). Overlaid are individual events (translucent traces) and the mean of all events for all three animals for a given epoch (solid black traces) along with the 95% confidence interval (gray transparency). **(d)** Mean neural activity and 95% confidence interval associated with the proximity events (traces color-coded by experimental epoch). Overlaid are traces (translucent gray) representing the mean activity of each set of randomly selected events. The means and confidence intervals of all randomly selected events are also shown for comparison (black traces). Mean traces were compared at each time point using a permutation test followed by a correction for multiple comparisons using the false discovery rate for positively correlated tests. All comparisons were statistically similar to one another (i.e., $P \geq 0.05$).

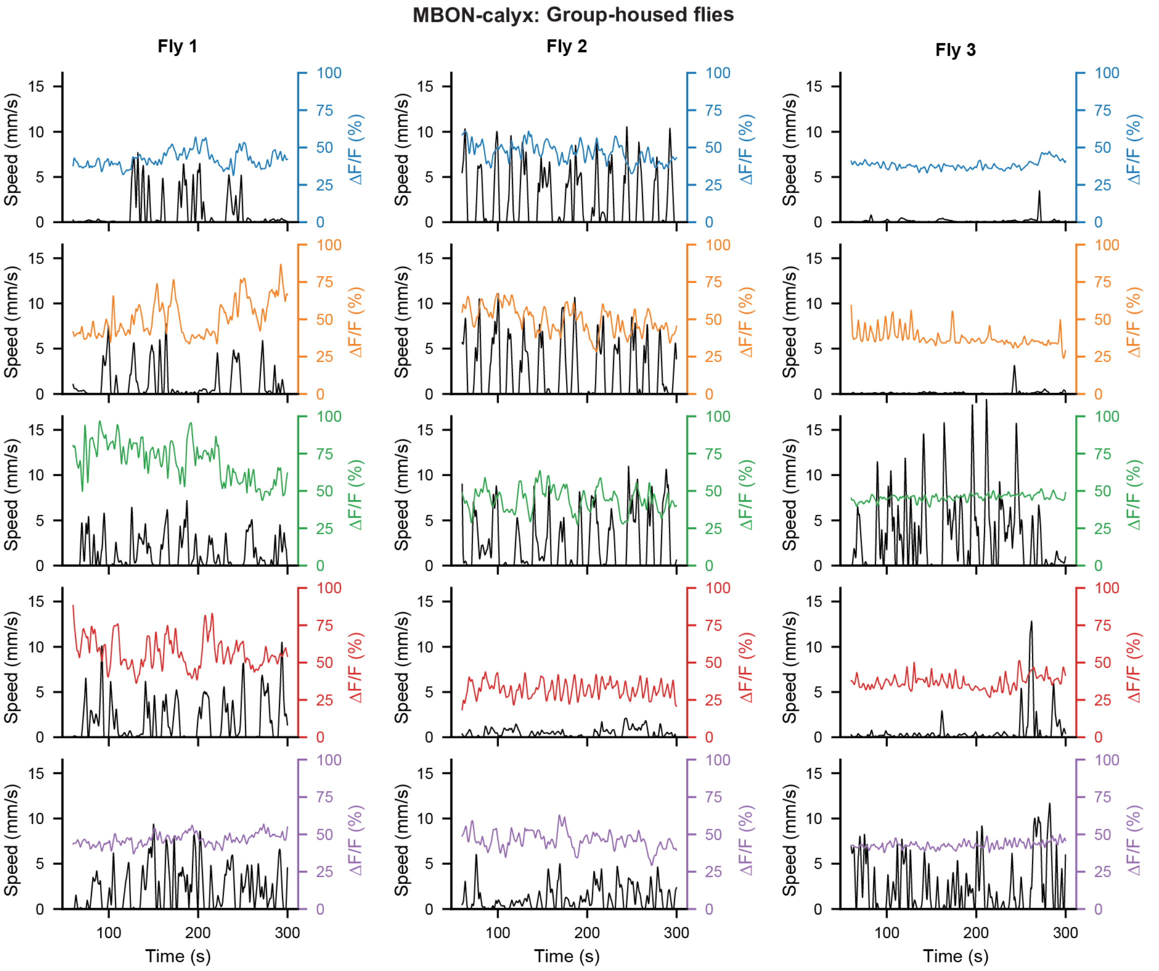


**Supp. File Fig.** 16: **MBON-calyx: Group-housed flies neural activity and spherical treadmill speed.** Treadmill-derived locomotor speed (black) and ΔF/F neural activity traces for each group-housed fly recorded across the five experimental epochs (color-coded as in **Supp. File Fig.** 1).

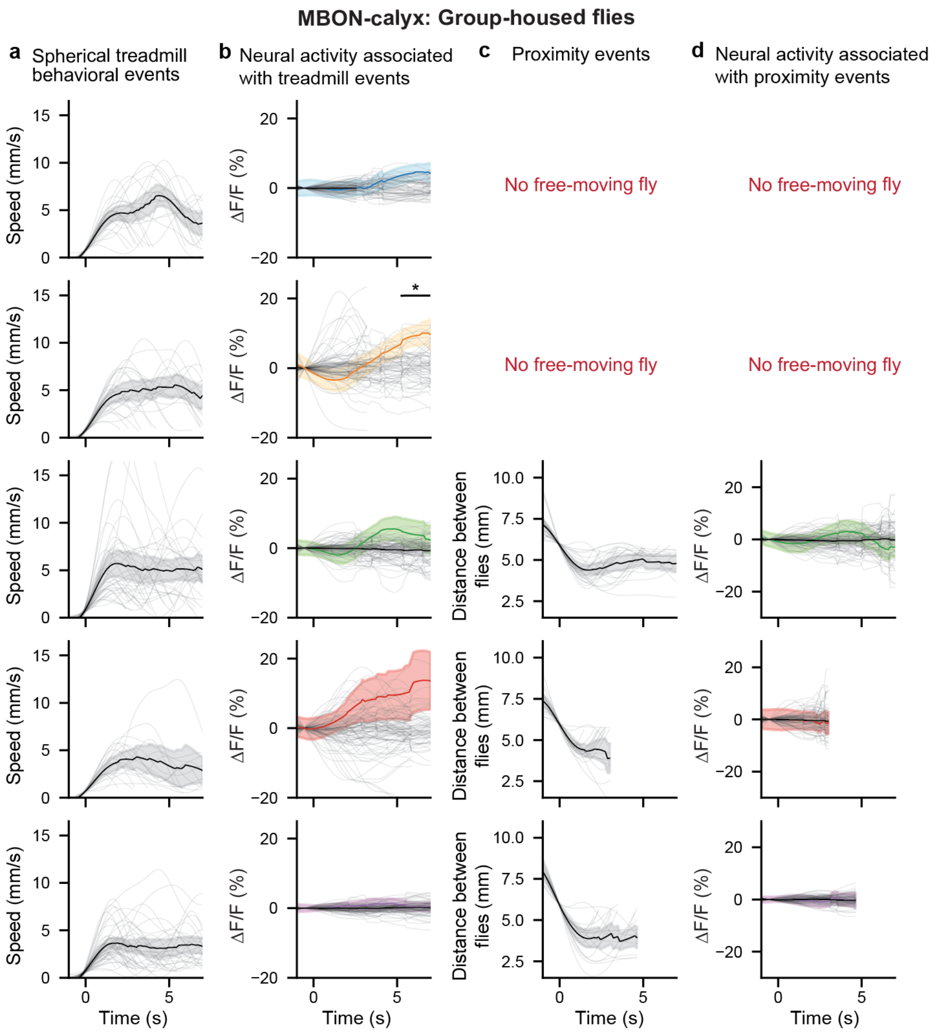


**Supp. File Fig.** 17: **MBON-calyx: Group-housed flies spherical treadmill speed and proximity events. (a)** Behavioral events were selected for each epoch by thresholding treadmill speed and time-locked to event onset (0 s). Overlaid are individual events (translucent traces) and the mean of all events for all three animals for a given epoch (solid black traces) along with the 95% confidence interval (gray transparency). **(b)** Mean neural activity and 95% confidence interval associated with the spherical treadmill events (traces color-coded by experimental epoch **Supp. File Fig.** 1). Overlaid are traces (translucent gray) representing the mean activity of each set of randomly selected events. The means and confidence intervals of all randomly selected events are also shown for comparison (black traces). Mean traces were compared at each time point using a permutation test followed by a correction for multiple comparisons using the false discovery rate for positively correlated tests. * $P < 0.05$. All other comparisons were statistically similar to one another (i.e., $P \geq 0.05$). **(c)** Behavioral events were selected for each epoch by thresholding the distance between flies and time-locked to event onset (0 s). Overlaid are individual events (translucent traces) and the mean of all events for all three animals for a given epoch (solid black traces) along with the 95% confidence interval (gray transparency). **(d)** Mean neural activity and 95% confidence interval associated with the proximity events (traces color-coded by experimental epoch). Overlaid are traces (translucent gray) representing the mean activity of each set of randomly selected events. The means and confidence intervals of all randomly selected events are also shown for comparison (black traces). Mean traces were compared at each time point using a permutation test followed by a correction for multiple comparisons using the false discovery rate for positively correlated tests. All comparisons were statistically similar to one another (i.e., $P \geq 0.05$).

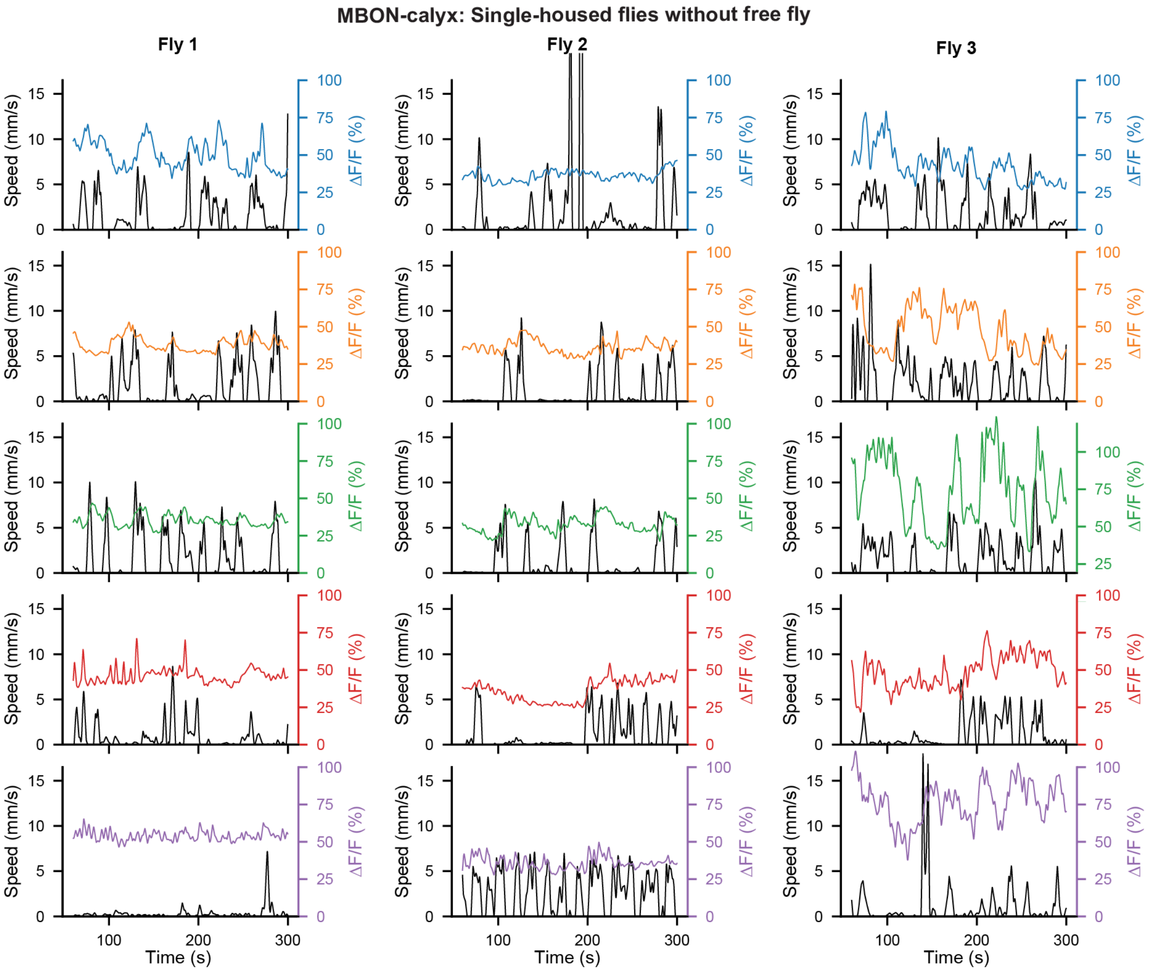


**Supp. File Fig.** 18: **MBON-calyx: Single-housed flies neural activity and spherical treadmill speed without a freely-moving fly in the arena.** Treadmill-derived locomotor speed (black) and ΔF/F neural activity traces for a single-housed fly across the five experimental epochs (color-coded as in **Supp. File Fig.** 1). There was no free-moving fly during the experiment.

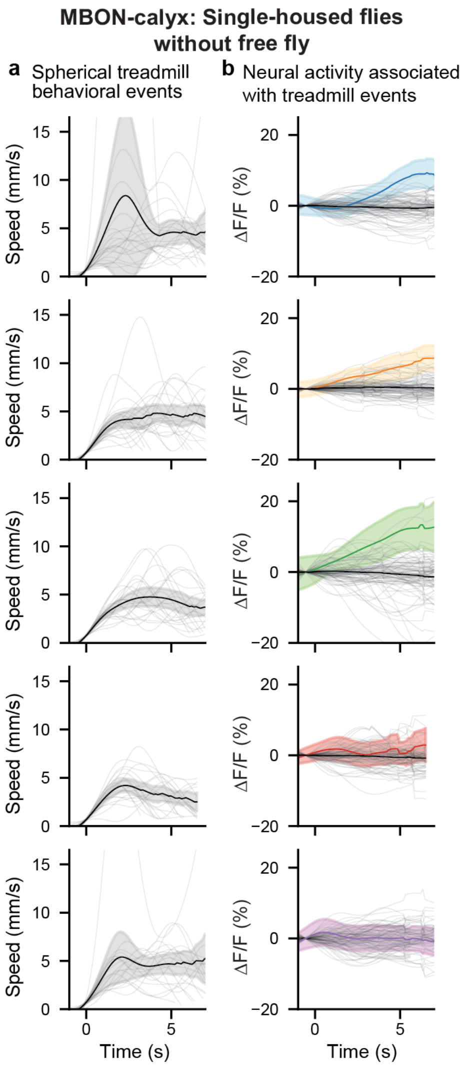


**Supp. File Fig.** 19: **MBON-calyx: Single-housed flies spherical treadmill speed, and proximity events without a freely-moving fly in the arena.** **(a)** Behavioral events were selected for each epoch by thresholding treadmill speed and time-locked to event onset (0 s). Overlaid are individual events (translucent traces) and the mean of all events for all three animals for a given epoch (solid black traces) along with the 95% confidence interval (gray transparency). **(b)** Mean neural activity and 95% confidence interval associated with the spherical treadmill events (traces color-coded by experimental epoch **Supp. File Fig.** 1). Overlaid are traces (translucent gray) representing the mean activity of each set of randomly selected events. The means and confidence intervals of all randomly selected events are also shown for comparison (black traces). Mean traces were compared at each time point using a permutation test followed by a correction for multiple comparisons using the false discovery rate for positively correlated tests. All comparisons were statistically similar to one another (i.e., $P \geq 0.05$).
